# Comparing the effect of imputation reference panel composition in four distinct Latin American cohorts

**DOI:** 10.1101/2024.04.11.589057

**Authors:** Jennifer N French, Victor Borda Pua, Roland Laboulaye, Thiago Peixoto Leal, Mario Cornejo Olivas, Maria Fernanda Lima-Costa, Bernardo L Horta, Mauricio L Barreto, Eduardo Tarazona-Santos, Ignacio Mata, Timothy D. O’Connor

## Abstract

Genome-wide association studies have been useful in identifying genetic risk factors for various phenotypes. These studies rely on imputation and many existing panels are largely composed of individuals of European ancestry, resulting in lower levels of imputation quality in underrepresented populations. We aim to analyze how the composition of imputation reference panels affects imputation quality in four target Latin American cohorts. We compared imputation quality for chromosomes 7 and X when altering the imputation reference panel by: 1) increasing the number of Latin American individuals; 2) excluding either Latin American, African, or European individuals, or 3) increasing the Indigenous American (IA) admixture proportions of included Latin Americans. We found that increasing the number of Latin Americans in the reference panel improved imputation quality in the four populations; however, there were differences between chromosomes 7 and X in some cohorts. Excluding Latin Americans from analysis resulted in worse imputation quality in every cohort, while differential effects were seen when excluding Europeans and Africans between and within cohorts and between chromosomes 7 and X. Finally, increasing IA-like admixture proportions in the reference panel increased imputation quality at different levels in different populations. The difference in results between populations and chromosomes suggests that existing and future reference panels containing Latin American individuals are likely to perform differently in different Latin American populations.

## Background

Genome-wide association studies (GWAS) have been useful in identifying genetic risk factors for various phenotypes. However, these analyses often include mostly individuals of European ancestry, leaving other populations vastly understudied and underrepresented in research, representing less than 6% of all GWAS participants ^[1,2]^. Furthermore, GWAS typically use genotype panels and rely on imputation to include non-genotyped SNPs. Imputation increases coverage of the genome, allowing for analysis of non-genotyped SNPs with the disease or trait of interest. Many existing imputation reference panels are mostly based on individuals of European ancestry. Imputation relies on linkage disequilibrium (LD) patterns, which vary between populations ^[3]^. Therefore, low diversity in imputation reference panels results in a lack of LD patterns, making it difficult to reach the same level of imputation quality in non-European populations as in European populations. Latin Americans are one of the fastest growing minorities in the United States ^[4]^ but represent only 0.64% of all GWAS participants ^[1,2]^.

Imputation is especially challenging in Latin American individuals due to the lack of Indigenous American data available and the high degree of admixture between Europeans, Africans, and Indigenous Americans in Latin American individuals ^[5,6]^.

Another issue arises with the exclusion of chromosome X from most GWAS studies due to the complexity of analysis. Sex is associated with many phenotypes and excluding the X chromosome may leave the true genetic risk for phenotypes underestimated. Latin American populations resulted from a recent sex-biased admixture process involving continental ancestry. Historical events, like colonization, have led to differing ancestry proportions between the autosomes and the sex chromosomes ^[7]^, and different ancestry background in men and women, specifically with higher European ancestry in males than females. Over time, European ancestry is likely to be lower on the X chromosome than the autosomes ^[7–10]^.

Imputation studies have compared existing panels containing some diverse populations, including the Haplotype Reference Consortium (HRC), 1000 Genomes Project (1KGP), the Consortium on Asthma among African-ancestry Populations in the Americas (CAAPA), and Trans-Omics for Precision Medicine (TOPMed) panels ^[11–14]^. The HRC contains haplotypes for 64,976 individuals, most of which are of European ancestry ^[12]^. The 1KG panel contains 2,504 individuals from 26 populations from Europe, East Asian, sub-Saharan African, and the Americas ^[11]^. Individuals from 1KG are included in the HRC panel. CAAPA is composed of 883 individuals with African Ancestry from the Americas ^[13]^. Finally, TOPMed freeze 9 contains around 158,000 individuals from more than 80 studies and includes diverse populations, ^[14,15]^. A subset of individuals is available in the TOPMed imputation reference panel (n=97,256, including 17,085 Latin Americans) ^[14]^.

The addition of diverse samples to existing panels appears to improve imputation quality of reference panels in a variety of studies ^[11,14,16,17]^. Studies have found that adding whole genome sequencing or genotyping data of Latin American individuals resulted in increased number and quality of imputed SNPs ^[18,19]^. Previous studies have suggested that roughly 3,000 individuals with Indigenous American (IA) ancestry are needed in the panel to match the quality of variants in European ancestry tracts ^[18]^. Another study found that the combination of 500 study participants with HRC reference panel increased coverage by 9% compared to using the HRC alone but did not affect imputation quality ^[17]^. When adding WGS of 1,171 unrelated individuals from São Paulo, Brazil to 1KGP to create a reference panel, imputation results improved in a Brazilian cohort, with more well-imputed SNPs, more rare SNPs, and improved imputation quality ^[20]^. Going even further, one group created an entirely new imputation panel of 2,269 unrelated individuals of Sub-Saharan African ancestry ^[13]^. The new panel, DV-GLx AFAM, outperformed HRC, 1KGP, and CAAPA panels when conducting imputation in African Americans. However, DV-GLx didn’t outperform TOPMed, likely due to having a smaller sample size ^[13]^. While much research has gone into how to improve imputation quality in diverse populations by adding diverse samples to existing panels, little has been done on how the composition of the panel affects imputation quality, and how this may differ between the autosomes and the X chromosome.

To address this gap, we aim to analyze how the composition of the reference panel affects imputation quality in four distinct Latin American target populations (Figure S1) on both the X chromosome and chromosome 7, as it is most similar in size to chromosome X (by number of base pairs). First, we created five reference panels, each with 12,000 samples from TOPMed freeze10b. Each panel contained an increased number of Latin American individuals from 1,000 - 5,000. For the second comparison, we built four additional panels: all populations (Africans, Europeans, Latin Americans), without Africans, without Europeans, and without Latin Americans. We performed imputation and compared imputation quality between panels. Finally, we altered the Latin Americans included in the reference panel by using a sliding window of Latin Americans when ordered by their Indigenous American (IA) admixture proportion. We demonstrate that the composition and number of Latin American individuals included in the imputation reference panel affects imputation quality in Latin American populations and has an impact in genetic epidemiology studies.

## Methods

### Data source for reference panels

We included a subset of 58,646 individuals from the TOPMed Imputation Panel ^[14]^ to create the base reference population for our imputation panels (Table S1). To create the subset, we selected self-reported Latin American individuals included in the Genetics of Latin American Diversity (GLAD) project ^[21]^, as well as individuals from the Women’s Health Initiative, the Jackson Heart Study, the 1KGP, the Human Genome Diversity Project, the Framingham Heart Study, the Barbados Asthma Genetics Study, and the MultiEthnic Study of Atherosclerosis (MESA) ^[11,22–27]^. We calculated the genetic relationship using KING ^[28]^ and excluded related individuals using NAToRA ^[29]^. In this work, we considered individuals with third-degree (kinship coefficient > 0.0442) or higher as related. We also excluded any individuals with missing self-reported race/ethnicity or self-reported race/ethnicity other than White, Black, or Hispanic/Latin American. This resulted in a sample size of 35,310 unrelated individuals: 17,807 self-described Latin Americans; 11,256 self-described Europeans, and 6,247 Africans.

### Minor Allele Count Comparison

We compared reference panels with various minor allele count (MAC) thresholds to visualize the effect of each on imputation results. We created 13 imputation panels containing the 35,310 individuals with self-described race/ethnicity data available, with minor allele count (MAC) cutoffs of: 1, 2, 5, 10, 20, 30, 40, 50, 60, 70, 80, 90, and 100. After applying the MAC cutoff, we also limited the panels to biallelic SNPs on chromosome 21 to save computation time. We then used the 13 reference panels to impute genotype data for the Latin American Research Consortium on the Genetics of Parkinson’s Disease (LARGE-PD) ^[30]^ and compared the R^2^ values of imputed SNPs with each panel. Based on the results from the MAC comparison, we concluded that removing singletons was sufficient for panel creation (Figures S2, S3, & S4).

### Panel creation

#### Increasing number of Latin Americans (NoLA)

To address underrepresentation of Latin Americans in existing panels, we created seven panels for both chromosome X and chromosome 7 to investigate how increasing the number of Latin Americans affects imputation quality. We generated seven NoLA panels with different sample size and proportion between European (EUR), African (AFR), and Latin American (LatAm) individuals: (i) 5,500 EUR, 5,500 AFR, and 1,000 LatAm (NoLA-1); (ii) 5,000 EUR, 5,000 AFR, and 2,000 LatAm (NoLA-2); (iii) 4,500 EUR, 4,500 AFR, and 3,000 LatAm (NoLA-3); (iv) 4,000 EUR, 4,000 AFR, and 4,000 LatAm (NoLA-4); (v) 3,500 EUR, 3,500 AFR and 5,000 LatAm (NoLA-5); (vi) 1,000 EUR, 1,000 AFR, and 1,000 LatAm (NoLA-S), and (vii) 2,500 EUR, 2,500 AFR, and 2,500 LatAm (NoLA-M) (Table S2). We then created new VCF files that included only individuals selected for each of the panels and limited to SNPs with a minor allele count (MAC) > 1 using bcftools v.11.1^[31]^. We also limited the dataset to include only biallelic SNPs, removed SNPs with missingness greater than 10% and SNPs with a Hardy Weinberg Equilibrium (HWE) test p-value < 1 x 10^-6^ using plink 1.9 ^[32]^. We then converted the vcf to m3vcf using minimac3, resulting in a total of 14 panels: 7 panels for each of the two chromosomes of interest.

#### Leave one population out (LOPO)

To address how sex-biased admixture in Latin Americans creates different admixture patterns in the autosomes and sex chromosomes, we randomly selected 6,000 individuals from each of the three population categories (Latin American, European, and African). We created three panels, each excluding one of the three populations. This resulted in panels containing: 1) European and African (LOPO-EA) individuals, 2) European and Latin American (LOPO-EL) individuals, and 3) African and Latin American (LOPO-AL) individuals. We also created a fourth panel that included 4,000 individuals from all three populations (LOPO-all) (Table S3). We created five versions of each LOPO panel by randomly sampling the individuals four additional times. This was to ensure that any findings are not due to the origin of Latin Americans sampled. We created 40 imputation reference panels following the same steps outlined in NoLA methods.

#### Increasing proportion of IA ancestry (PIAA)

Our first step was to run unsupervised admixture analysis using the program ADMIXTURE ^[33]^. First we created a reference for ADMIXTURE analysis using a subset of 1KGP which included individuals from the following subpopulations: 1) Mexican Ancestry in Los Angeles, California (MXL), Peruvian in Lima, Peru (PEL) from superpopulation Admixed Americans (AMR); 2) Utah residents (CEPH) with Northern and Western Ancestry (CEU), Iberian in Spain (IBS), Finnish in Finland (FIN), British in England and Scotland (GBR), and Toscani in Italia (TSI) from superpopulation European (EUR), and 3) Yoruba in Ibadan, Nigeria (YRI), Esan in Nigeria (ESN), Mandinka in The Gambia (MAG), Mende in Sierra Leone (MSL), and Luhya in Webuye, Kenya (LWK) from superpopulation African (AFR) ^[11]^. When limiting to only individuals within 1KG that had genotype data, we found 149 MXL and PEL, which were grouped to represent Latin Americans. In order to keep representation among populations the same, we randomly selected 149 individuals from the EUR and AFR populations to include in the reference panel for ADMIXTURE analysis. We then selected one individual at a time from the 35,310 individuals in the subset of the TOPMed Imputation Panel, running ADMIXTURE 35,310 times. Independent ADMIXTURE analyses were then performed on the 447 selected individuals from 1KGP plus the target TOPMed individual. The output of this analysis is African, European, and Indigenous American -like proportions for each individual. We repeated this until we had ancestry proportions for all 35,310 individuals.

To create the panels, we randomly selected 4,000 European and African individuals to be included in all panels. We then ordered the 17,807 Latin American individuals by IA ancestry proportions and selected individuals for each panel in windows of 4,000, sliding down the rank by 2,000 each time until we reached position 16,000 (Table S4, Figure S5). Then, we selected individuals from position 13,808 - 17,807 (Table S4). Finally, we created a panel with a random selection of 900 individuals from panels PIAA-1, PIAA-3, PIAA-5, PIAA-7, and 400 individuals from position 16,0001-17,807 to represent Latin American individuals with varying IA-like proportions. After combining the Latin American individuals selected for each panel with the randomly selected European and African individuals, we created 18 imputation reference panels following the same steps outlined in NoLA methods.

### Target Populations

We selected four cohorts as target populations: 1) the Latin American Research Consortium on the Genetics of Parkinson’s Disease (LARGE-PD); 2) the Columbia University Study of Caribbean Hispanics with Familial or Sporadic Late Onset Alzheimer’s Disease (CUSCH-LOAD), 3) the Slim Initiative in Genomic Medicine for the Americas (SIGMA), and 4) Genetic Epidemiology of Complex Diseases in Brazilian population-based cohorts (EPIGEN)-Brazil^[30,34–36]^.

LARGE-PD is a Parkinson’s disease cohort composed of 1,504 individuals from Brazil, Chile, Colombia, Peru, and Uruguay with available genotyped data from the Multi-Ethnic Genotyping Array (MEGA) from Illumina ^[30]^. On average, the ancestry proportions in LARGE-PD are 5.9% African, 54.1% European, and 40.0% Indigenous American (Table S5, Figure S6). CUSCH-LOAD contains 3,967 individuals mostly from Puerto Rico and the Dominican Republic with genotyped data on the Illumina Omni 1M chip ^[35]^, and average ancestry proportions of 33.7% African, 57.4% European, and 8.9% Indigenous American. We analyzed this cohort as a whole, as well as split into two cohorts based on country of origin. Average ancestry proportions differ between individuals from the Dominican Republic (36.8% African, 55.7% European, 7.6% Indigenous American) and Puerto Rico (17.6% African, 68.8% European, 13.5% Indigenous American). SIGMA is a Mexican cohort of 8,214 individuals focused on type 2 diabetes with available genotyped data from the Illumina Omni 2.5 array ^[36]^. Average ancestry proportions in SIGMA are 2.1% African, 27.0% European, and 70.9% Indigenous American. Finally, EPIGEN-Brazil is a Brazilian genetic study of 6,487 individuals with available genotyped data from the Illumina Omni 2.5 array ^[34]^. We split the EPIGEN-Brazil cohort into the three cohorts (Salvador^[37]^, Bambuí^[38]^, and Pelotas^[39]^). Individuals from each region have different average genetic ancestry proportions (Bambuí: 16% African, 76.3% European, 7.4% Indigenous American; Pelotas: 14.9% African, 77.3% European, 7.9% Indigenous American, and Salvador: 50.0% African, 43.5% European, and 6.5% Indigenous American).

### Quality control for target populations

Autosomal data for each target population were split into individual files by chromosome number and lifted to human genome build 38 via UCSC LiftOver ^[40]^. After lifting, the autosomes were recombined into one file per target population. For CUSCH-LOAD, we also created two additional plink files containing only individuals from Puerto Rico (PR) or the Dominican Republic (DR). We then ran relatedness via KING ^[28]^ to get kinship coefficients for each pair of individuals in each target population, and individuals with a third degree or closer relationship were removed using NAToRA ^[29]^.

The X chromosome was limited to the non-pseudoautosomal region. We followed data cleaning steps outlined in XWAS ^[41,42]^, which includes removing any individuals failing a sex check (on LD pruned data), and SNPs with different missingness or MAF between males and females. Finally, we filtered SNPs to exclude those with a MAC ≤ 1, missingness greater than 5%, and SNPs with a HWE test p-value < 1×10^-6^ using plink 2.0 ^[32]^.

For chromosome 7, we included only individuals included in the X chromosome data and also excluded SNPs with a MAC <= 1, missingness > 5%, and HWE p-value < 1 x 10^-6^ via plink 2.0.

### Imputation

Imputation was completed using all reference panels in all target populations using minimac4, totaling 576 imputations in all populations. Minimac4 outputs R^2^ value for every SNP and an empirical R^2^ value for every genotyped SNP. Empirical R^2^ (EmpR^2^) is a measure of the squared correlation between the true genotype and imputed dosage at each genotyped SNP. This is done via masking the true genotype and comparing it to the imputed dosage.

### Analysis

#### EmpR^2^ comparison

We found the overlap of genotyped SNPs included in imputation results using each of the imputation reference panels by target population and chromosome. We combined these EmpR^2^ values for the overlapped SNPs into one file. We then used ggplot2 in R to create a boxplot, violin plot, and Empirical Cumulative Distribution Function plot of EmpR^2^ values in each panel grouped by population. We used the Wilcoxon-signed rank test in R to analyze differences between panels in each population and on each chromosome separately. We account for multiple testing using a Bonferroni correction ^[43]^.

#### R^2^ categories

We placed imputed SNPs into categories based on having an R^2^ of < 0.2; 0.2-0.4; 0.4-0.6; 0.6-0.8, or ≥ 0.8. Using R^[44]^, we created a stacked bar plot of the number and proportion of SNPs in each category based on population and chromosome.

#### Minor allele frequency

We also placed imputed SNPs into categories based on having a MAF of: < 0.0001; 0.0001 - 0.01; 0.1-0.05, or ≥ 0.05. We created a stacked bar plot of the number and proportion of SNPs in each MAF category based on population and chromosome using R^[44]^.

#### R^2^ by MAF bin

We placed each SNP into bins based on having a MAF of: < 0.0001; 0.001-0.005; 0.005-0.01; 0.01-0.05, or ≥ 0.05t. We then used R^[44]^ to create a plot of the mean R^2^ by MAF bin.

## Results

### Increasing number of Latin Americans (NoLA)

With the limited availability of Latin American individuals, it is important to compare imputation quality with increasing numbers of Latin Americans in the imputation reference panel to see where the ratio of Latin Americans to individuals from other populations performs the best between cohorts and chromosomes. In all populations and on both chromosome 7 and X, the Number of Latin American - Small (NoLA-S) panel containing 1,000 African, European, and Latin American individuals each performed worse than Panel NoLA-4 (containing 4,000 individuals from each population) (Figures 1 & S7, Tables S6 & S7). Similarly, the NoLA-M panel (2,500 individuals from each population) resulted in significantly lower empirical R^2^ values than Panel NoLA-4 (Bonferonni corrected p-values < 4.93 x 10^-14^ ), except for chromosome 7 in the Puerto Rican target population, where no difference was found (p-value = 0.866). The observed differences between panels NoLA-S, NoLA-M, and NoLA-4 could be due to either total sample size in the reference panel or the number of Latin American individuals included.

**Figure 1.**
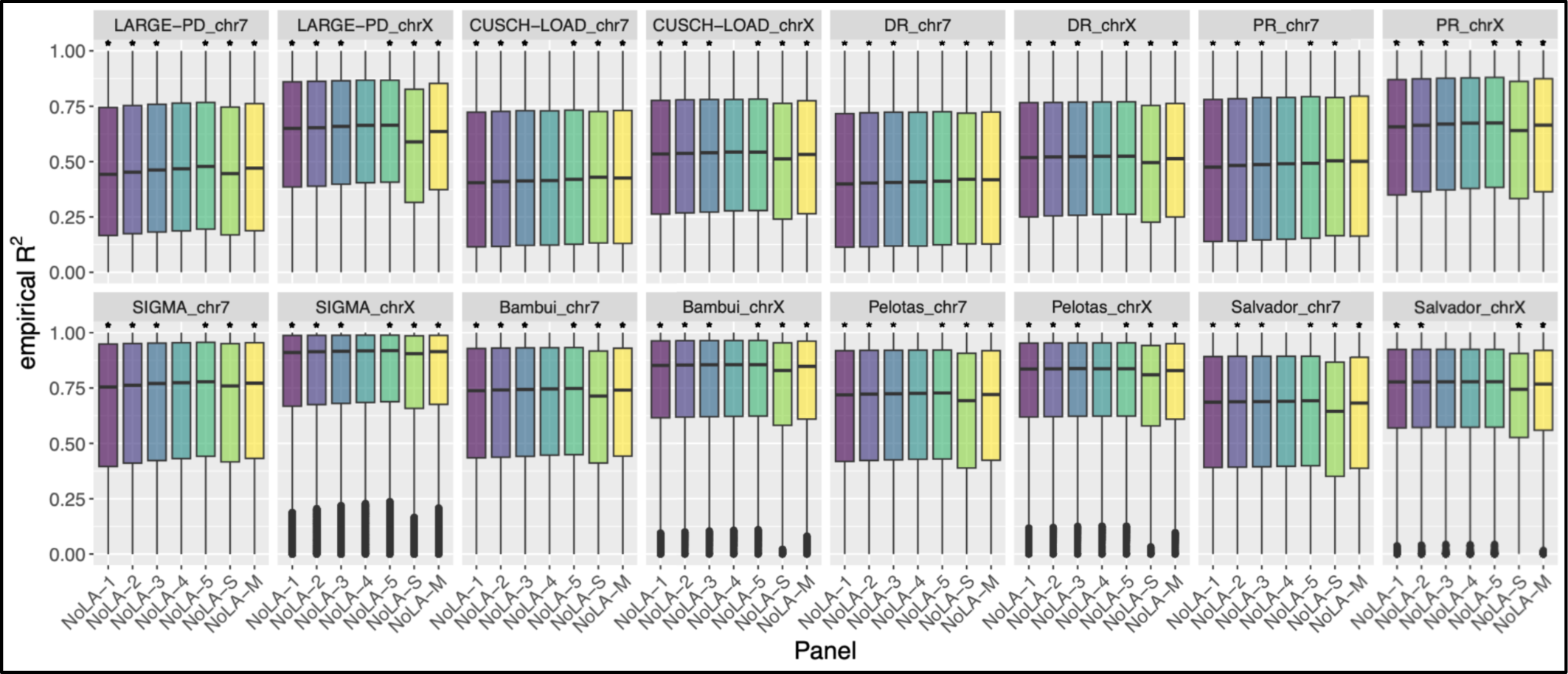
Comparison of empirical R^2^ values for genotyped SNPs. Empirical R^2^ values for each population, using imputation reference panels containing varying numbers of Latin American individuals (NoLA). Pairwise comparisons done with NoLA-4 as a reference using a paired, two-sided Wilcoxon signed rank test. *: p-value < 0.05. The number of Latin Americans increase from panel NoLA-1 to NoLA-5. NoLA-S, NoLA-M, and NoLA-4 contain equal proportions of African, European, and Latin American individuals, with increasing total sample size. LARGE-PD: Latin American Research Consortium on the Genetics of Parkinson’s Disease, CUSCH-LOAD: Columbia University study of Caribbean Hispanics with familial or sporadic late-onset Alzheimer’s disease, DR: subset of CUSCH-LOAD individuals from the Dominican Republic, PR: subset of CUSCH-LOAD individuals from Puerto Rico, SIGMA: the Slim Initiative in genomic medicine for the Americas.

To better isolate the difference due to the number of Latin Americans included in the panel, we kept the sample size the same and altered the number of Latin Americans included in each reference panel. We observed an improvement in imputation quality when more Latin Americans were included in the reference panel. Panels NoLA-1 - NoLA-3 resulted in lower differences in empR2 values on chromosome 7 for all populations (p-values < 0.0143). Panels NoLA-3, NoLA-4, and NoLA-5 were similar for chromosome X in the Salvador cohort of EPIGEN-Brazil (p-values: > 0.99 & 0.879, respectively), while panel NoLA-5 outperformed panel NoLA-4 on chr 7 in the same cohort (p-value =0.00107).

There was a larger median of differences seen on chromosome 7 than X in LARGE-PD, Bambuí, and Pelotas, while the opposite was true for SIGMA and CUSCH-LOAD, both as a whole and as subpopulations (Tables S6 & S7), suggesting that increasing the number of Latin Americans included may affect the autosomes and sex chromosomes differently based on population structure.

We noted similar proportions of SNPs in each MAF category and well imputed SNPs (R^2^ >= 0.8) in each population on each chromosome for panels NoLA-1 - NoLA-5 (Figures S8). Panel NoLA-S resulted in higher proportions of well imputed SNPs, but a much lower number of total SNPs imputed (Figure S9.) In LARGE-PD and SIGMA, panels NoLA-1 - NoLA-4 had noticeably lower mean R^2^ values in imputed SNPs with a MAF of 0.005 - 0.01 (Figure 2). In all populations, imputation on chr X resulted in higher median empirical R^2^ values than on chromosome 7 (Figure 1).

**Figure 2.**
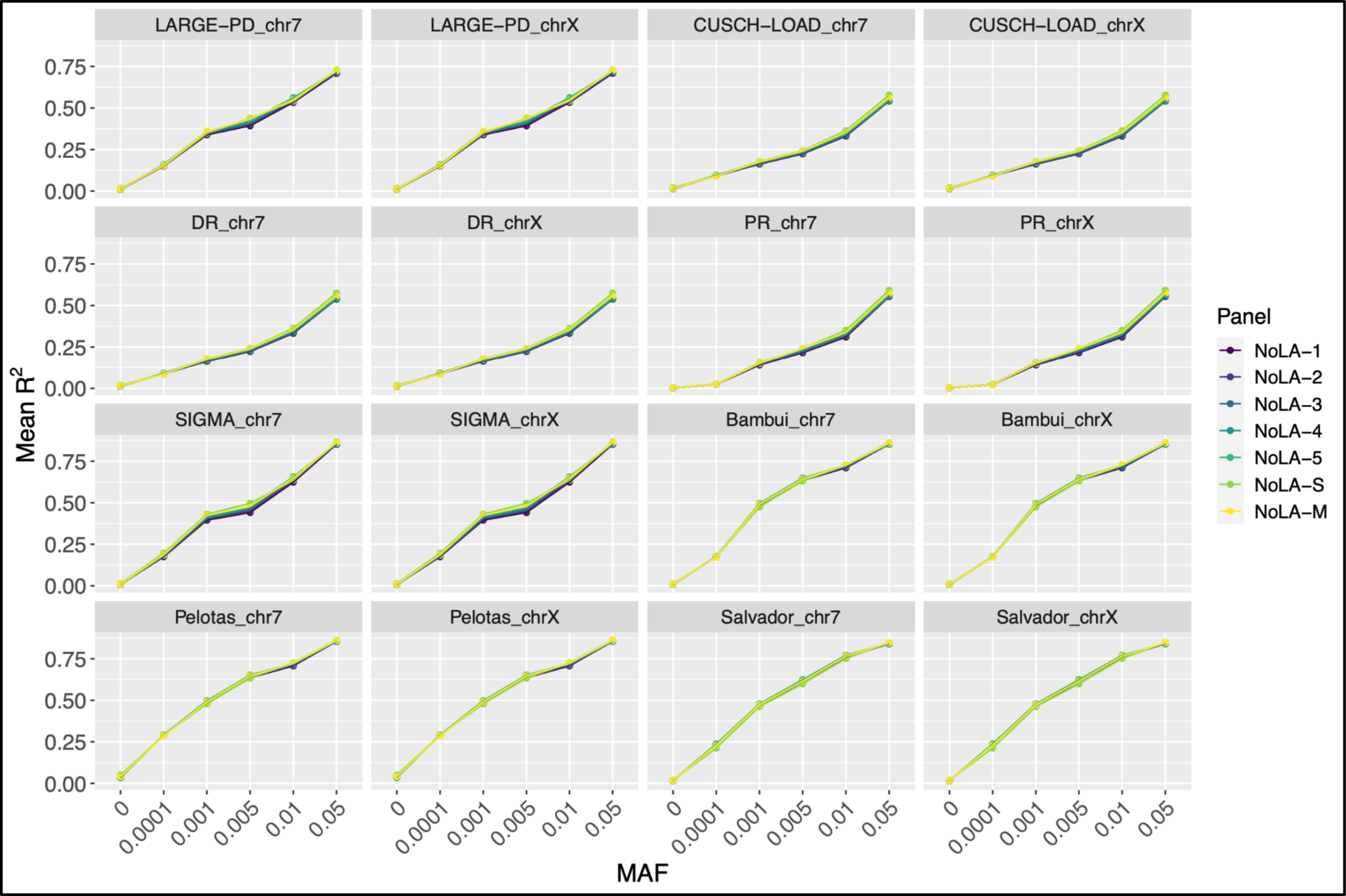
Mean R^2^ of imputed SNPs by MAF bin for NoLA reference panels. The mean R^2^ of imputed SNPs in each MAF bin for each reference panel in each population. Colors correspond to imputation reference panels including Latin American individuals with increasing numbers of Latin American individuals (NoLA). The number of Latin Americans increase from panel NoLA-1 to NoLA-5. NoLA-S, NoLA-M, and NoLA-4 contain equal proportions of African, European, and Latin American individuals, with increasing total sample size. LARGE-PD: Latin American Research Consortium on the Genetics of Parkinson’s Disease, CUSCH-LOAD: Columbia University study of Caribbean Hispanics with familial or sporadic late-onset Alzheimer’s disease, DR: subset of CUSCH-LOAD individuals from the Dominican Republic, PR: subset of CUSCH-LOAD individuals from Puerto Rico, SIGMA: the Slim Initiative in genomic medicine for the Americas.

### Leave one population out (LOPO)

The exclusion of one superpopulation may have different effects on imputation quality in different cohorts or different chromosomes within the same cohort. Excluding Latin Americans individuals in the reference panel resulted in significantly worse imputation quality than the panel containing individuals from Europe, Africa, and Latin America in every target cohort for both chromosomes 7 and X (Figures 3 & S10, Tables S8 & S9).

**Figure 3.**
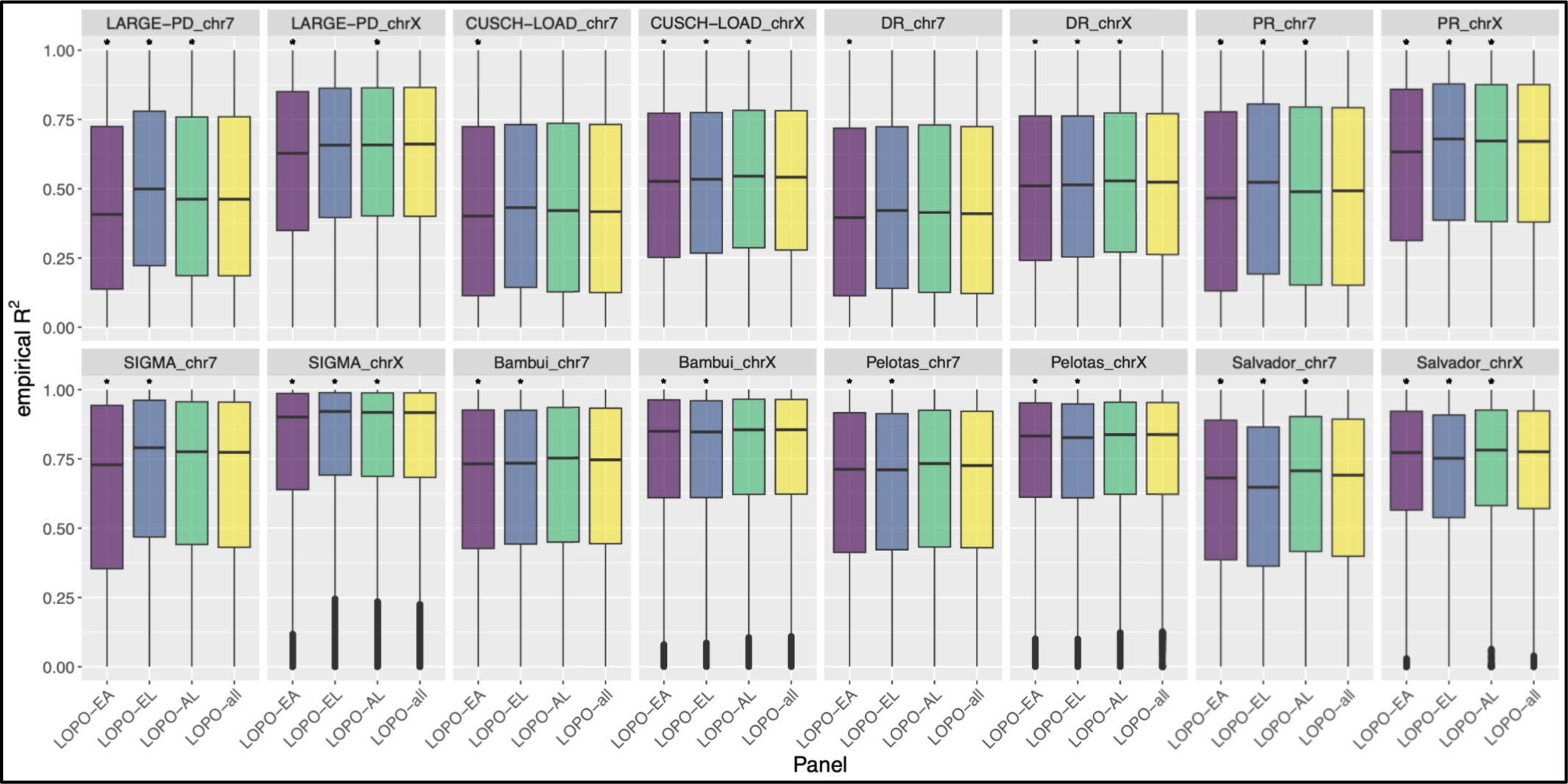
Comparison of empirical R^2^ values for genotyped SNPs. Empirical R^2^ values for each population, when leaving one population out (LOPO) of the imputation reference panels. Pairwise comparisons done with panel 4 as a reference using a paired, two-sided Wilcoxon signed rank test. *: p-value < 0.05. LOPO-EA: Europeans and Africans included in reference panel, LOPO-EL: Europeans and Latin Americans included, LOPO-AL: Africans and Latin Americans included, LOPO-all: Africans, Europeans, and Latin Americans included, LARGE-PD: Latin American Research Consortium on the Genetics of Parkinson’s Disease, CUSCH-LOAD: Columbia University study of Caribbean Hispanics with familial or sporadic late-onset Alzheimer’s disease, DR: subset of CUSCH-LOAD individuals from the Dominican Republic, PR: subset of CUSCH-LOAD individuals from Puerto Rico, SIGMA: the Slim Initiative in genomic medicine for the Americas.

In some populations, different results were seen between chromosome 7 and chromosome X. In LARGE-PD, the LOPO-all panel outperformed the LOPO-EA or LOPO-AL panels on both chromosomes. In LARGE-PD, the LOPO-EL panel performed better on chromosome 7 and the same as the LOPO-all panel on chromosome X (p-values = 2.04×10^-195^ & 0.553, respectively) (Tables S8 & S9). In the SIGMA cohort, the LOPO-EL outperformed the LOPO-all panel on both chromosomes 7 and X (p-values = 3.94×10^-173^ & 3.61×10^-165^, respectively) (Figure 3). The LOPO-AL panel outperformed the LOPO-all panel only on chromosome X (p-value = 3.65×10^-22^). In the Bambuí and Pelotas cohorts, the LOPO-all panel outperformed the LOPO-EA and LOPO-EL panels (p-values < 1.62×10^-32^), while there wasn’t a statistically significant difference between the LOPO-all and LOPO-AL panels (p-values > 0.0776). In CUSCH-LOAD, the panel LOPO-AL, LOPO-EL, and LOPO-all panels performed similarly on chromosome 7 (p-values > 0.99 in CUSCH-LOAD), while the LOPO-EL panel performed worse than LOPO-all (p-value = 7.65×10^-191^) and the LOPO-AL panel performed better than all on chromosome X (p-value =2.35×10^-122^). When we divided CUSCH-LOAD into two subpopulations - Puerto Rico (PR) and the Dominican Republic (DR), we observed different results. The DR subpopulation follows the same pattern as CUSCH-LOAD as a whole, while in the PR cohort, the LOPO-EL panel outperformed the LOPO-all panel on both chromosomes (chr7: p-value = 5.80×10^-38^; chr X: p-value = 2.25×10^-29^). In the PR subpopulation, the LOPO-all panel outperformed the LOPO-AL panel on chromosome 7, while the LOPO-AL panel outperformed the LOPO-all panel on chromosome X (p-values = 4.20×10^-8^ & 4.46×10^-10^, respectively).

In one cohort, the Salvador cohort, we observed the same results on both chromosomes. The LOPO-all panel outperformed the LOPO-EA and LOPO-EL panels (p-values < 4.50×10^-141^), and the LOPO-AL panel outperformed the LOPO-all panel on both chromosomes (p-values < 1.34×10^-307^). The LOPO-EL panel resulted in higher mean R^2^ values of imputed SNPs on both chromosomes in each population (Figure 4). A similar pattern as seen in the LOPO panels was seen in the NoLA panels, with the panels with lower sample sizes having higher proportions of common and well imputed SNPs (Figures S11 & S12).

**Figure 4.**
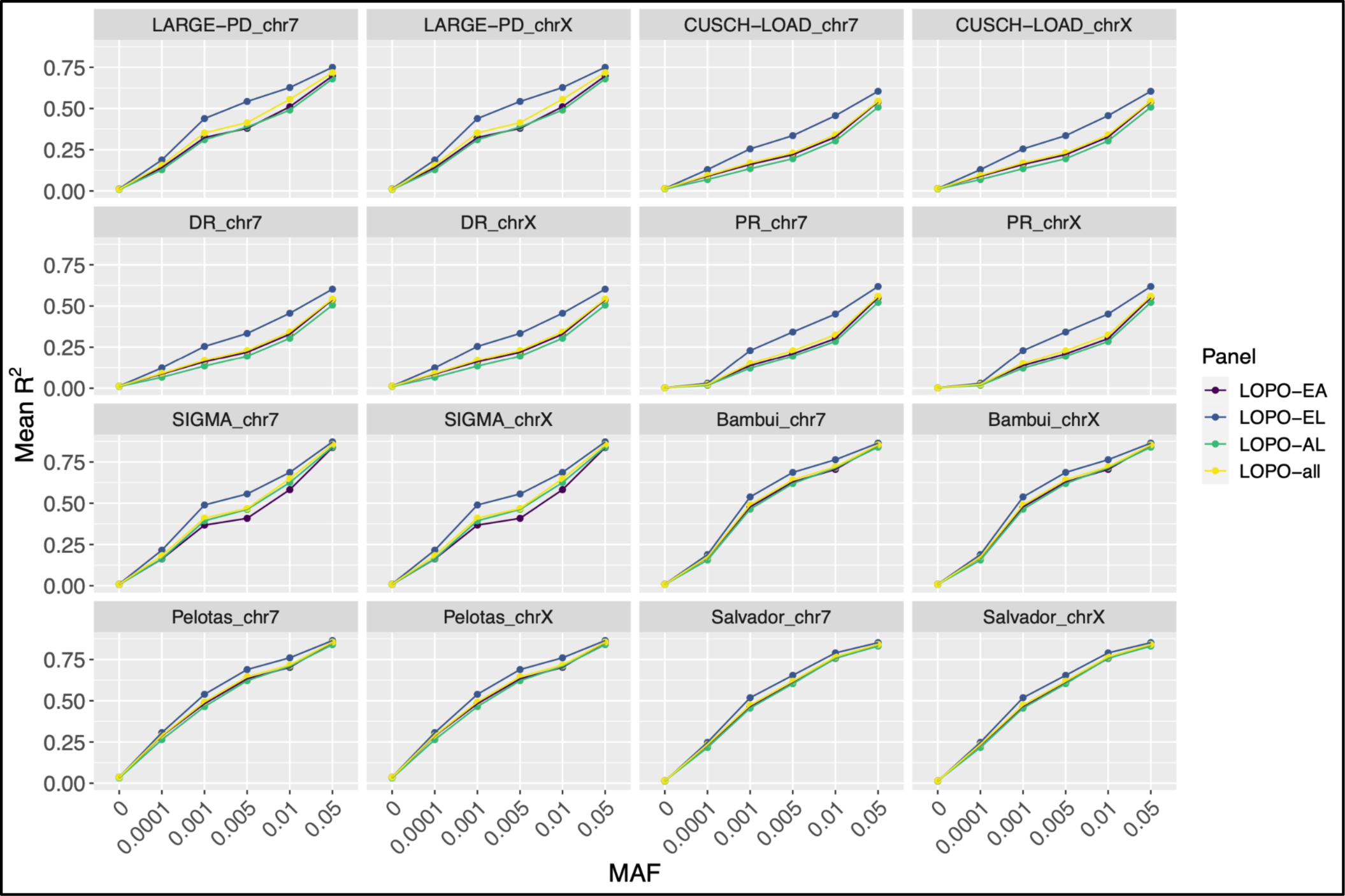
Mean R^2^ of imputed SNPs by MAF bin for LOPO reference panels. The mean R^2^ of imputed SNPs in each MAF bin for each reference panel in each population. Colors correspond to imputation reference panels which leave one population out (LOPO) of the imputation reference panels. LOPO-EA: Europeans and Africans included in reference panel, LOPO-EL: Europeans and Latin Americans included, LOPO-AL: Africans and Latin Americans included, LOPO-all: Africans, Europeans, and Latin Americans included, LARGE-PD: Latin American Research Consortium on the Genetics of Parkinson’s Disease, CUSCH-LOAD: Columbia University study of Caribbean Hispanics with familial or sporadic late-onset Alzheimer’s disease, DR: subset of CUSCH-LOAD individuals from the Dominican Republic, PR: subset of CUSCH-LOAD individuals from Puerto Rico, SIGMA: the Slim Initiative in genomic medicine for the Americas.

### Increasing proportion of IA-like ancestry (PIAA)

Comparing imputation quality through empirical R^2^ values allows us to see how increasing IA ancestry proportions in the reference panel affects imputation quality and see differences between cohorts and chromosomes. Increasing the IA ancestry proportions of Latin Americans included in the imputation reference panel was beneficial in each population except on chromosome X in the Pelotas cohort, where there was no significant difference between panels PIAA-1 and PIAA-1 (p-value > 0.99).

In CUSCH-LOAD, DR, Bambuí, Salvador, and Pelotas (only chr 7), improvements in imputation quality were only seen up to PIAA-2 or PIAA-3. In CUSCH-LOAD as a whole, panel PIAA-3 performed better than PIAA-1 on chromosome 7 and X (p-values = 2.38X10^-16^ & 7.22×10^-99^, respectively) (Figure 5, Tables S10 & S11). Panels PIAA-1, PIAA-4 and PIAA-5 compared similarly on chr 7 ( p-values > 0.327), and panels PIAA-4 and PIAA-5 performed better than panel PIAA-1 on chromosome X (p-values < 4.54×10^-142^). In CUSCH-LOAD, similar results were found only for individuals from the DR. We observed similar findings between each of the EPIGEN-Brazil cohorts, with panels PIAA-2 and PIAA-3 outperforming panel PIAA-1 on chromosome 7 in all three sub-cohorts (p-values < 4.85×10^-7^), and panel PIAA-1 outperforming PIAA-5 - PIAA-9 in all three (p-values < 1.22×10^-8^). Panels PIAA-4 - PIAA-9 also performed worse in all three cohorts than panel PIAA-1 on chromosome X (p-values < 3.43×10^-15^) (Figures 5 & S13, Tables S10 & S11).

**Figure 5.**
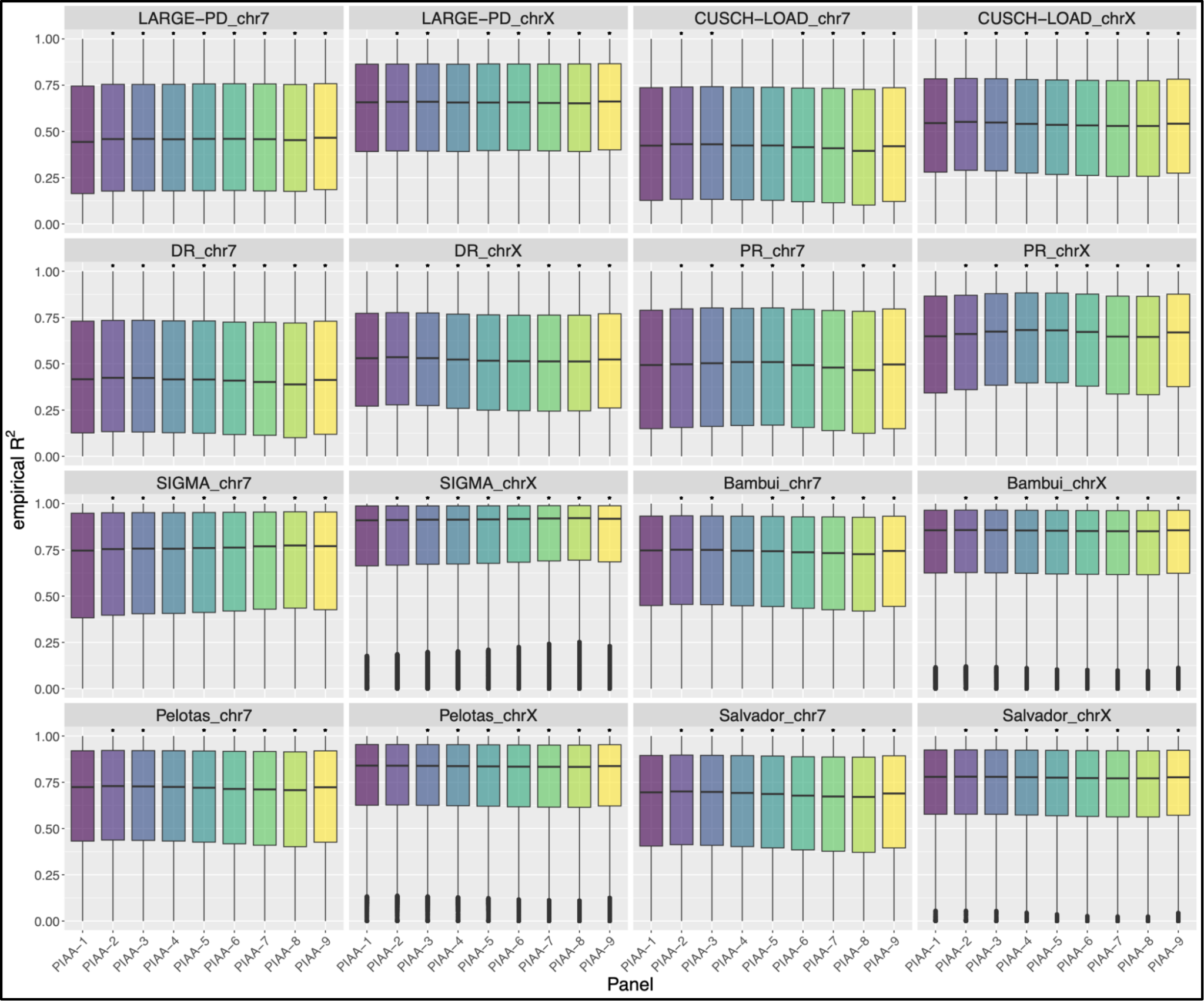
Comparison of empirical R^2^ values for genotyped SNPs using PIAA reference panels. Empirical R^2^ values for each population, using imputation reference panels containing increasing proportions of Indigenous American ancestry (PIAA) of included Latin American individuals. Pairwise comparisons done with panel 1 as a reference using a paired, two-sided Wilcoxon signed rank test. *: p-value < 0.0014. LARGE-PD: Latin American Research Consortium on the Genetics of Parkinson’s Disease, CUSCH-LOAD: Columbia University study of Caribbean Hispanics with familial or sporadic late-onset Alzheimer’s disease, DR: subset of CUSCH-LOAD individuals from the Dominican Republic, PR: subset of CUSCH-LOAD individuals from Puerto Rico, SIGMA: the Slim Initiative in genomic medicine for the Americas.

In LARGE-PD, PR, and SIGMA, improvements in imputation quality were seen with higher increases in the IA ancestry proportion of Latin Americans included in the reference panel. On both chromosomes in the PR cohort, panels PIAA-2, PIAA-3, PIAA-4, PIAA-5, PIAA-6, and PIAA-9 outperformed panel PIAA-1, and panels PIAA-8 performed worse than PIAA-1 (chr7, p-value = 1.34×10^-14^; chrX, 1.37×10^-45^). There was no difference seen in empirical R^2^ values in panels PIAA-1 and PIAA-7 on chr 7 (p-value > 0.99), but PIAA-7 resulted in lower values on chr X (p-value = 7.29×10^-31^). In both SIGMA and LARGE-PD, all panels resulted in better imputation on chr 7, and panels PIAA-2, PIAA-3, and PIAA-5 - PIAA-9 performed better on X. In the SIGMA cohort, panel PIAA-4 performed better than panel PIAA-1 on chromosome X (p-value = 8.18×10^-202^), and no difference was seen between panels PIAA-1 and PIAA-4 on chromosome X in the LARGE-PD cohort (p-value = 0.456).

There were also differences seen between cohorts in the magnitude of the effect of increasing IA-like proportions in the imputation reference panels between chromosome X and 7. In CUSCH-LOAD, as well as the PR and DR cohorts, there were stronger medians of differences on chromosome X compared to 7. Alternatively, in the LARGE-PD and SIGMA cohort, higher medians were seen on chromosome 7 compared to chromosome X. The biggest differences in mean R^2^ between panels was seen in the SIGMA and LARGE-PD cohorts (Figure 6). Again, we observed more common and well imputed variants in the panels resulting in fewer SNPs (Figures S14 & S15).

**Figure 6.**
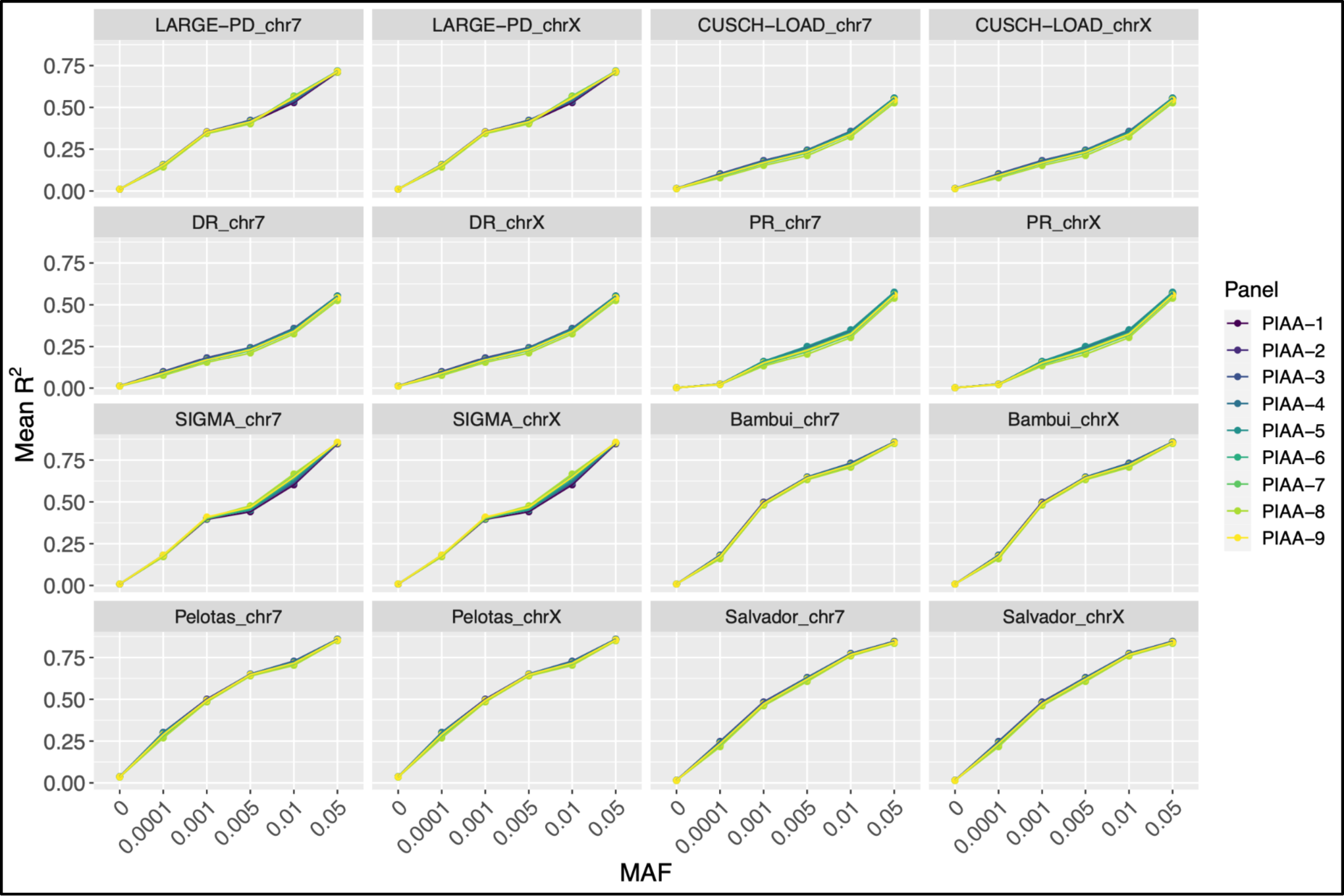
Mean R^2^ of imputed SNPs by MAF bin for PIAA reference panels. The mean R^2^ of imputed SNPs in each MAF bin for each reference panel in each population. Colors correspond to imputation reference panels including Latin American individuals with increasing proportions of Indigenous American ancestry (PIAA). LARGE-PD: Latin American Research Consortium on the Genetics of Parkinson’s Disease, CUSCH-LOAD: Columbia University study of Caribbean Hispanics with familial or sporadic late-onset Alzheimer’s disease, DR: subset of CUSCH-LOAD individuals from the Dominican Republic, PR: subset of CUSCH-LOAD individuals from Puerto Rico, SIGMA: the Slim Initiative in genomic medicine for the Americas.

### Comparing females only with mixed reference panels

Differences in imputation quality between chromosomes 7 and X could be due to differences in sample size, due to there being fewer copies of the X chromosome than chromosome 7 within the same population. To address this, we created two imputation reference panels, one containing both males and females, and one containing only females. When comparing these results, we observed no difference between the panels (Figure S16).

## Discussion

There is limited research analyzing how the composition of the imputation reference panel affects imputation quality in Latin American cohorts. Our study aimed to analyze this in four populations including individuals from: the Caribbean, South America, Mexico, and Brazil. We created sub-populations for CUSCH-LOAD, analyzing individuals from the Dominican Republic and Puerto Rico together and independently, as well as separating the Brazilian cohort into three populations: Bambuí, Pelotas, and Salvador. Our work expands upon previous research, extending to various alterations of imputation panels and including the X chromosome. Similar to previous studies, we found that increasing the Indigenous American ancestry of individuals included in the reference panel did not increase imputation quality in all Latin American populations ^[18]^.

Increasing the number of Latin Americans included in the imputation reference panel had different effects in different populations. However, having 4,000 Latin American individuals in the reference panel was significantly better than having 1,000. Importantly, the effect of increasing the numbers of Latin Americans had different effects of chromosome 7 and X in the same populations. In some populations, like CUSCH-LOAD, increasing the number of Latin American individuals to 5,000 had stronger effects on chr 7 than X, even though both resulted in better imputation compared to having 4,000. In the Salvador cohort, having 5,000 Latin American individuals resulted in better imputation than having 4,000 on chromosome 7, but there was no additional benefit seen of increasing the number of Latin Americans beyond 3,000 for chromosome X.

Similarly, in the LOPO comparison, different effects were seen on chromosome X and 7, but also between cohorts. In cohorts with high average African ancestry proportions (CUSCH-LOAD, DR, and Salvador), LOPO-all outperformed LOPO-EL, indicating that having Africans included in the reference panel was especially important. LOPO-AL outperformed LOPO-all on the X chromosome in Salvador, further highlighting this importance. In LARGE-PD and SIGMA, the two cohorts with the highest average Indigenous American ancestry proportions and low average African ancestry proportions, LOPO-EL outperformed all other imputation reference panels. Three populations had high average European ancestry proportions - Bambuí, Pelotas, and PR. In Bambuí and Pelotas, LOPO-all performed the best, while in PR LOPO-EL performed the best. This may be due to PR having nearly twice the average Indigenous American ancestry proportion of Bambuí or Pelotas. It is also important to highlight how different panels performed the best within the CUSCH-LOAD cohort. CUSCH-LOAD as a whole and DR had similar results, as well as similar average ancestry proportions, while PR has higher average European and Indigenous American ancestry proportions.

Increasing Indigenous American ancestry was beneficial in every population except on chromosome X in the Pelotas cohort, which may be due to the low average Indigenous American ancestry in this population (7.4%). However, there were differences in the level at which increasing IA ancestry proportions of the Latin Americans included in the panel became detrimental to imputation quality between populations and chromosomes in some populations. CUSCH-LOAD, DR, Bambuí, Pelotas, and Salvador had the lowest average Indigenous American ancestry proportions (up to 9.4%) of all the target populations. In these cohorts, improvements in imputation quality were only seen up to PIAA-2 or PIAA-3. In PR, with an average Indigenous American ancestry of 12.4%, PIAA-6 outperformed PIAA-1. The gain in imputation quality was seen even further in populations with high average Indigenous American ancestry, such as LARGE-PD and SIGMA, where PIAA-8 outperformed PIAA-1 on chromosome 7. This highlights the importance of taking ancestry into consideration when selecting which Latin American individuals to include in the imputation reference panel.

One limitation of this study is while the magnitude of differences between imputation panels are statistically significant, they are not very large. This could be due to a limited sample size, which could be expanded upon with further studies including larger numbers of Latin American individuals. It could also be due to complexities of studying admixed populations and the zoomed-out approach of looking at global ancestry. Investigating local ancestry and the impact on imputation quality in specific ancestry tracts may highlight larger differences in affect. While the genome-wide trend might display small differences, there are still examples of larger differences between populations within a single cohort or country.

Large sample sizes containing more diverse Latin American populations are needed to validate the results described here. Further studies looking at the impact of including diverse Latin American samples compared to individuals from similar populations in the imputation reference panel are also warranted. These results highlight the heterogeneity of Latin Americans and the importance of not viewing this superpopulation as a single entity. Furthermore, when exclusively studying the X chromosome, a different reference panel than what would be best for the autosomes may result in better imputation quality. Developing imputation panels for Latin American populations will need to take into account the population structure of the target population, as well as their history, and will vary from place to place and person to person.

## Acknowledgements

T32 AG000262, LARGE-PD RO1 NS112499, R35 HG010692, ASAP-GP2, Michael J Fox Foundation, Parkinson’s Foundation, Conselho Nacional de Desenvolvimento Científico e Tecnológico (Brazil), Brazilian Ministry of Health (Genomas Brasil Program), Fundação de Amparo a Pesquisa de Minas Gerais (FAPEMIG).

## Notes

### Competing Interest Statement

The authors have declared no competing interest.

