## Supplementary Figures for "Comparing the effect of imputation reference panel composition in four distinct Latin American cohorts"

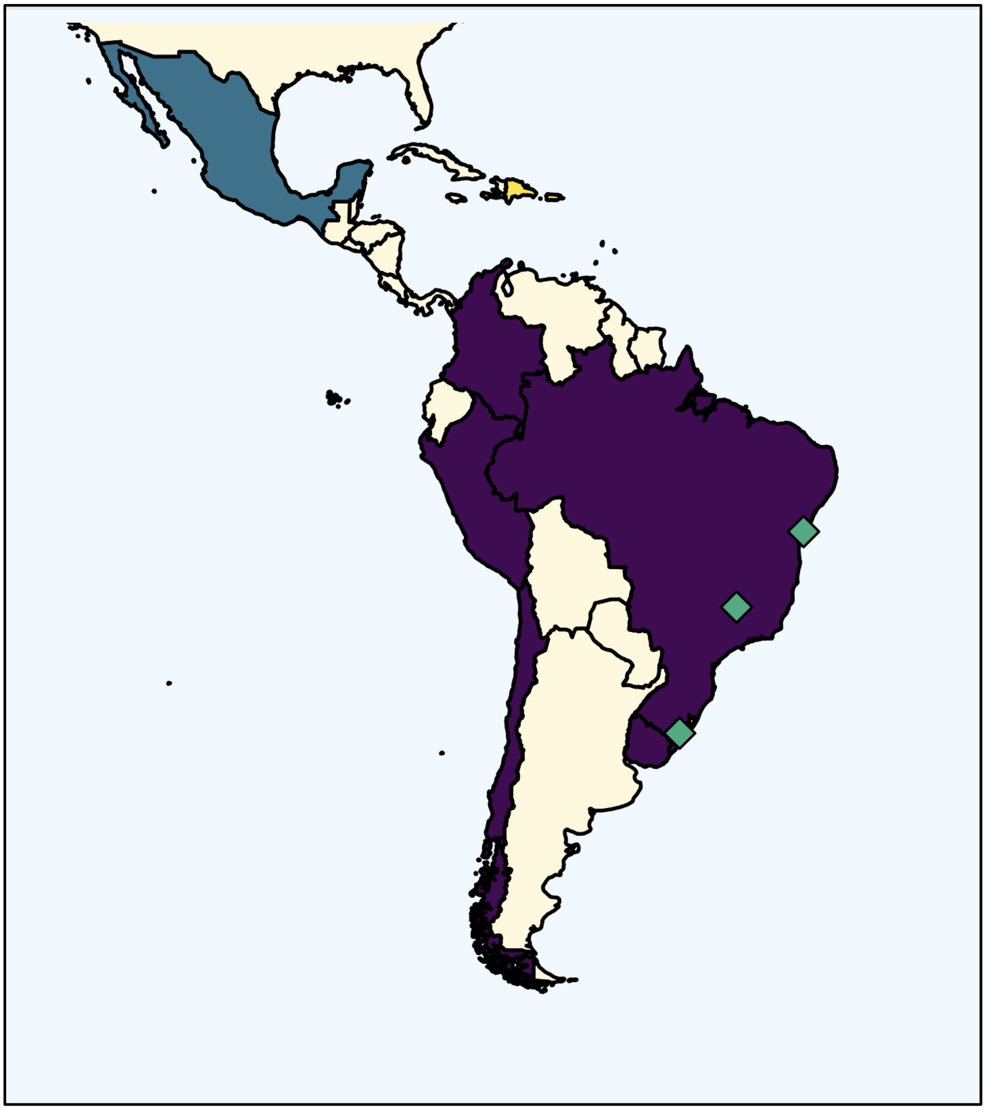

**Figure S1. Map of target populations for imputation.** Four target populations are used: 1) the Latin American Research Consortium on the Genetics of Parkinson's Disease (LARGE-PD) (purple), 2) the Columbia University Study of Caribbean Hispanics with Familial or Sporadic Late Onset Alzheimer's Disease (CUSCH-LOAD) (yellow), 3) the Slim Initiative in Genomic Medicine for the Americas (SIGMA) (blue), and 4) Genetic Epidemiology of Complex Diseases in Brazilian population-based cohorts (EPIGEN)-Brazil (green diamonds). CUSCH-LOAD was analyzed as a whole and split into individuals from Puerto-Rico and the Dominican Republic.

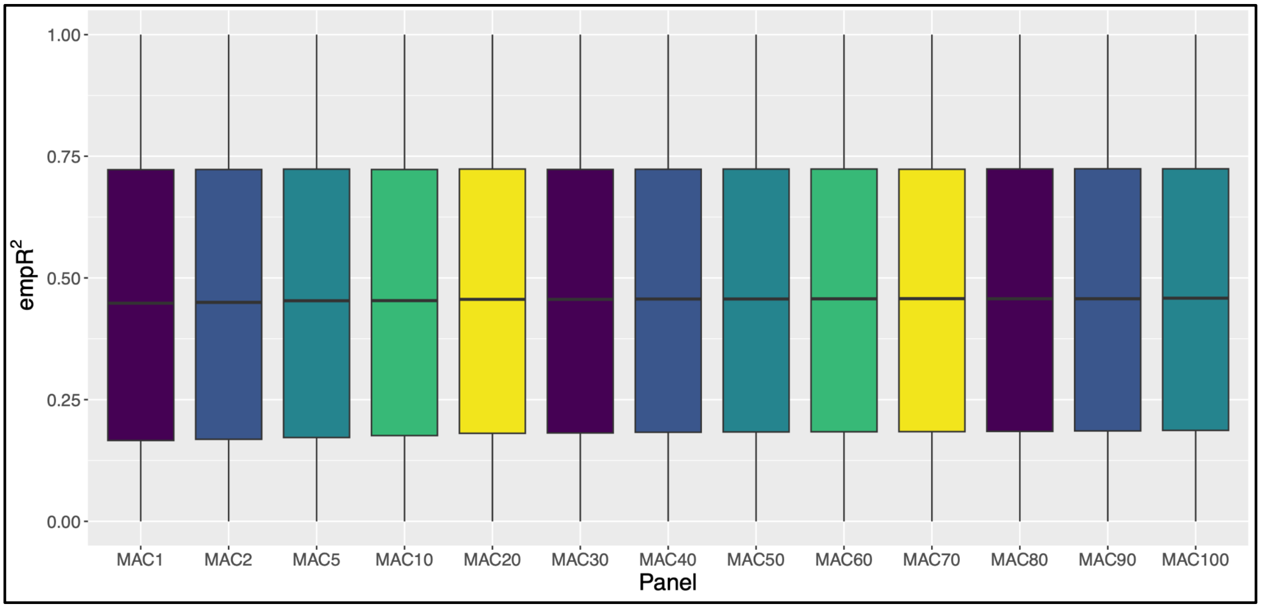


**Figure S2. Boxplot of empR$^2$ values using reference panels with various MAC cutoffs.** Boxplot of genotyped SNPs in LARGE-PD when imputed using 13 reference panels with increasing MAC cutoffs.

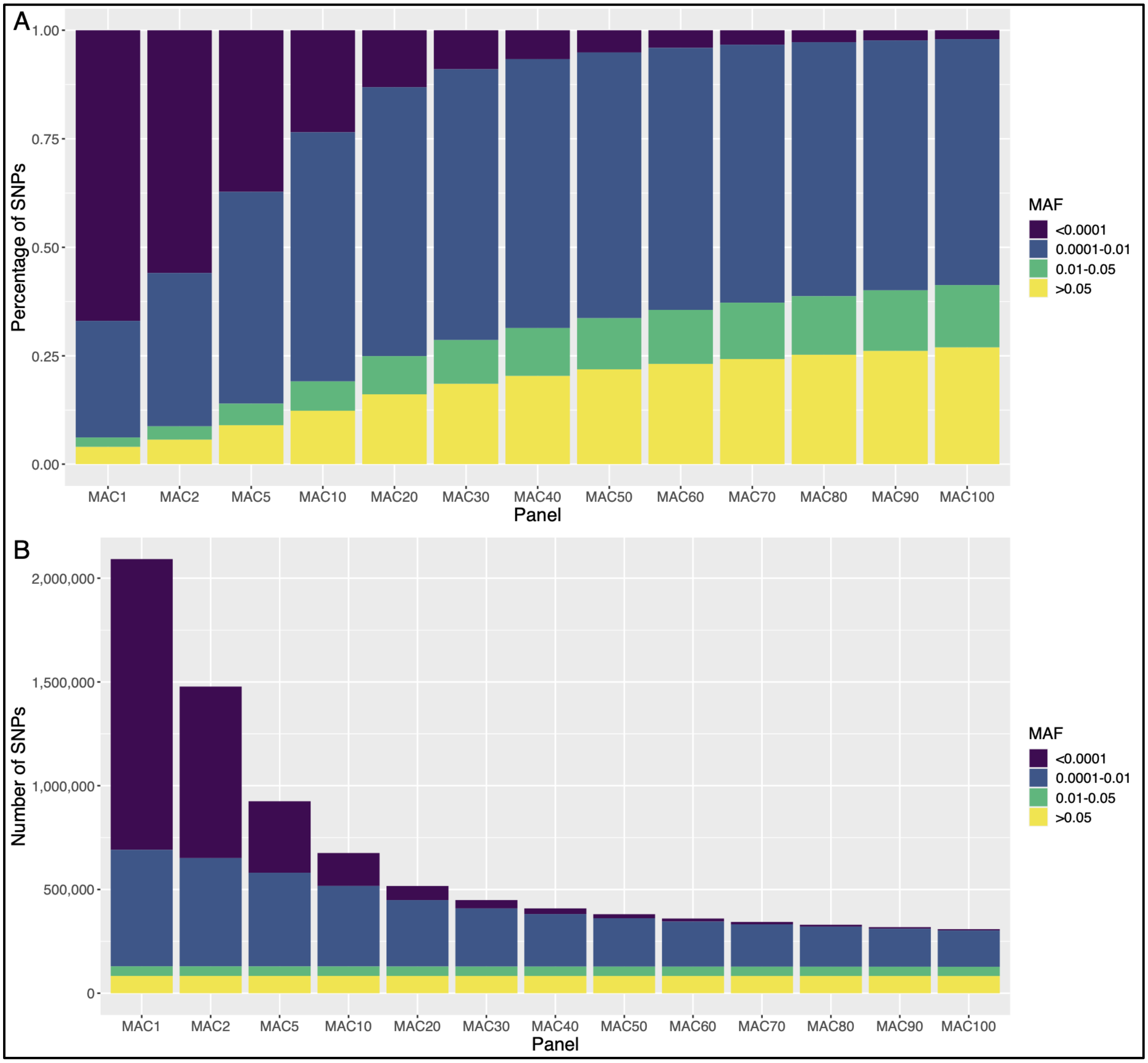


**Figure S3. Comparison of MAF of imputed SNPs in MAC comparison reference panels.** a) The percentage of SNPs in each MAF category by population and imputation reference panel. b) The number of SNPs in each MAF category by population and imputation reference panel.

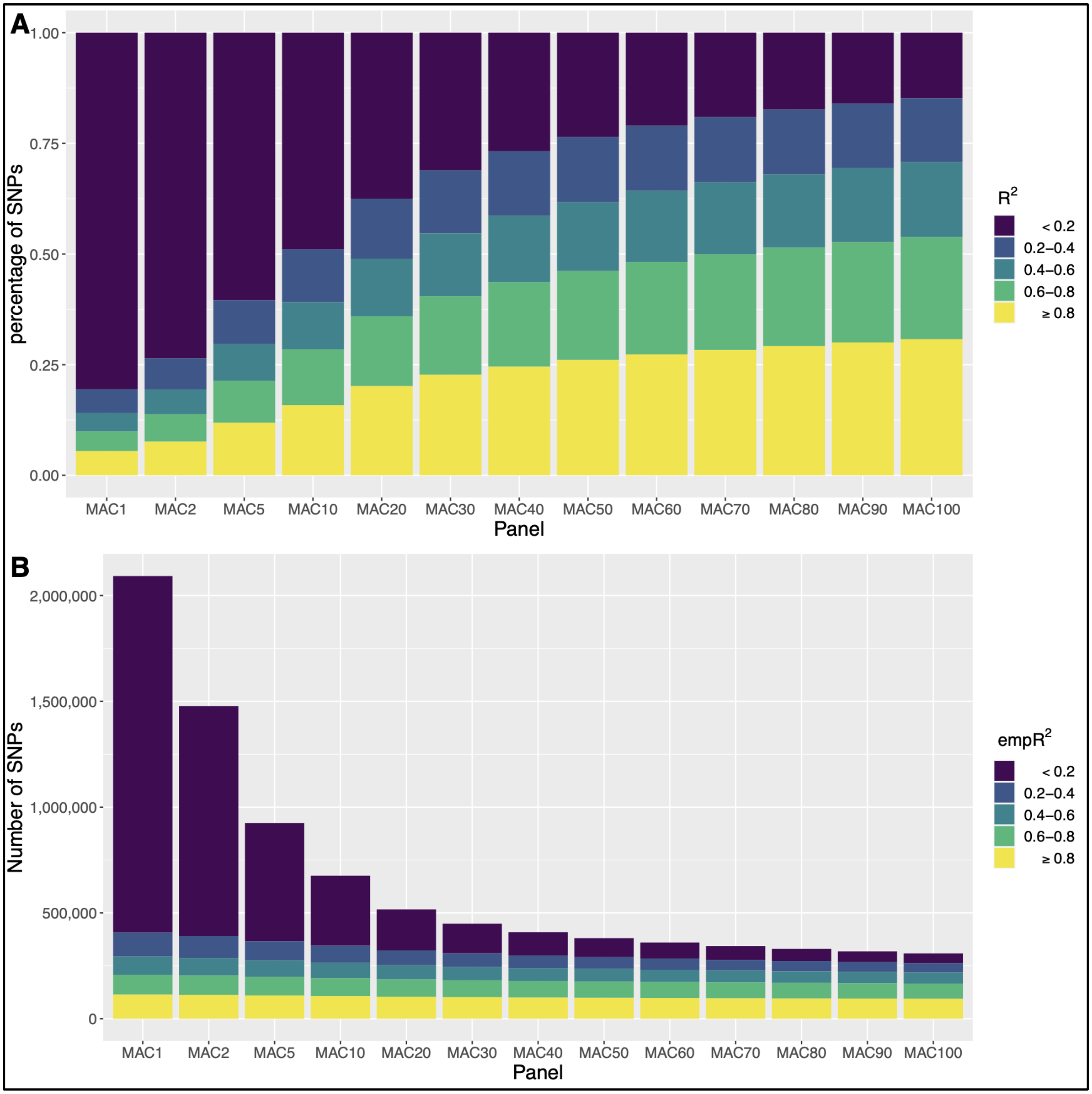


**Figure S4. Comparison of $R^2$ of imputed SNPs in MAC comparison reference panels.** a) The percentage of SNPs in each $R^2$ category by population and imputation reference panel. b) The number of SNPs in each $R^2$ category by population and imputation reference panel.

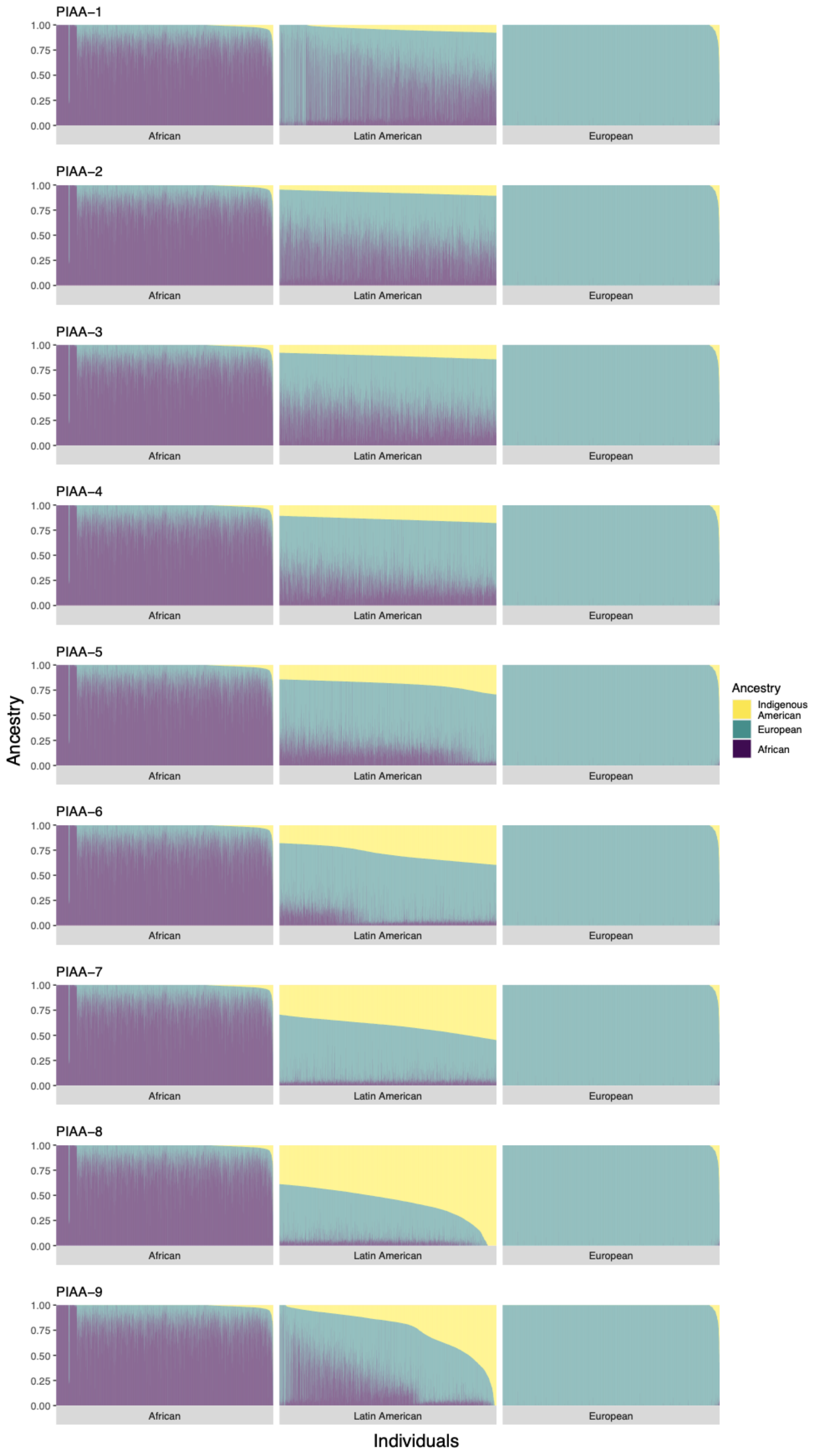

PIAA–1
PIAA–2
PIAA–3
PIAA–4
PIAA–5
PIAA–6
PIAA–7
PIAA–8
PIAA–9
1.00
0.75
0.50
0.25
0.00
African
Latin American
European
Ancestry
Indigenous American
European
African
Ancestry
Individuals

**Figure S5. Admixture plot of individuals included in PIAA imputation reference panels.** Ancestry proportions for each individual in each imputation reference panel containing increasing proportions of Indigenous American ancestry (PIAA) of included Latin American individuals, using the 10000 Genomes Project as a reference. LARGE-PD: Latin American Research Consortium on the Genetics of Parkinson's Disease, CUSCH-LOAD: Columbia University study of Caribbean Hispanics with familial or sporadic late-onset Alzheimer's disease, DR: subset of CUSCH-LOAD individuals from the Dominican Republic, PR: subset of CUSCH-LOAD individuals from Puerto Rico, SIGMA: the Slim Initiative in genomic medicine for the Americas.

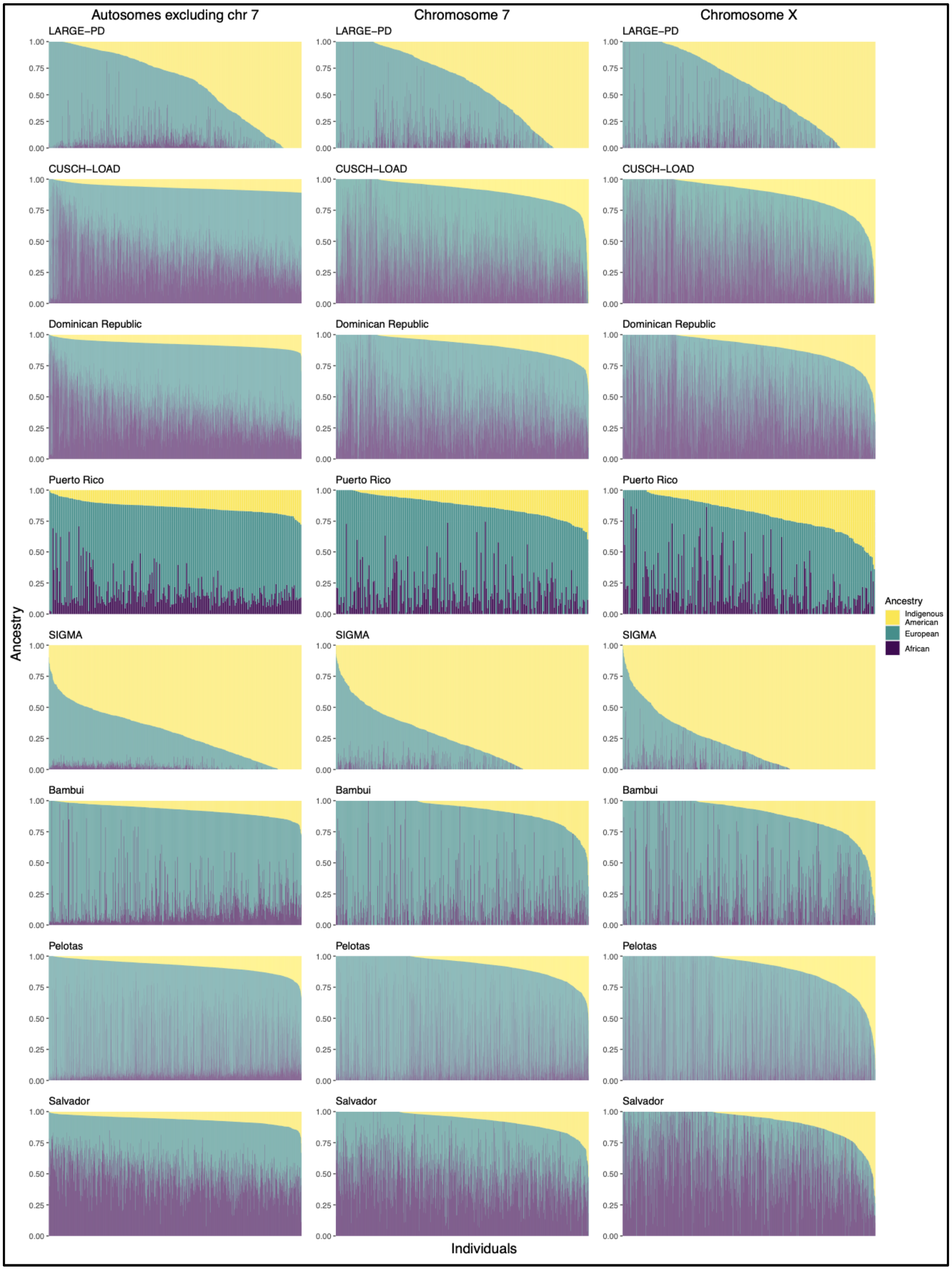


**Figure S6. Admixture plot of individuals in each target population.** Ancestry proportions for each individual in each target population based on 1) autosomes excluding chromosome 7; 2) chromosome 7, and 3) chromosome X. LARGE-PD: Latin American Research Consortium on the Genetics of Parkinson's Disease, CUSCH-LOAD: Columbia University study of Caribbean

Hispanics with familial or sporadic late-onset Alzheimer's disease, DR: subset of CUSCH-LOAD individuals from the Dominican Republic, PR: subset of CUSCH-LOAD individuals from Puerto Rico, SIGMA: the Slim Initiative in genomic medicine for the Americas.

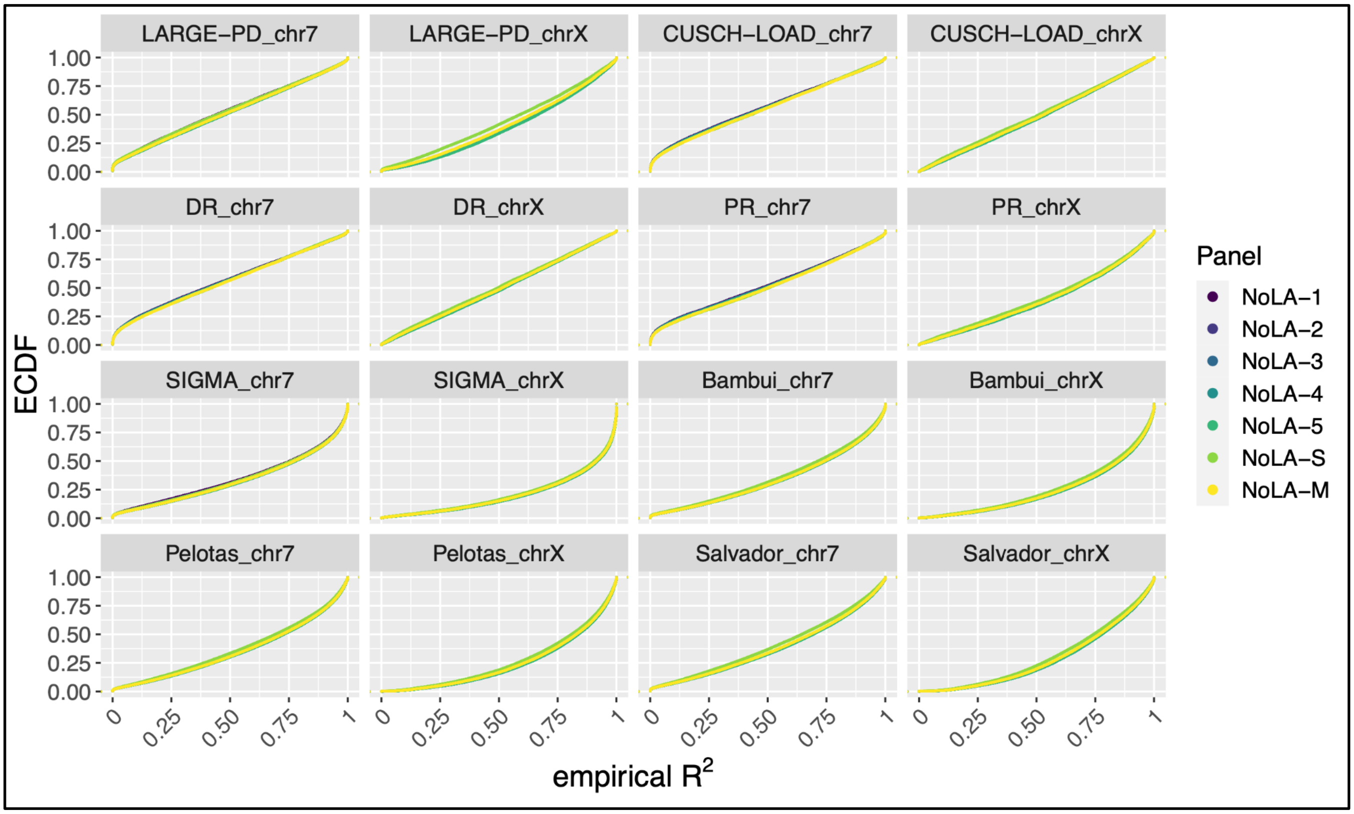


**Figure S7. Empirical Cumulative Distribution Function (ECDF) of NoLA reference panels.** ECDF of imputation reference panels with imputation reference panels containing varying numbers of Latin American individuals (NoLA). The number of Latin Americans increase from panel NoLA-1 to NoLA-5. NoLA-S, NoLA-M, and NoLA-4 contain equal proportions of African, European, and Latin American individuals, with increasing total sample size. LARGE-PD: Latin American Research Consortium on the Genetics of Parkinson's Disease, CUSCH-LOAD: Columbia University study of Caribbean Hispanics with familial or sporadic late-onset Alzheimer's disease, DR: subset of CUSCH-LOAD individuals from the Dominican Republic, PR: subset of CUSCH-LOAD individuals from Puerto Rico, SIGMA: the Slim Initiative in genomic medicine for the Americas.

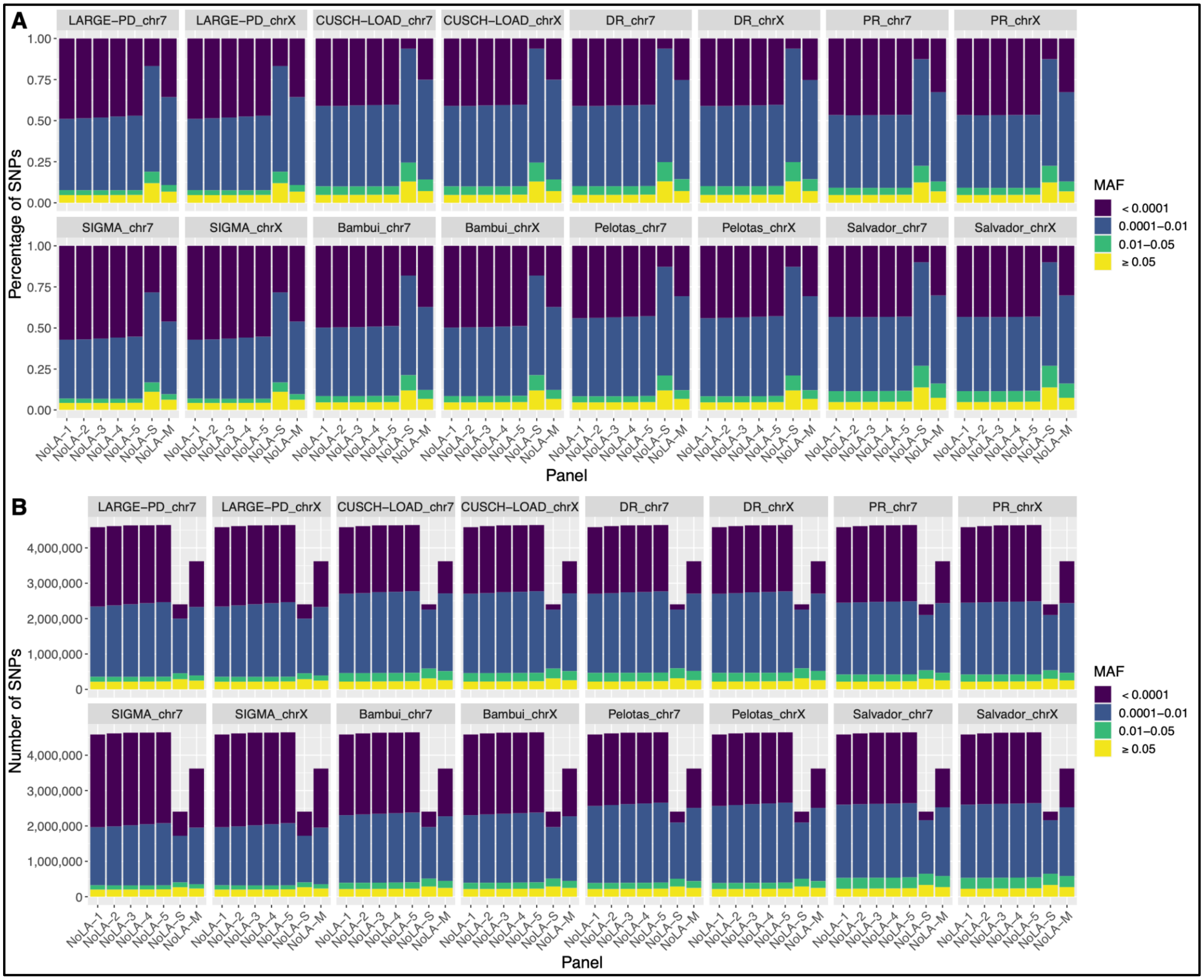


**Figure S8. Comparison of MAF of imputed SNPs in NoLA panels.** a) The percentage of SNPs in each MAF category by population and imputation reference panel. b) The number of SNPs in each MAF category by population and imputation reference panel. The number of Latin Americans increase from panel NoLA-1 to NoLA-5. NoLA-S, NoLA-M, and NoLA-4 contain equal proportions of African, European, and Latin American individuals, with increasing total sample size. LARGE-PD: Latin American Research Consortium on the Genetics of Parkinson's Disease, CUSCH-LOAD: Columbia University study of Caribbean Hispanics with familial or sporadic late-onset Alzheimer's disease, DR: subset of CUSCH-LOAD individuals from the Dominican Republic, PR: subset of CUSCH-LOAD individuals from Puerto Rico, SIGMA: the Slim Initiative in genomic medicine for the Americas.

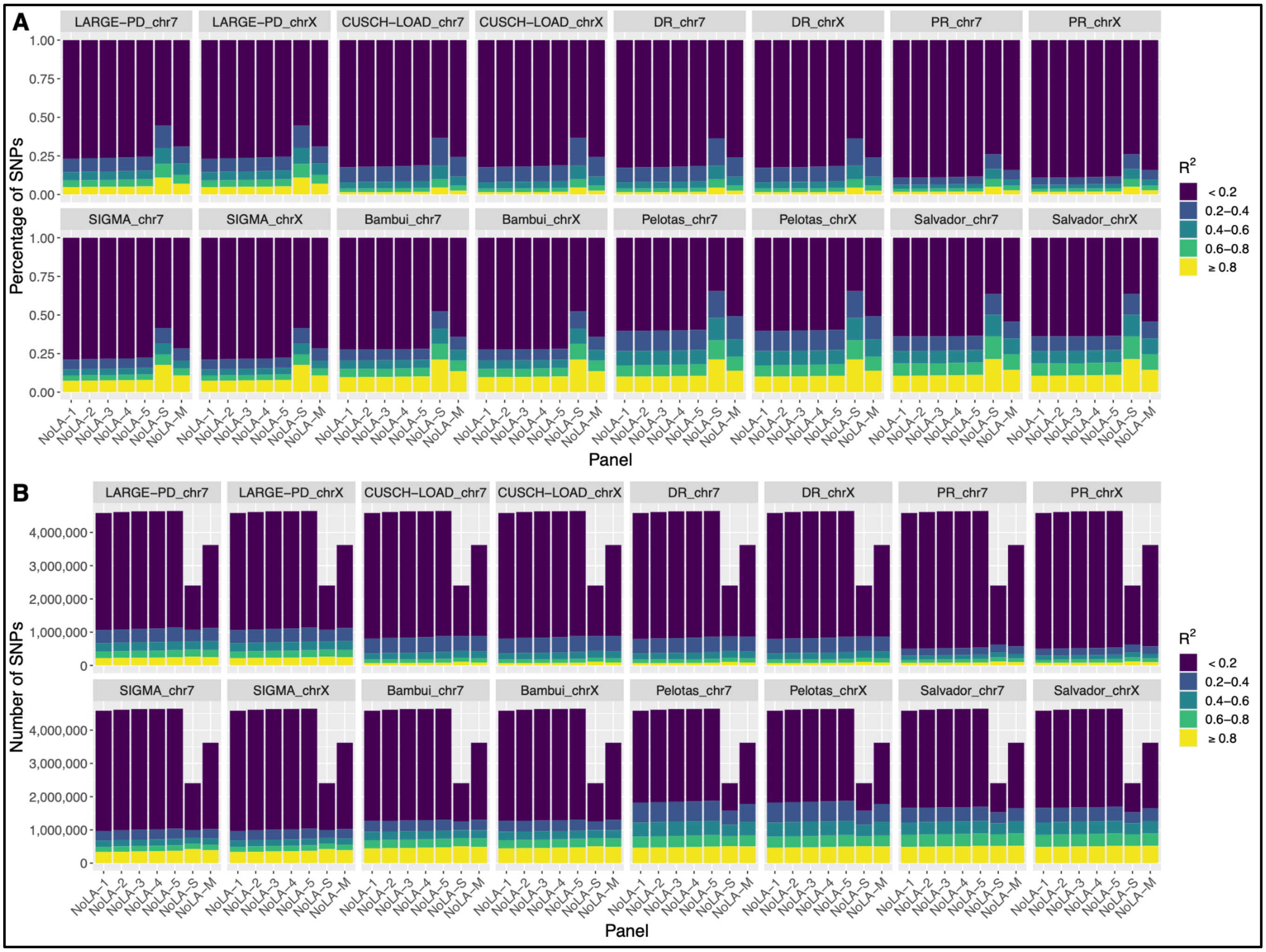


**Figure S9. Comparison of $R^2$ of imputed SNPs in NoLA panels.** a) The percentage of SNPs in each $R^2$ category by population and imputation reference panel. b) The number of SNPs in each $R^2$ category by population and imputation reference panel. The number of Latin Americans increase from panel NoLA-1 to NoLA-5. NoLA-S, NoLA-M, and NoLA-4 contain equal proportions of African, European, and Latin American individuals, with increasing total sample size. LARGE-PD: Latin American Research Consortium on the Genetics of Parkinson's Disease, CUSCH-LOAD: Columbia University study of Caribbean Hispanics with familial or sporadic late-onset Alzheimer's disease, DR: subset of CUSCH-LOAD individuals from the Dominican Republic, PR: subset of CUSCH-LOAD individuals from Puerto Rico, SIGMA: the Slim Initiative in genomic medicine for the Americas.

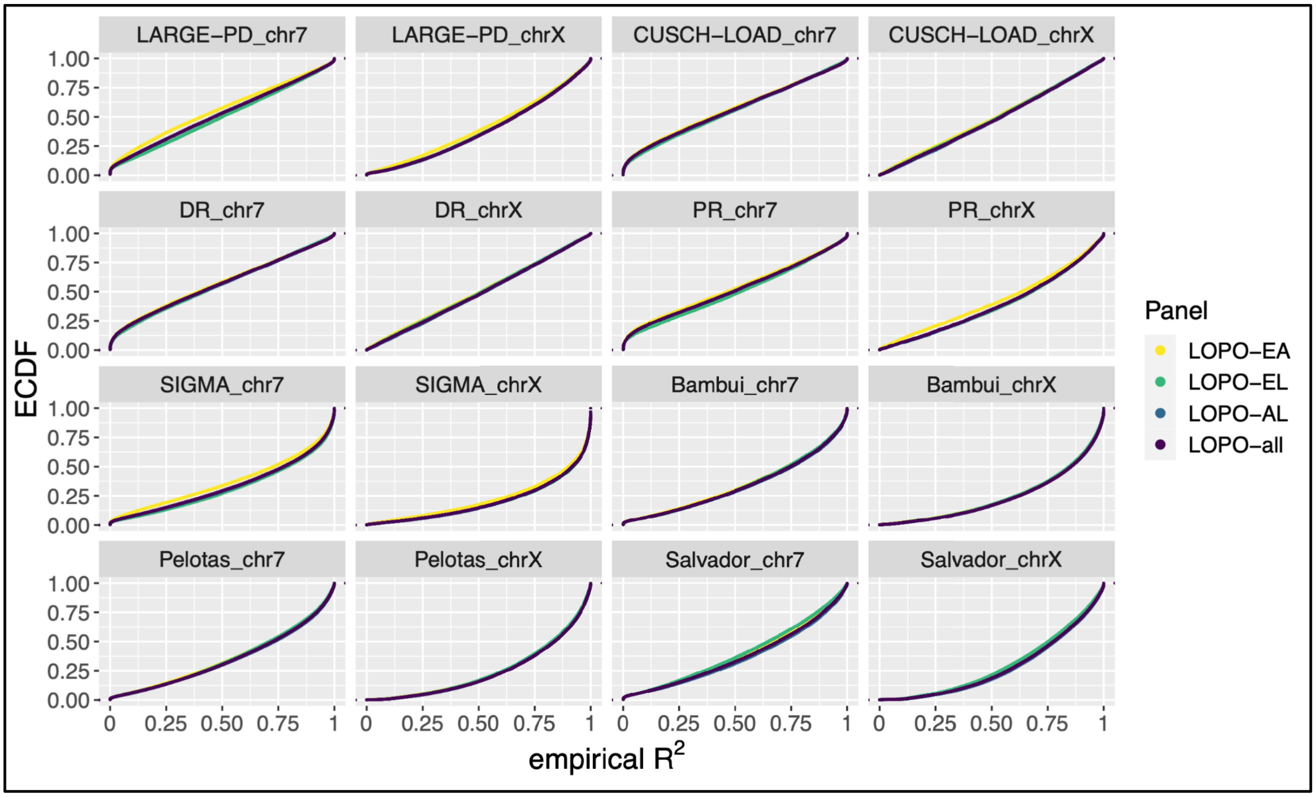


**Figure S10. ECDF of LOPO reference panels.** ECDF of imputation reference panels, when leaving one population out (LOPO) of the imputation reference panels. LOPO-EA: Europeans and Africans included in reference panel, LOPO-EL: Europeans and Latin Americans included, LOPO-AL: Africans and Latin Americans included, LOPO-all: Africans, Europeans, and Latin Americans included, LARGE-PD: Latin American Research Consortium on the Genetics of Parkinson's Disease, CUSCH-LOAD: Columbia University study of Caribbean Hispanics with familial or sporadic late-onset Alzheimer's disease, DR: subset of CUSCH-LOAD individuals from the Dominican Republic, PR: subset of CUSCH-LOAD individuals from Puerto Rico, SIGMA: the Slim Initiative in genomic medicine for the Americas.

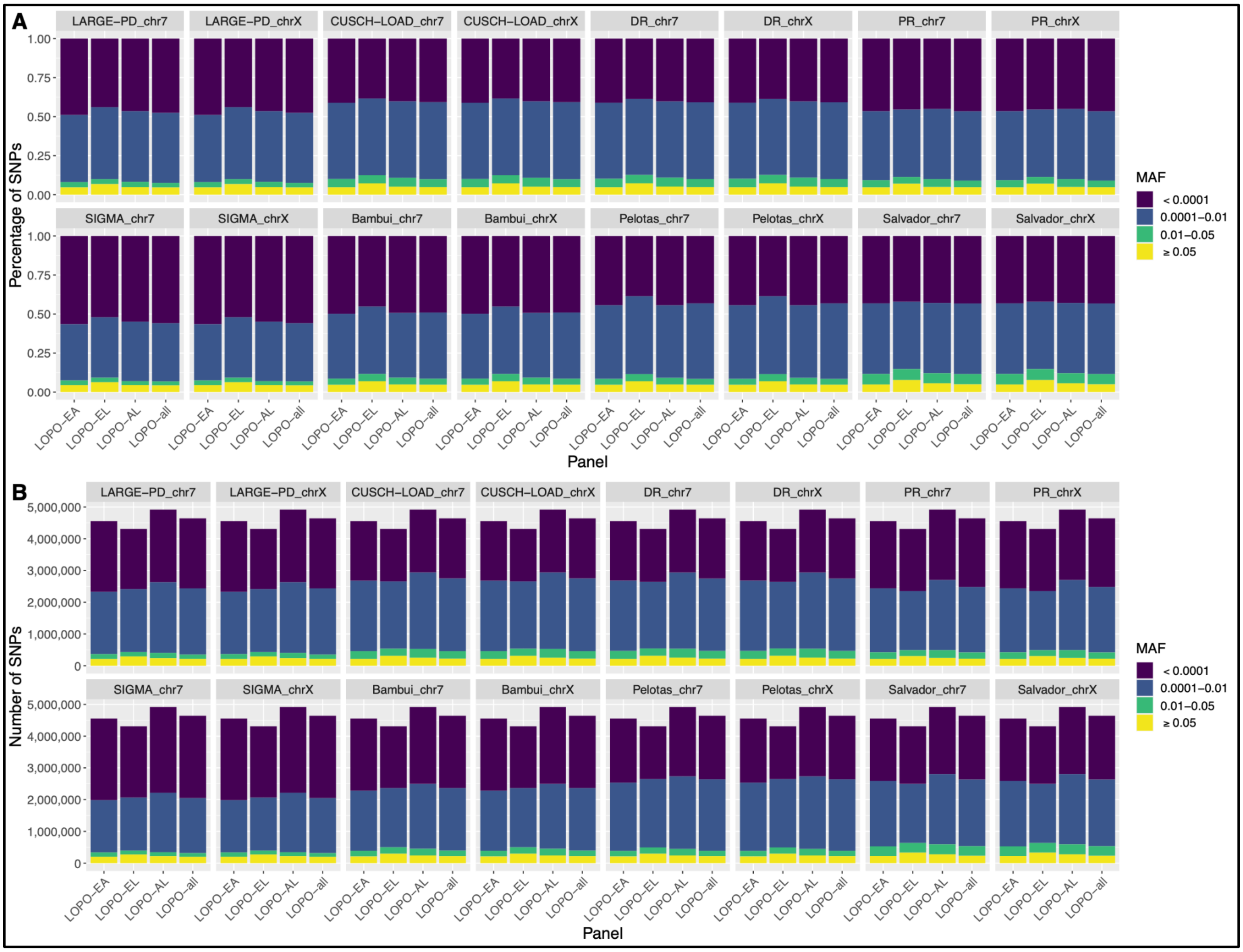


**Figure S11. Comparison of MAF of imputed SNPs in LOPO panels.** a) The percentage of SNPs in each MAF category by population and imputation reference panel. b) The number of SNPs in each MAF category by population and imputation reference panel. LOPO-EA: Europeans and Africans included in reference panel, LOPO-EL: Europeans and Latin Americans included, LOPO-AL: Africans and Latin Americans included, LOPO-all: Africans, Europeans, and Latin Americans included, LARGE-PD: Latin American Research Consortium on the Genetics of Parkinson’s Disease, CUSCH-LOAD: Columbia University study of Caribbean Hispanics with familial or sporadic late-onset Alzheimer’s disease, DR: subset of CUSCH-LOAD individuals from the Dominican Republic, PR: subset of CUSCH-LOAD individuals from Puerto Rico, SIGMA: the Slim Initiative in genomic medicine for the Americas.

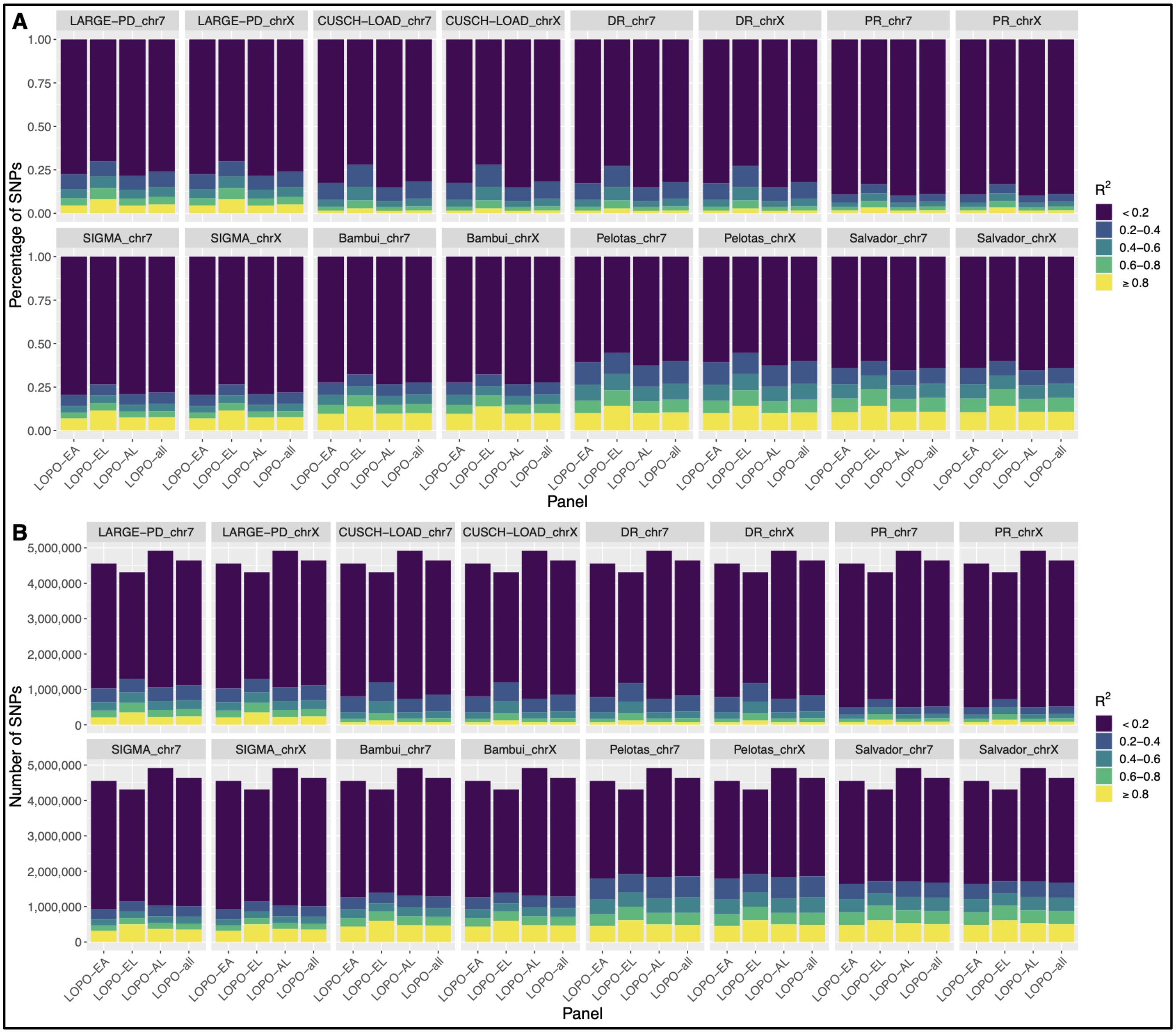


**Figure S12. Comparison of $R^2$ of imputed SNPs in LOPO panels.** a) The percentage of SNPs in each $R^2$ category by population and imputation reference panel. b) The number of SNPs in each $R^2$ category by population and imputation reference panel. LOPO-EA: Europeans and Africans included in reference panel, LOPO-EL: Europeans and Latin Americans included, LOPO-AL: Africans and Latin Americans included, LOPO-all: Africans, Europeans, and Latin Americans included, LARGE-PD: Latin American Research Consortium on the Genetics of Parkinson's Disease, CUSCH-LOAD: Columbia University study of Caribbean Hispanics with familial or sporadic late-onset Alzheimer's disease, DR: subset of CUSCH-LOAD individuals from the Dominican Republic, PR: subset of CUSCH-LOAD individuals from Puerto Rico, SIGMA: the Slim Initiative in genomic medicine for the Americas.

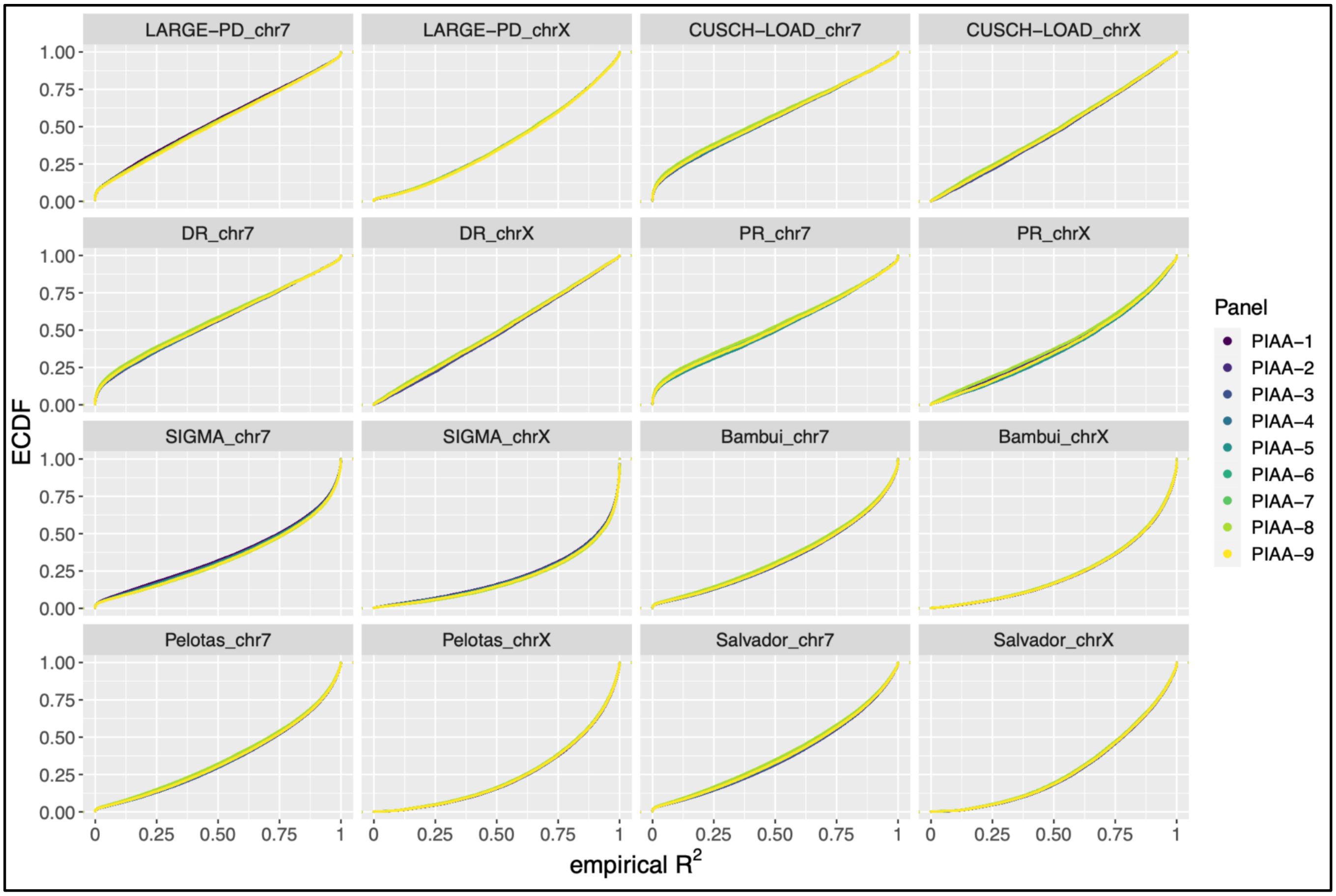


**Figure S13. ECDF of PIAA reference panels.** ECDF of imputation reference panels containing increasing proportions of Indigenous American ancestry (PIAA) of included Latin American individuals, in each target population. LARGE-PD: Latin American Research Consortium on the Genetics of Parkinson's Disease, CUSCH-LOAD: Columbia University study of Caribbean Hispanics with familial or sporadic late-onset Alzheimer's disease, DR: subset of CUSCH-LOAD individuals from the Dominican Republic, PR: subset of CUSCH-LOAD individuals from Puerto Rico, SIGMA: the Slim Initiative in genomic medicine for the Americas.

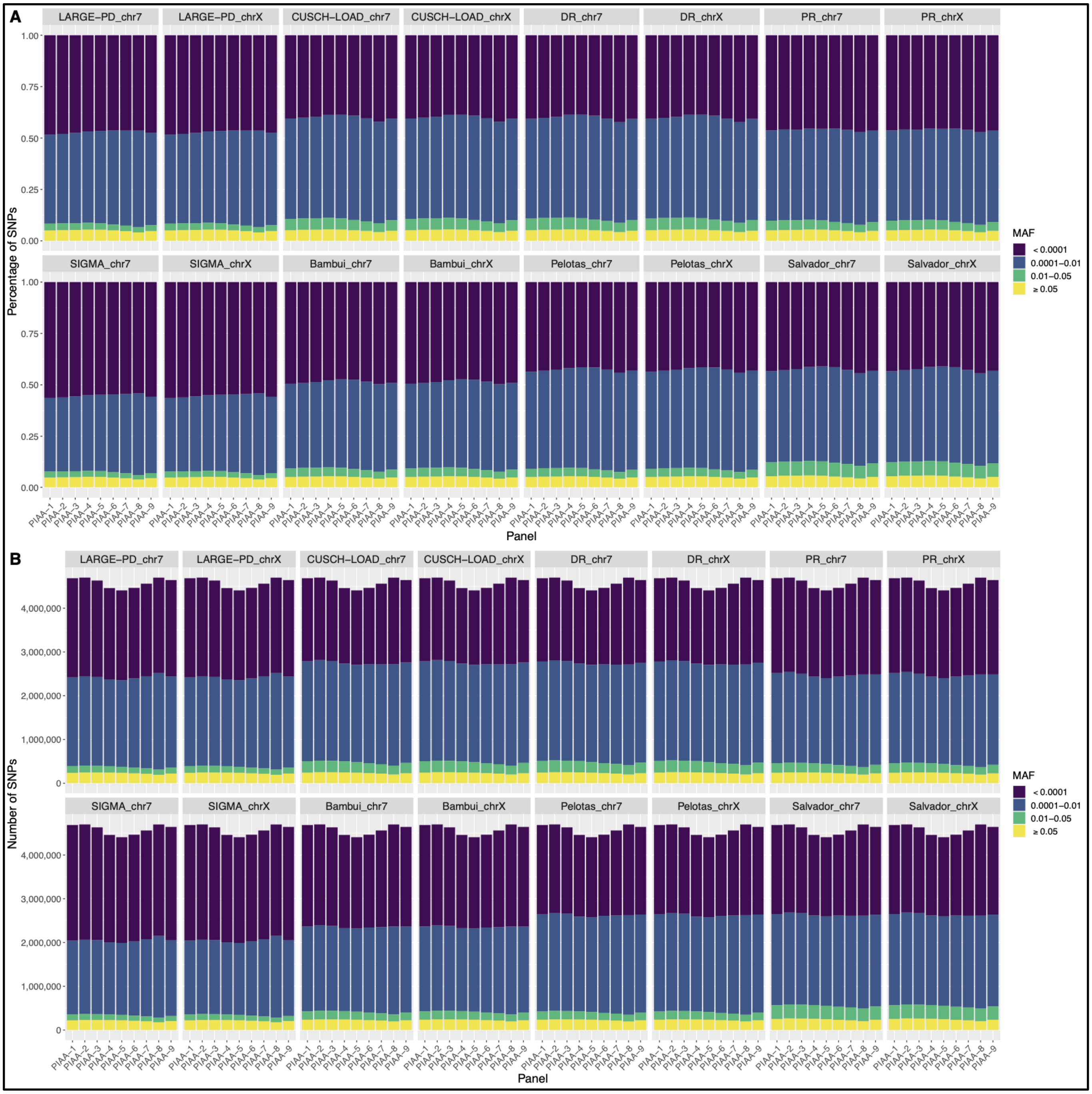


**Figure S14. Comparison of MAF of imputed SNPs in PIAA panels.** a) The percentage of SNPs in each MAF category by population and imputation reference panel. b) The number of SNPs in each MAF category by population and imputation reference panel. LARGE-PD: Latin American Research Consortium on the Genetics of Parkinson's Disease, CUSCH-LOAD: Columbia University study of Caribbean Hispanics with familial or sporadic late-onset Alzheimer's disease, DR: subset of CUSCH-LOAD individuals from the Dominican Republic, PR: subset of CUSCH-LOAD individuals from Puerto Rico, SIGMA: the Slim Initiative in genomic medicine for the Americas.

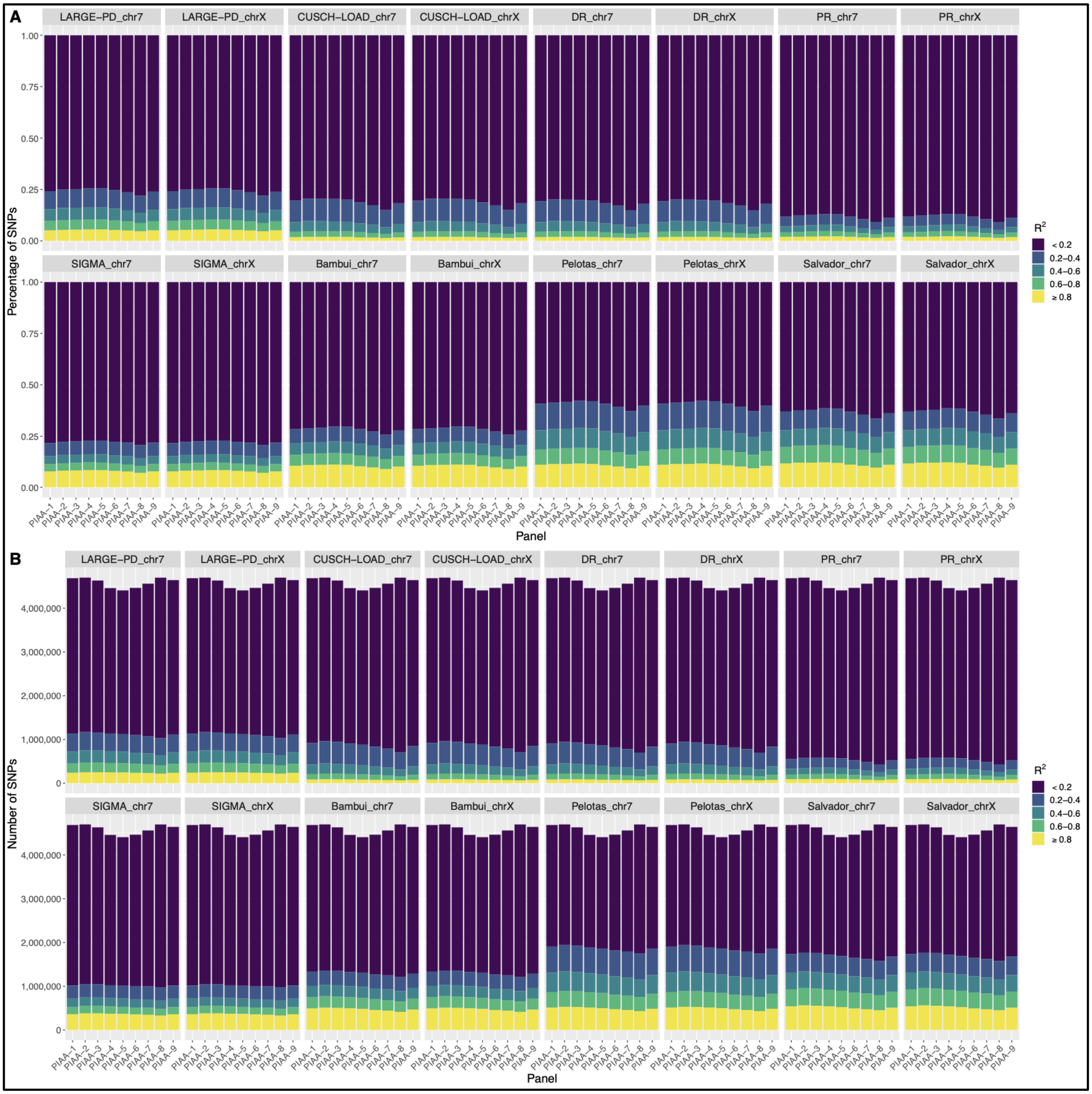


**Figure S15. Comparison of $R^2$ of imputed SNPs in PIAA panels.** a) The percentage of SNPs in each $R^2$ category by population and imputation reference panel. b) The number of SNPs in each $R^2$ category by population and imputation reference panel. LARGE-PD: Latin American Research Consortium on the Genetics of Parkinson's Disease, CUSCH-LOAD: Columbia University study of Caribbean Hispanics with familial or sporadic late-onset Alzheimer's disease, DR: subset of CUSCH-LOAD individuals from the Dominican Republic, PR: subset of

CUSCH-LOAD individuals from Puerto Rico, SIGMA: the Slim Initiative in genomic medicine for the Americas.

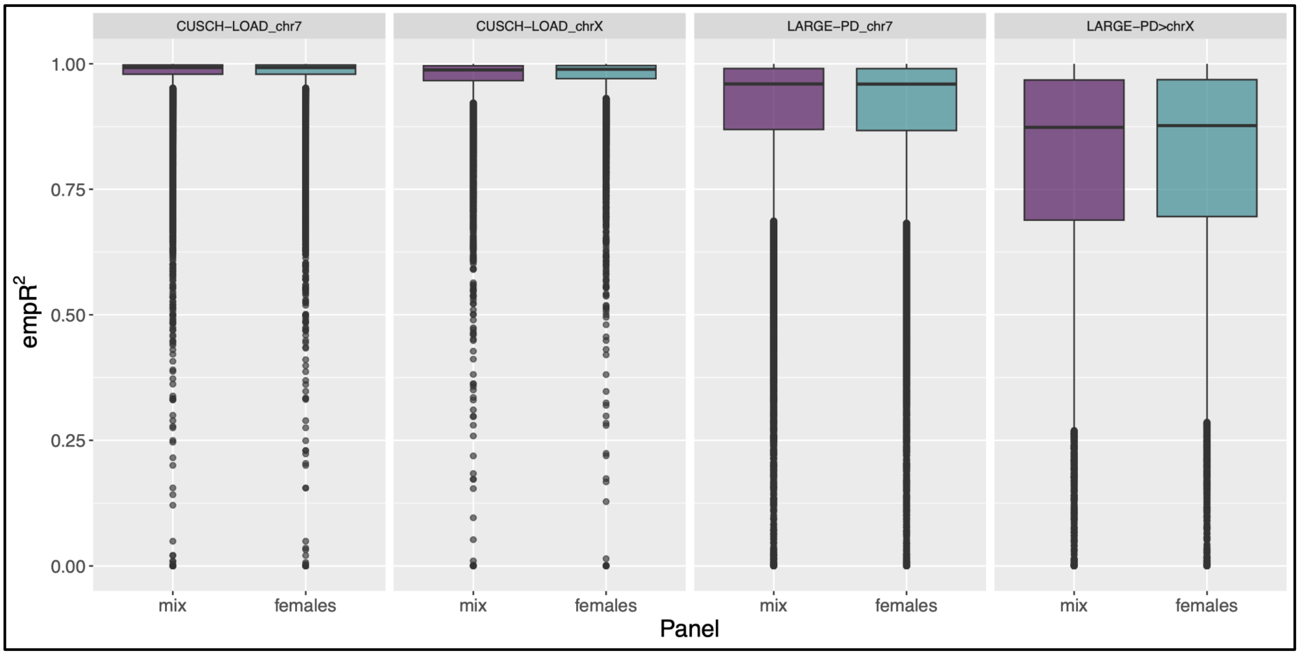


**Figure S16. Boxplot of empR$^2$ values of females and a combination of males and females.** Comparison of empirical $R^2$ values in two cohorts, after imputation using two panels - one containing both males and females and one with only females. LARGE-PD: Latin American Research Consortium on the Genetics of Parkinson's Disease, CUSCH-LOAD: Columbia University study of Caribbean Hispanics with familial or sporadic late-onset Alzheimer's disease, mix: panel contains a mix of females and males, females: panel only contains females.
