## Supplementary Tables for "Comparing the effect of imputation reference panel composition in four distinct Latin American cohorts"

**Table S1.** Cohorts included in subset of TOPMed freeze 10b used for imputatior

| **Cohort** | **Number of Individuals** |
|---|---|
| Women's Health Initiative (WHI) | 11,310 |
| Jackson Heart Study (JHS) | 3,418 |
| the 1000 Genomes Project (1KGP) | 3,199 |
| Human Genome Diversity Project (HGDP) | 827 |
| Framingham Heart Study (FHS) | 7,183 |
| Barbados Asthma Genetics Study (BAGS) | 1,084 |
| Multiethnic Study of Atherosclerosis (MESA) | 7,872 |
| The Genetic Epidemiology of Asthma in Costa Rica | 3,722 |
| San Antonio Family Heart Study (SAFHS) | 1,744 |
| Hispanic Community Health Study/Study of Latinos (HCHS/SOL) | 7,728 |
| Severe Asthma Research Program (SARP) | 185 |
| Recipient Epidemiology and Donor Evaluation Study-III Brazil Sickle Cell Disease Cohort (REDS-BSCDC) | 2,611 |
| Boston-Brazil Sickle Cell Disease (SCD) Cohort | 406 |
| Children's Health Study (CHS) Integrative Genomics and Environmental Research of Asthma (IGERA) | 153 |
| Children's Health Study (CHS) Effects of Air Pollution on the Development of Obesity in Children (Meta-AIR) | 52 |
| The BioMe Biobank at Mount Sinai | 5,267 |
| Lung Tissue Research Consortium (LTRC) | 115 |
| Childhood Asthma Management Program (CAMP) | 1,770 |
| Total individuals: | 58,646 |

ı reference panels.

| Origin |
|---|
| United States |
| United States |
| Africa, America, East Asia, South Asia, and Europe |
| Africa, West Asia, Central/South Asia, East Asia, American, Europe, and Oceania |
| United States |
| Barbados |
| United States |
| Costa Rica |
| United States |
| United States |
| United States |
| Brazil |
| Brazil |
| United States |
| United States |
| United States |
| United States |
| United States |

**Table S2. NoLA imputation reference panels.** Number of individuals from each population included in imputation reference panels for NoLA comparison.

| **Panel** | **# of Europeans** | **# of Africans** | **# of Latin Americans** |
|---|---|---|---|
| NoLA-1 | 5,500 | 5,500 | 1,000 |
| NoLA-2 | 5,000 | 5,000 | 2,000 |
| NoLA-3 | 4,500 | 4,500 | 3,000 |
| NoLA-4 | 4,000 | 4,000 | 4,000 |
| NoLA-5 | 3,500 | 3,500 | 5,000 |
| NoLA-S | 1,000 | 1,000 | 1,000 |
| NoLA-M | 2,500 | 2,500 | 2,500 |

**Table S3. LOPO imputation reference panel.** Number of individuals from each population included in imputation reference panels for LOPO comparison.

| Panel | # of Europeans | # of Africans | # of Latin Americans |
|---|---|---|---|
| LOPO-EA | 6,000 | 6,000 | 0 |
| LOPO-EL | 6,000 | 0 | 6,000 |
| LOPO-AL | 0 | 6,000 | 6,000 |
| LOPO-all | 4,000 | 4,000 | 4,000 |

**Table S4. Latin Americans in PIAA imputation reference panels.** Position of self-described Latin Americans included in each panel after ordering by percent Indigenous American ancestry.

| Panel | First individual position | Last individual position |
|---|---|---|
| PIAA-1 | 1 | 4,000 |
| PIAA-2 | 2,001 | 6,000 |
| PIAA-3 | 4,001 | 8,000 |
| PIAA-4 | 6,001 | 10,000 |
| PIAA-5 | 8,001 | 12,000 |
| PIAA-6 | 10,001 | 14,000 |
| PIAA-7 | 12,001 | 16,000 |
| PIAA-8 | 13,808 | 17,807 |
| PIAA-9 | mix | mix |

**Table S5. Admixture proprotions for target populatior**
(while excluding chromosome 7), chromosome 7, and chr

| Population | Average ancestry proportions on autosomes, except 7 |
|---|---|
| LARGE-PD | AFR: 0.0590422<br>EUR: 0.541003<br>IA: 0.399955 |
| CUSCH-LOAD | AFR: 0.337042<br>EUR: 0.574489<br>IA: 0.0884692 |
| DR | AFR: 0.367585<br>EUR: 0.556675<br>IA: 0.0757399 |
| PR | AFR: 0.176488<br>EUR: 0.688121<br>IA: 0.135391 |
| SIGMA | AFR: 0.0207229<br>EUR: 0.270198<br>IA: 0.709079 |
| Bambui | AFR: 0.162373<br>EUR: 0.763318<br>IA: 0.0743093 |
| Pelotas | AFR: 0.148656<br>EUR: 0.7728<br>IA: 0.0785431 |
| Salvador | AFR: 0.500386<br>EUR: 0.434671<br>IA: 0.0649433 |

**ns.** Average ancestry proportions for each target population based on autosomes omosome X, conducted via ADMIXTURE.

| Average ancestry proportions on chromosome 7 | Average ancestry proportions on chromosome X |
|---|---|
| AFR: 0.0564582<br>EUR: 0.518826<br>IA: 0.424715 | AFR: 0.0676172<br>EUR: 0.487022<br>IA: 0.44536 |
| AFR: 0.326542<br>EUR: 0.57946<br>IA: 0.0939976 | AFR: 0.399142<br>EUR: 0.482358<br>IA: 0.1185 |
| AFR: 0.357347<br>EUR: 0.560032<br>IA: 0.0826211 | AFR: 0.436715<br>EUR: 0.46186<br>IA: 0.101425 |
| AFR: 0.166315<br>EUR: 0.709147<br>IA: 0.124537 | AFR: 0.194805<br>EUR: 0.61553<br>IA: 0.189665 |
| AFR: 0.0197097<br>EUR: 0.220632<br>IA: 0.759658 | AFR: 0.0246009<br>EUR: 0.174601<br>IA: 0.800798 |
| AFR: 0.147034<br>EUR: 0.774672<br>IA: 0.0782941 | AFR: 0.186302<br>EUR: 0.699301<br>IA: 0.114397 |
| AFR: 0.148637<br>EUR: 0.778271<br>IA: 0.0730915 | AFR: 0.180945<br>EUR: 0.715868<br>IA: 0.103187 |
| AFR: 0.499594<br>EUR: 0.431603<br>IA: 0.0688023 | AFR: 0.604125<br>EUR: 0.314458<br>IA: 0.0808975 |

**Table S6. Pairwise comparison results for NoLA reference panels.** Medians
using panel NoLA-4 as a reference. All pairwise comparison results can be four

| | Chromosome 7 | | | | |
|---|---|---|---|---|---|
| | **Panel** | **Median** | **Mean** | **Std dev** | **Median of differences** |
| **CUSCH-LOAD** | **NoLA-1** | 0.4047 | 0.4317 | 0.3266 | -0.00240 |
| | **NoLA-2** | 0.4100 | 0.4341 | 0.3269 | -0.00128 |
| | **NoLA-3** | 0.4126 | 0.4363 | 0.3265 | -0.00042 |
| | **NoLA-4** | 0.4142 | 0.4375 | 0.3260 | - |
| | **NoLA-5** | 0.4200 | 0.4398 | 0.3258 | 0.00058 |
| | **NoLA-S** | 0.4293 | 0.4414 | 0.3217 | -0.00416 |
| | **NoLA-M** | 0.4257 | 0.4426 | 0.3242 | -0.00136 |
| **DR** | **NoLA-1** | 0.3984 | 0.4281 | 0.3258 | -0.00194 |
| | **NoLA-2** | 0.4026 | 0.4304 | 0.3261 | -0.00100 |
| | **NoLA-3** | 0.4051 | 0.4323 | 0.3258 | -0.00031 |
| | **NoLA-4** | 0.4076 | 0.4332 | 0.3252 | - |
| | **NoLA-5** | 0.4108 | 0.4354 | 0.3250 | 0.00049 |
| | **NoLA-S** | 0.4196 | 0.4359 | 0.3209 | -0.00474 |
| | **NoLA-M** | 0.4175 | 0.4380 | 0.3235 | -0.00145 |
| **PR** | **NoLA-1** | 0.4743 | 0.4718 | 0.3367 | -0.00415 |
| | **NoLA-2** | 0.4818 | 0.4744 | 0.3370 | -0.00222 |
| | **NoLA-3** | 0.4860 | 0.4779 | 0.3365 | -0.00071 |
| | **NoLA-4** | 0.4896 | 0.4798 | 0.3362 | - |
| | **NoLA-5** | 0.4913 | 0.4830 | 0.3360 | 0.00121 |
| | **NoLA-S** | 0.5028 | 0.4868 | 0.3319 | -0.00240 |
| | **NoLA-M** | 0.4985 | 0.4866 | 0.3343 | -0.00046 |
| **LARGE-PD** | **NoLA-1** | 0.4423 | 0.4606 | 0.3205 | -0.0116559 |
| | **NoLA-2** | 0.4521 | 0.4670 | 0.3203 | -0.00608 |
| | **NoLA-3** | 0.4621 | 0.4729 | 0.3199 | -0.00220 |
| | **NoLA-4** | 0.4674 | 0.4771 | 0.3196 | - |
| | **NoLA-5** | 0.4779 | 0.4829 | 0.3189 | 0.00238 |
| | **NoLA-S** | 0.4459 | 0.4615 | 0.3208 | -0.01710 |
| | **NoLA-M** | 0.4706 | 0.4773 | 0.3194 | -0.00376 |

| | | | | | |
|---|---|---|---|---|---|
| | **NoLA-1** | 0.7547 | 0.6522 | 0.3244 | -0.00845 |
| | **NoLA-2** | 0.7621 | 0.6590 | 0.3214 | -0.00427 |
| | **NoLA-3** | 0.7703 | 0.6649 | 0.3186 | -0.00154 |
| **SIGMA** | **NoLA-4** | 0.7739 | 0.6687 | 0.3169 | - |
| | **NoLA-5** | 0.7782 | 0.6730 | 0.3151 | 0.00133 |
| | **NoLA-S** | 0.7592 | 0.6591 | 0.3202 | -0.00698 |
| | **NoLA-M** | 0.7716 | 0.6691 | 0.3163 | -0.00141 |
| | **NoLA-1** | 0.7379 | 0.6567 | 0.3025 | -0.00260 |
| | **NoLA-2** | 0.7413 | 0.6590 | 0.3021 | -0.00141 |
| | **NoLA-3** | 0.7434 | 0.6607 | 0.3016 | -0.00063 |
| **Bambui** | **NoLA-4** | 0.7459 | 0.6630 | 0.3006 | - |
| | **NoLA-5** | 0.7476 | 0.6647 | 0.3001 | 0.00043 |
| | **NoLA-S** | 0.7136 | 0.6404 | 0.3052 | -0.01619 |
| | **NoLA-M** | 0.7407 | 0.6601 | 0.3008 | -0.00307 |
| | **NoLA-1** | 0.7187 | 0.6459 | 0.3001 | -0.00228 |
| | **NoLA-2** | 0.7220 | 0.6482 | 0.2994 | -0.00106 |
| | **NoLA-3** | 0.7235 | 0.6496 | 0.2989 | -0.00052 |
| **Pelotas** | **NoLA-4** | 0.7255 | 0.6513 | 0.2984 | - |
| | **NoLA-5** | 0.7272 | 0.6526 | 0.2979 | 0.00028 |
| | **NoLA-S** | 0.6926 | 0.6267 | 0.3041 | -0.01963 |
| | **NoLA-M** | 0.7201 | 0.6473 | 0.2991 | -0.00430 |
| | **NoLA-1** | 0.6855 | 0.6216 | 0.3005 | -0.00152 |
| | **NoLA-2** | 0.6875 | 0.6234 | 0.3001 | -0.00058 |
| | **NoLA-3** | 0.6882 | 0.6244 | 0.2997 | -0.00018 |
| **Salvador** | **NoLA-4** | 0.6896 | 0.6252 | 0.2992 | - |
| | **NoLA-5** | 0.6922 | 0.6266 | 0.2988 | 0.00024 |
| | **NoLA-S** | 0.6443 | 0.5932 | 0.3032 | -0.02737 |
| | **NoLA-M** | 0.6808 | 0.6187 | 0.2998 | -0.00660 |

, p-values and median of differences calculated using a paired, two-sided Wilcoxon signe
nd in Table S7.

| | | Chromosome X | | | |
|---|---|---|---|---|---|
| **p-value** | **Panel** | **Median** | **Mean** | **Std dev** | **Median of differences** |
| 3.05E-49 | **NoLA-1** | 0.5345 | 0.5178 | 0.2941 | -0.00604 |
| 3.30E-28 | **NoLA-2** | 0.5366 | 0.5208 | 0.2929 | -0.00345 |
| 3.00E-09 | **NoLA-3** | 0.5392 | 0.5229 | 0.2921 | -0.00153 |
| - | **NoLA-4** | 0.5422 | 0.5249 | 0.2912 | - |
| 3.59E-15 | **NoLA-5** | 0.5419 | 0.5257 | 0.2909 | 0.00065 |
| 1.84E-22 | **NoLA-S** | 0.5114 | 0.5013 | 0.2974 | -0.02170 |
| 4.93E-14 | **NoLA-M** | 0.5316 | 0.5164 | 0.2937 | -0.00763 |
| **-** | | | | | |
| 5.64E-32 | **NoLA-1** | 0.5176 | 0.5074 | 0.2950 | -0.00524 |
| 1.09E-17 | **NoLA-2** | 0.5205 | 0.5097 | 0.2941 | -0.00310 |
| 1.30E-04 | **NoLA-3** | 0.5220 | 0.5115 | 0.2935 | -0.00140 |
| - | **NoLA-4** | 0.5234 | 0.5134 | 0.2927 | - |
| 2.03E-10 | **NoLA-5** | 0.5238 | 0.5141 | 0.2924 | 0.00055 |
| 5.35E-28 | **NoLA-S** | 0.4949 | 0.4903 | 0.2983 | -0.02126 |
| 1.96E-15 | **NoLA-M** | 0.5123 | 0.5051 | 0.2950 | -0.00753 |
| **-** | | | | | |
| 7.50E-64 | **NoLA-1** | 0.6542 | 0.5965 | 0.3009 | -0.01134 |
| 6.86E-36 | **NoLA-2** | 0.6613 | 0.6045 | 0.2974 | -0.00534 |
| 1.30E-09 | **NoLA-3** | 0.6670 | 0.6099 | 0.2952 | -0.00145 |
| - | **NoLA-4** | 0.6707 | 0.6122 | 0.2944 | - |
| 8.28E-27 | **NoLA-5** | 0.6717 | 0.6145 | 0.2938 | 0.00122 |
| 6.05E-06 | **NoLA-S** | 0.6374 | 0.5852 | 0.3037 | -0.02129 |
| 0.86570316 | **NoLA-M** | 0.6620 | 0.6045 | 0.2971 | -0.00591 |
| **-** | | | | | |
| < 4.68E-307 | **NoLA-1** | 0.6501 | 0.6071 | 0.2850 | -0.00925 |
| 6.38E-292 | **NoLA-2** | 0.6529 | 0.6100 | 0.2840 | -0.00625 |
| 5.85E-107 | **NoLA-3** | 0.6589 | 0.6154 | 0.2819 | -0.00200 |
| - | **NoLA-4** | 0.6637 | 0.6185 | 0.2807 | - |
| 6.55E-133 | **NoLA-5** | 0.6642 | 0.6195 | 0.2799 | 0.00063 |
| < 4.68E-307 | **NoLA-S** | 0.5890 | 0.5612 | 0.2974 | -0.04428 |
| 7.07E-77 | **NoLA-M** | 0.6358 | 0.5983 | 0.2865 | -0.01408 |

| | | | | | |
|---|---|---|---|---|---|
| **-** | | | | | |
| < 4.68E-307 | **NoLA-1** | 0.9103 | 0.7885 | 0.2625 | -0.00450 |
| < 4.68E-307 | **NoLA-2** | 0.9137 | 0.7925 | 0.2597 | -0.00243 |
| 4.02E-206 | **NoLA-3** | 0.9156 | 0.7956 | 0.2572 | -0.00098 |
| - | **NoLA-4** | 0.9178 | 0.7979 | 0.2556 | - |
| 7.56E-171 | **NoLA-5** | 0.9190 | 0.7995 | 0.2545 | 0.00063 |
| 1.73E-237 | **NoLA-S** | 0.9051 | 0.7828 | 0.2652 | -0.00666 |
| 6.05E-53 | **NoLA-M** | 0.9141 | 0.7930 | 0.2589 | -0.00190 |
| **-** | | | | | |
| 1.51E-125 | **NoLA-1** | 0.8518 | 0.7582 | 0.2515 | -0.00189 |
| 5.28E-65 | **NoLA-2** | 0.8532 | 0.7596 | 0.2506 | -0.00100 |
| 4.26E-33 | **NoLA-3** | 0.8545 | 0.7611 | 0.2494 | -0.00032 |
| - | **NoLA-4** | 0.8550 | 0.7617 | 0.2493 | - |
| 2.86E-17 | **NoLA-5** | 0.8549 | 0.7622 | 0.2489 | 0.00015 |
| < 4.68E-307 | **NoLA-S** | 0.8288 | 0.7380 | 0.2596 | -0.01419 |
| 1.93E-143 | **NoLA-M** | 0.8473 | 0.7548 | 0.2523 | -0.00374 |
| **-** | | | | | |
| 2.65E-143 | **NoLA-1** | 0.8353 | 0.7564 | 0.2375 | -0.00160 |
| 1.27E-57 | **NoLA-2** | 0.8360 | 0.7574 | 0.2365 | -0.00087 |
| 5.36E-31 | **NoLA-3** | 0.8371 | 0.7585 | 0.2357 | -0.00021 |
| - | **NoLA-4** | 0.8364 | 0.7588 | 0.2355 | - |
| 4.50E-10 | **NoLA-5** | 0.8364 | 0.7589 | 0.2354 | 0.00012 |
| < 4.68E-307 | **NoLA-S** | 0.8093 | 0.7317 | 0.2496 | -0.01814 |
| < 4.68E-307 | **NoLA-M** | 0.8283 | 0.7504 | 0.2399 | -0.00496 |
| **-** | | | | | |
| 2.39E-43 | **NoLA-1** | 0.7759 | 0.7208 | 0.2357 | -0.00077 |
| 2.23E-10 | **NoLA-2** | 0.7758 | 0.7214 | 0.2350 | -0.00035 |
| 0.0143174 | **NoLA-3** | 0.7765 | 0.7222 | 0.2344 | 0.00001 |
| - | **NoLA-4** | 0.7767 | 0.7222 | 0.2342 | - |
| 1.07E-03 | **NoLA-5** | 0.7769 | 0.7220 | 0.2342 | -0.00007 |
| < 4.68E-307 | **NoLA-S** | 0.7427 | 0.6926 | 0.2451 | -0.02261 |
| < 4.68E-307 | **NoLA-M** | 0.7665 | 0.7132 | 0.2376 | -0.00620 |

d rank test,

| p-value |
|---|
| < 4.68E-307 |
| < 4.68E-307 |
| 5.46E-175 |
| - |
| 4.15E-37 |
| < 4.68E-307 |
| < 4.68E-307 |
| **-** |
| < 4.68E-307 |
| 4.69E-276 |
| 6.34E-141 |
| - |
| 8.15E-23 |
| < 4.68E-307 |
| < 4.68E-307 |
| **-** |
| < 4.68E-307 |
| 1.37E-291 |
| 9.66E-78 |
| - |
| 2.96E-52 |
| < 4.68E-307 |
| < 4.68E-307 |
| **-** |
| 1.03E-271 |
| 3.42E-292 |
| 2.17E-99 |
| - |
| 9.73E-12 |
| < 4.68E-307 |
| < 4.68E-307 |

| |
|---|
| - |
| < 4.68E-307 |
| < 4.68E-307 |
| < 4.68E-307 |
| - |
| 1.47E-227 |
| < 4.68E-307 |
| < 4.68E-307 |
| - |
| 2.14E-201 |
| 1.47E-126 |
| 3.64E-39 |
| - |
| 4.03E-11 |
| < 4.68E-307 |
| < 4.68E-307 |
| - |
| 7.56E-196 |
| 1.30E-120 |
| 1.17E-23 |
| - |
| 2.81E-05 |
| < 4.68E-307 |
| < 4.68E-307 |
| - |
| 1.97E-34 |
| 1.59E-12 |
| > 0.99 |
| - |
| 0.09815883 |
| < 4.68E-307 |
| < 4.68E-307 |

**Table S7. Full pairwise comparison results for NoLA reference panels.** Medians, rank test. P-values are found in the lower triangle of the comparison, while the media is column - row.

**Population** **CUS(**

| **Panel** | **Median** | | **NoLA-1** | **NoLA-2** |
|---|---|---|---|---|
| NoLA-1 | 0.4047 | NoLA-1 | - | 0.000947658 |
| NoLA-2 | 0.40997 | NoLA-2 | 2.01E-45 | - |
| NoLA-3 | 0.41257 | NoLA-3 | 8.84E-60 | 2.75E-27 |
| NoLA-4 | 0.41417 | NoLA-4 | 3.05E-49 | 3.30E-28 |
| NoLA-5 | 0.41997 | NoLA-5 | 8.08E-60 | 9.61E-42 |
| NoLA-S | 0.42929 | NoLA-S | 2.81E-01 | 1.79E-06 |
| NoLA-M | 0.42565 | NoLA-M | 1.30E-08 | > 0.99 |

| **Panel** | **Median** | | **NoLA-1** | **NoLA-2** |
|---|---|---|---|---|
| NoLA-1 | 0.39836 | NoLA-1 | - | 0.000804839 |
| NoLA-2 | 0.40256 | NoLA-2 | 2.41E-31 | - |
| NoLA-3 | 0.40513 | NoLA-3 | 1.60E-39 | 4.89E-17 |
| NoLA-4 | 0.40757 | NoLA-4 | 5.64E-32 | 1.09E-17 |
| NoLA-5 | 0.41076 | NoLA-5 | 2.74E-42 | 1.88E-27 |
| NoLA-S | 0.41958 | NoLA-S | 8.47E-05 | 3.56E-11 |
| NoLA-M | 0.41747 | NoLA-M | 1.58E-03 | > 0.99 |

| **Panel** | **Median** | | **NoLA-1** | **NoLA-2** |
|---|---|---|---|---|
| NoLA-1 | 0.47432 | NoLA-1 | - | 0.001625584 |
| NoLA-2 | 0.48177 | NoLA-2 | 2.33E-44 | - |
| NoLA-3 | 0.48596 | NoLA-3 | 3.81E-69 | 6.45E-30 |
| NoLA-4 | 0.48957 | NoLA-4 | 7.50E-64 | 6.86E-36 |
| NoLA-5 | 0.49132 | NoLA-5 | 4.23E-84 | 1.84E-53 |
| NoLA-S | 0.50277 | NoLA-S | 3.25E-04 | > 0.99 |
| NoLA-M | 0.49853 | NoLA-M | 1.09E-40 | 2.26E-16 |

**LA**

| **Panel** | **Median** | | **NoLA-1** | **NoLA-2** |
|---|---|---|---|---|
| NoLA-1 | 0.44231 | NoLA-1 | - | 0.004463189 |

| | | | | |
|---|---|---|---|---|
| NoLA-2 | 0.45211 | NoLA-2 | < 4.68E-307 | - |
| NoLA-3 | 0.46207 | NoLA-3 | < 4.68E-307 | 1.244E-214 |
| NoLA-4 | 0.46742 | NoLA-4 | < 4.68E-307 | 6.3814E-292 |
| NoLA-5 | 0.47791 | NoLA-5 | < 4.68E-307 | < 4.68E-307 |
| NoLA-S | 0.44586 | NoLA-S | 1.05215E-25 | 6.5299E-112 |
| NoLA-M | 0.47056 | NoLA-M | 1.3087E-177 | 1.65568E-26 |

**S[illegible]**

| **Panel** | **Median** | | **NoLA-1** | **NoLA-2** |
|---|---|---|---|---|
| NoLA-1 | 0.75466 | NoLA-1 | - | 0.003141615 |
| NoLA-2 | 0.76206 | NoLA-2 | < 4.68E-307 | - |
| NoLA-3 | 0.77032 | NoLA-3 | < 4.68E-307 | < 4.68E-307 |
| NoLA-4 | 0.77391 | NoLA-4 | < 4.68E-307 | < 4.68E-307 |
| NoLA-5 | 0.77818 | NoLA-5 | < 4.68E-307 | < 4.68E-307 |
| NoLA-S | 0.7592 | NoLA-S | 2.22198E-07 | 7.68794E-26 |
| NoLA-M | 0.77155 | NoLA-M | < 4.68E-307 | 9.3139E-116 |

**B**

| **Panel** | **Median** | | **NoLA-1** | **NoLA-2** |
|---|---|---|---|---|
| NoLA-1 | 0.73785 | NoLA-1 | - | 0.000827112 |
| NoLA-2 | 0.7413 | NoLA-2 | 6.68E-60 | - |
| NoLA-3 | 0.74343 | NoLA-3 | 8.70E-83 | 3.89E-32 |
| NoLA-4 | 0.7459 | NoLA-4 | 1.51E-125 | 5.28E-65 |
| NoLA-5 | 0.74756 | NoLA-5 | 2.95E-144 | 2.69E-83 |
| NoLA-S | 0.71355 | NoLA-S | < 4.68E-307 | < 4.68E-307 |
| NoLA-M | 0.74067 | NoLA-M | > 0.99 | 2.08E-27 |

**P**

| **Panel** | **Median** | | **NoLA-1** | **NoLA-2** |
|---|---|---|---|---|
| NoLA-1 | 0.71872 | NoLA-1 | - | 0.000871061 |
| NoLA-2 | 0.722 | NoLA-2 | 1.16E-94 | - |
| NoLA-3 | 0.72354 | NoLA-3 | 4.54E-111 | 6.60E-32 |
| NoLA-4 | 0.725505 | NoLA-4 | 2.65E-143 | 1.27E-57 |
| NoLA-5 | 0.72715 | NoLA-5 | 2.85E-149 | 5.83E-73 |
| NoLA-S | 0.692645 | NoLA-S | < 4.68E-307 | < 4.68E-307 |
| NoLA-M | 0.72011 | NoLA-M | 8.31E-27 | 2.85E-121 |

| | | | | Sa |
|---|---|---|---|---|
| **Panel** | **Median** | | **NoLA-1** | **NoLA-2** |
| NoLA-1 | 0.685525 | NoLA-1 | - | 0.000642972 |
| NoLA-2 | 0.68753 | NoLA-2 | 5.78E-32 | - |
| NoLA-3 | 0.68819 | NoLA-3 | 1.06E-39 | 2.13E-07 |
| NoLA-4 | 0.68957 | NoLA-4 | 2.39E-43 | 2.23E-10 |
| NoLA-5 | 0.692235 | NoLA-5 | 3.69E-57 | 7.19E-22 |
| NoLA-S | 0.64427 | NoLA-S | < 4.68E-307 | < 4.68E-307 |
| NoLA-M | 0.68078 | NoLA-M | 3.18E-159 | < 4.68E-307 |

p-values and median of differences calculated using a paired, two-sided Wilcoxo
n of differences are in the upper triangle of the table (above the "-"). The median

**:H-LOAD_chr7**

| NoLA-3 | NoLA-4 | NoLA-5 | NoLA-S | NoLA-M |
|---|---|---|---|---|
| 0.001918923 | 0.002399005 | 0.003393727 | -0.001034052 | 0.0014583 |
| 0.000749004 | 0.001270951 | 0.002230723 | -0.002236156 | 0.0002357 |
| - | 0.000418784 | 0.001185556 | -0.003361354 | -0.000797 |
| 3.00E-09 | - | 0.000580167 | -0.004163542 | -0.001355 |
| 9.24E-23 | 3.59E-15 | - | -0.005408975 | -0.002289 |
| 4.38E-15 | 1.84E-22 | 1.32E-38 | - | 0.0051133 |
| 1.29E-04 | 4.93E-14 | 4.29E-43 | 1.79E-68 | - |

**DR_chr7**

| NoLA-3 | NoLA-4 | NoLA-5 | NoLA-S | NoLA-M |
|---|---|---|---|---|
| 0.001534726 | 0.001936631 | 0.002823579 | -0.001986503 | 0.0009188 |
| 0.00060833 | 0.000998777 | 0.001829624 | -0.002987563 | -0.000113 |
| - | 0.000309585 | 0.000966558 | -0.004054583 | -0.000988 |
| 1.30E-04 | - | 0.000485893 | -0.004740234 | -0.001451 |
| 9.74E-15 | 2.03E-10 | - | -0.005850325 | -0.00225 |
| 3.59E-21 | 5.35E-28 | 4.91E-44 | - | 0.0053 |
| 1.22E-06 | 1.96E-15 | 2.57E-41 | 1.40E-72 | - |

**PR_chr7**

| NoLA-3 | NoLA-4 | NoLA-5 | NoLA-S | NoLA-M |
|---|---|---|---|---|
| 0.003332261 | 0.004153755 | 0.00593369 | 0.002197291 | 0.0045023 |
| 0.001331011 | 0.002219484 | 0.003755677 | 0.000322635 | 0.0023547 |
| - | 0.000710386 | 0.002012115 | -0.001371799 | 0.000568 |
| 1.30E-09 | - | 0.001211886 | -0.002401167 | -0.000461 |
| 2.17E-28 | 8.28E-27 | - | -0.004203683 | -0.00179 |
| 6.18E-02 | 6.05E-06 | 1.79E-17 | - | 0.0040429 |
| 3.25E-01 | 8.66E-01 | 1.03E-14 | 2.82E-30 | - |

**RGE-PD_chr7**

| NoLA-3 | NoLA-4 | NoLA-5 | NoLA-S | NoLA-M |
|---|---|---|---|---|
| 0.008524568 | 0.01165592 | 0.01556019 | -0.004256322 | 0.0081639 |

| | | | | |
|---|---|---|---|---|
| 0.003263655 | 0.006084406 | 0.009659008 | -0.009526438 | 0.002419 |
| - | 0.002203391 | 0.005267915 | -0.01386543 | -0.001242 |
| 5.8535E-107 | - | 0.002382832 | -0.01709539 | -0.003765 |
| 1.1332E-243 | 6.551E-133 | - | -0.02128115 | -0.006926 |
| 7.102E-219 | < 4.68E-307 | < 4.68E-307 | - | 0.0143927 |
| 9.14889E-10 | 7.06552E-77 | 5.6326E-248 | < 4.68E-307 | - |

**IGMA_chr7**

| NoLA-3 | NoLA-4 | NoLA-5 | NoLA-S | NoLA-M |
|---|---|---|---|---|
| 0.006130727 | 0.008447366 | 0.01070432 | 0.001003759 | 0.0066635 |
| 0.002246857 | 0.004267021 | 0.006285639 | -0.001966625 | 0.0026225 |
| - | 0.001541539 | 0.003231483 | -0.004810248 | 3.93E-05 |
| 4.0168E-206 | - | 0.001329124 | -0.006981204 | -0.001408 |
| < 4.68E-307 | 7.5575E-171 | - | -0.009219798 | -0.00306 |
| 1.525E-126 | 1.7298E-237 | < 4.68E-307 | - | 0.0057132 |
| 9.1825062 | 6.05371E-53 | 3.0249E-214 | < 4.68E-307 | - |

**ambui_chr7**

| NoLA-3 | NoLA-4 | NoLA-5 | NoLA-S | NoLA-M |
|---|---|---|---|---|
| 0.0015583 | 0.00259631 | 0.00346704 | -0.0122945 | -0.000154 |
| 0.000584329 | 0.001406414 | 0.002121336 | -0.01385306 | -0.001423 |
| - | 0.000625998 | 0.001307227 | -0.0149645 | -0.002261 |
| 4.26E-33 | - | 0.0004256 | -0.01619252 | -0.00307 |
| 6.89E-55 | 2.86E-17 | - | -0.01722285 | -0.00384 |
| < 4.68E-307 | < 4.68E-307 | < 4.68E-307 | - | 0.0125498 |
| 1.23E-76 | 1.93E-143 | 1.10E-248 | < 4.68E-307 | - |

**'elotas_chr7**

| NoLA-3 | NoLA-4 | NoLA-5 | NoLA-S | NoLA-M |
|---|---|---|---|---|
| 0.001497158 | 0.002284419 | 0.002884571 | -0.01597866 | -0.001374 |
| 0.000503819 | 0.00106256 | 0.001614099 | -0.01767961 | -0.002733 |
| - | 0.000518434 | 0.000932607 | -0.01859718 | -0.00347 |
| 5.36E-31 | - | 0.000284085 | -0.01963079 | -0.004304 |
| 3.59E-44 | 4.50E-10 | - | -0.02033077 | -0.004833 |
| < 4.68E-307 | < 4.68E-307 | < 4.68E-307 | - | 0.0150225 |
| 2.49E-216 | < 4.68E-307 | < 4.68E-307 | < 4.68E-307 | - |

**lvador_chr7**

| NoLA-3 | NoLA-4 | NoLA-5 | NoLA-S | NoLA-M |
|---|---|---|---|---|
| 0.001089433 | 0.001522955 | 0.002126468 | -0.02474893 | -0.004363 |
| 0.000307842 | 0.000576913 | 0.001091767 | -0.02600408 | -0.005585 |
| - | 0.000182531 | 0.000553473 | -0.02670968 | -0.006164 |
| 1.43E-02 | - | 0.000236156 | -0.0273668 | -0.006603 |
| 1.27E-09 | 1.07E-03 | - | -0.02798758 | -0.006969 |
| < 4.68E-307 | < 4.68E-307 | < 4.68E-307 | - | 0.0201708 |
| < 4.68E-307 | < 4.68E-307 | < 4.68E-307 | < 4.68E-307 | - |

on signed
difference

**CU**

| Panel | Median | | NoLA-1 | NoLA-2 |
|---|---|---|---|---|
| NoLA-1 | 0.53448 | NoLA-1 | - | 0.002412292 |
| NoLA-2 | 0.53663 | NoLA-2 | 2.34E-222 | - |
| NoLA-3 | 0.53917 | NoLA-3 | < 4.68E-307 | 8.66E-259 |
| NoLA-4 | 0.542155 | NoLA-4 | < 4.68E-307 | < 4.68E-307 |
| NoLA-5 | 0.54185 | NoLA-5 | < 4.68E-307 | 6.42E-291 |
| NoLA-S | 0.511375 | NoLA-S | < 4.68E-307 | < 4.68E-307 |
| NoLA-M | 0.53155 | NoLA-M | 1.17E-09 | 4.70E-235 |

| Panel | Median | | NoLA-1 | NoLA-2 |
|---|---|---|---|---|
| NoLA-1 | 0.51764 | NoLA-1 | - | 0.002011204 |
| NoLA-2 | 0.52053 | NoLA-2 | 7.46E-160 | - |
| NoLA-3 | 0.52201 | NoLA-3 | 3.62E-272 | 4.28E-185 |
| NoLA-4 | 0.52339 | NoLA-4 | < 4.68E-307 | 4.69E-276 |
| NoLA-5 | 0.52379 | NoLA-5 | 5.07E-290 | 9.80E-230 |
| NoLA-S | 0.49491 | NoLA-S | < 4.68E-307 | < 4.68E-307 |
| NoLA-M | 0.51226 | NoLA-M | 1.33E-26 | 3.85E-254 |

| Panel | Median | | NoLA-1 | NoLA-2 |
|---|---|---|---|---|
| NoLA-1 | 0.65418 | NoLA-1 | - | 0.005100367 |
| NoLA-2 | 0.66132 | NoLA-2 | 1.08E-289 | - |
| NoLA-3 | 0.66698 | NoLA-3 | < 4.68E-307 | 3.74E-273 |
| NoLA-4 | 0.67074 | NoLA-4 | < 4.68E-307 | 1.37E-291 |
| NoLA-5 | 0.67171 | NoLA-5 | < 4.68E-307 | 8.80E-302 |
| NoLA-S | 0.63738 | NoLA-S | 9.37E-177 | < 4.68E-307 |
| NoLA-M | 0.66199 | NoLA-M | 2.33E-109 | > 0.99 |

**I**

| Panel | Median | | NoLA-1 | NoLA-2 |
|---|---|---|---|---|
| NoLA-1 | 0.650145 | NoLA-1 | - | 0.002520698 |

| | | | | |
|---|---|---|---|---|
| NoLA-2 | 0.65294 | NoLA-2 | 1.64E-84 | - |
| NoLA-3 | 0.65892 | NoLA-3 | 4.24E-247 | 4.01E-277 |
| NoLA-4 | 0.663735 | NoLA-4 | 1.03E-271 | 3.42E-292 |
| NoLA-5 | 0.664205 | NoLA-5 | 1.07E-248 | 2.22E-233 |
| NoLA-S | 0.588975 | NoLA-S | < 4.68E-307 | < 4.68E-307 |
| NoLA-M | 0.635845 | NoLA-M | 7.79E-35 | 2.55E-190 |

| **Panel** | **Median** | | **NoLA-1** | **NoLA-2** |
|---|---|---|---|---|
| NoLA-1 | 0.91025 | NoLA-1 | - | 0.001648261 |
| NoLA-2 | 0.91371 | NoLA-2 | < 4.68E-307 | - |
| NoLA-3 | 0.91563 | NoLA-3 | < 4.68E-307 | < 4.68E-307 |
| NoLA-4 | 0.91776 | NoLA-4 | < 4.68E-307 | < 4.68E-307 |
| NoLA-5 | 0.919 | NoLA-5 | < 4.68E-307 | < 4.68E-307 |
| NoLA-S | 0.90513 | NoLA-S | 4.30054E-70 | < 4.68E-307 |
| NoLA-M | 0.91411 | NoLA-M | 1.3414E-246 | 3.23239E-34 |

| **Panel** | **Median** | | **NoLA-1** | **NoLA-2** |
|---|---|---|---|---|
| NoLA-1 | 0.8518 | NoLA-1 | - | 0.000674374 |
| NoLA-2 | 0.8532 | NoLA-2 | 2.90E-105 | - |
| NoLA-3 | 0.85452 | NoLA-3 | 1.27E-186 | 9.85E-115 |
| NoLA-4 | 0.85496 | NoLA-4 | 2.14E-201 | 1.47E-126 |
| NoLA-5 | 0.854935 | NoLA-5 | 2.47E-196 | 2.21E-120 |
| NoLA-S | 0.828775 | NoLA-S | < 4.68E-307 | < 4.68E-307 |
| NoLA-M | 0.84725 | NoLA-M | 6.18E-83 | 2.71E-283 |

| **Panel** | **Median** | | **NoLA-1** | **NoLA-2** |
|---|---|---|---|---|
| NoLA-1 | 0.83531 | NoLA-1 | - | 0.000669469 |
| NoLA-2 | 0.836 | NoLA-2 | 3.49E-120 | - |
| NoLA-3 | 0.83709 | NoLA-3 | 8.50E-208 | 1.64E-136 |
| NoLA-4 | 0.8364 | NoLA-4 | 7.56E-196 | 1.30E-120 |
| NoLA-5 | 0.83643 | NoLA-5 | 5.00E-161 | 6.37E-88 |
| NoLA-S | 0.80931 | NoLA-S | < 4.68E-307 | < 4.68E-307 |
| NoLA-M | 0.82826 | NoLA-M | < 4.68E-307 | < 4.68E-307 |

| **Panel** | **Median** | | **NoLA-1** | **NoLA-2** |
|---|---|---|---|---|
| NoLA-1 | 0.775905 | NoLA-1 | - | 0.000357482 |
| NoLA-2 | 0.77576 | NoLA-2 | 3.07E-24 | - |
| NoLA-3 | 0.7765 | NoLA-3 | 6.93E-45 | 6.12E-27 |
| NoLA-4 | 0.776745 | NoLA-4 | 1.97E-34 | 1.59E-12 |
| NoLA-5 | 0.776895 | NoLA-5 | 7.27E-21 | 2.06E-05 |
| NoLA-S | 0.74274 | NoLA-S | < 4.68E-307 | < 4.68E-307 |
| NoLA-M | 0.76646 | NoLA-M | < 4.68E-307 | < 4.68E-307 |

**JSCH-LOAD_chrX**

| NoLA-3 | NoLA-4 | NoLA-5 | NoLA-S | NoLA-M |
|---|---|---|---|---|
| 0.0042748 | 0.006036008 | 0.006893241 | -0.01479068 | -0.000967075 |
| 0.001827601 | 0.003450772 | 0.004193238 | -0.01773778 | -0.003704027 |
| - | 0.001526845 | 0.002322499 | -0.01992656 | -0.005824736 |
| 5.46E-175 | - | 0.000653394 | -0.02170156 | -0.007625586 |
| 5.75E-153 | 4.15E-37 | - | -0.02231224 | -0.008274502 |
| < 4.68E-307 | < 4.68E-307 | < 4.68E-307 | - | 0.013559 |
| < 4.68E-307 | < 4.68E-307 | < 4.68E-307 | < 4.68E-307 | - |

**DR_chrX**

| NoLA-3 | NoLA-4 | NoLA-5 | NoLA-S | NoLA-M |
|---|---|---|---|---|
| 0.003570198 | 0.005243662 | 0.006057768 | -0.01533344 | -0.001675009 |
| 0.001546754 | 0.003100602 | 0.003749659 | -0.01773388 | -0.0039857 |
| - | 0.001395484 | 0.002099001 | -0.01957443 | -0.005832298 |
| 6.34E-141 | - | 0.000547196 | -0.02125503 | -0.007531798 |
| 5.85E-117 | 8.15E-23 | - | -0.02172302 | -0.00803243 |
| < 4.68E-307 | < 4.68E-307 | < 4.68E-307 | - | 0.01321299 |
| < 4.68E-307 | < 4.68E-307 | < 4.68E-307 | < 4.68E-307 | - |

**PR_chrX**

| NoLA-3 | NoLA-4 | NoLA-5 | NoLA-S | NoLA-M |
|---|---|---|---|---|
| 0.009272904 | 0.01133557 | 0.0133006 | -0.008260199 | 0.005585963 |
| 0.003407447 | 0.005336699 | 0.007078482 | -0.01482441 | -0.000140244 |
| - | 0.001447755 | 0.003030369 | -0.01934424 | -0.003925745 |
| 9.66E-78 | - | 0.001217347 | -0.02128557 | -0.00591252 |
| 1.01E-117 | 2.96E-52 | - | -0.02294115 | -0.007318201 |
| < 4.68E-307 | < 4.68E-307 | < 4.68E-307 | - | 0.01447526 |
| 1.17E-155 | < 4.68E-307 | < 4.68E-307 | < 4.68E-307 | - |

**ARGE-PD_chrX**

| NoLA-3 | NoLA-4 | NoLA-5 | NoLA-S | NoLA-M |
|---|---|---|---|---|
| 0.00667833 | 0.009253659 | 0.01045424 | -0.0320806 | -0.00365928 |

| | | | | |
|---|---|---|---|---|
| 0.003695234 | 0.006253347 | 0.007213005 | -0.03578568 | -0.006511882 |
| - | 0.002001432 | 0.003102992 | -0.04114059 | -0.0108701 |
| 2.17E-99 | - | 0.000630425 | -0.04427958 | -0.01407889 |
| 1.38E-84 | 9.73E-12 | - | -0.04539155 | -0.01510287 |
| < 4.68E-307 | < 4.68E-307 | < 4.68E-307 | - | 0.02668071 |
| < 4.68E-307 | < 4.68E-307 | < 4.68E-307 | < 4.68E-307 | - |

**SIGMA_chrX**

| NoLA-3 | NoLA-4 | NoLA-5 | NoLA-S | NoLA-M |
|---|---|---|---|---|
| 0.003206635 | 0.004496623 | 0.00530956 | -0.001099833 | 0.002107432 |
| 0.001303857 | 0.002432764 | 0.003216789 | -0.003214964 | 0.000421858 |
| - | 0.000979229 | 0.001754494 | -0.005019813 | -0.000659844 |
| < 4.68E-307 | - | 0.000634303 | -0.006662048 | -0.00190356 |
| < 4.68E-307 | 1.4669E-227 | - | -0.007693364 | -0.002797073 |
| < 4.68E-307 | < 4.68E-307 | < 4.68E-307 | - | 0.003983817 |
| 1.9733E-106 | < 4.68E-307 | < 4.68E-307 | < 4.68E-307 | - |

**Bambui_chrX**

| NoLA-3 | NoLA-4 | NoLA-5 | NoLA-S | NoLA-M |
|---|---|---|---|---|
| 0.001386772 | 0.00188851 | 0.002214783 | -0.01148762 | -0.00147088 |
| 0.000603651 | 0.001000917 | 0.001284942 | -0.01254757 | -0.002319503 |
| - | 0.000324928 | 0.000596421 | -0.01359312 | -0.003223628 |
| 3.64E-39 | - | 0.000147416 | -0.01418517 | -0.003742143 |
| 3.12E-48 | 4.03E-11 | - | -0.0145454 | -0.004000414 |
| < 4.68E-307 | < 4.68E-307 | < 4.68E-307 | - | 0.009212458 |
| < 4.68E-307 | < 4.68E-307 | < 4.68E-307 | < 4.68E-307 | - |

**Pelotas_chrX**

| NoLA-3 | NoLA-4 | NoLA-5 | NoLA-S | NoLA-M |
|---|---|---|---|---|
| 0.001325472 | 0.00160373 | 0.001723763 | -0.01586737 | -0.002808064 |
| 0.000588157 | 0.0008843 | 0.000970486 | -0.01684173 | -0.003704824 |
| - | 0.000211619 | 0.000379545 | -0.01776025 | -0.004597682 |
| 1.17E-23 | - | 0.000116699 | -0.01813745 | -0.004956366 |
| 1.97E-22 | 2.81E-05 | - | -0.01817715 | -0.005050913 |
| < 4.68E-307 | < 4.68E-307 | < 4.68E-307 | - | 0.01181029 |
| < 4.68E-307 | < 4.68E-307 | < 4.68E-307 | < 4.68E-307 | - |

**Salvador_chrX**

| NoLA-3 | NoLA-4 | NoLA-5 | NoLA-S | NoLA-M |
|---|---|---|---|---|
| 0.000709495 | 0.000772566 | 0.000696904 | -0.02164687 | -0.005223957 |
| 0.000313514 | 3.48E-04 | 0.000291997 | -0.02193952 | -0.005608987 |
| - | -9.38E-06 | -7.98E-05 | -0.02254113 | -0.006183611 |
| > 0.99 | - | -6.72E-05 | -0.02260586 | -0.006199734 |
| 8.79E-01 | 9.82E-02 | - | -0.02245506 | -0.006035567 |
| < 4.68E-307 | < 4.68E-307 | < 4.68E-307 | - | 0.01516446 |
| < 4.68E-307 | < 4.68E-307 | < 4.68E-307 | < 4.68E-307 | - |

**Table S8. Pairwise comparison results for LOPO reference panels.** P-values and med
reference. The final column for each chromsome contains the number of replicates in whi

| | | Chromosome 7 | | | | |
|---|---|---|---|---|---|---|
| | **Panel** | **Median** | **Mean** | **Std dev** | **Median of differences** | **p-value** |
| **CUSCH-LOAD** | **LOPO-EA** | 0.4014 | 0.4306 | 0.3272 | -0.00393 | 1.394E-75 |
| | **LOPO-EL** | 0.4319 | 0.4467 | 0.3206 | 0.00066 | 0.8590188 |
| | **LOPO-AL** | 0.4212 | 0.4419 | 0.3270 | 0.00009 | > 0.99 |
| | **LOPO-all** | 0.4164 | 0.4383 | 0.3264 | - | - |
| **DR** | **LOPO-EA** | 0.3943 | 0.4274 | 0.3264 | -0.00332 | 8.847E-57 |
| | **LOPO-EL** | 0.4210 | 0.4404 | 0.3197 | -0.00053 | > 0.99 |
| | **LOPO-AL** | 0.4132 | 0.4379 | 0.3262 | 0.00026 | > 0.99 |
| | **LOPO-all** | 0.4087 | 0.4343 | 0.3256 | - | - |
| **PR** | **LOPO-EA** | 0.4663 | 0.4677 | 0.3375 | -0.00793 | 9.55E-131 |
| | **LOPO-EL** | 0.5232 | 0.5025 | 0.3307 | 0.00807 | 5.798E-38 |
| | **LOPO-AL** | 0.4893 | 0.4824 | 0.3366 | -0.00206 | 4.197E-08 |
| | **LOPO-all** | 0.4928 | 0.4821 | 0.3365 | - | - |
| **LARGE-PD** | **LOPO-EA** | 0.4074 | 0.4388 | 0.3228 | -0.02626 | < 1.34E-307 |
| | **LOPO-EL** | 0.4990 | 0.4985 | 0.3155 | 0.01293 | 2.04E-195 |
| | **LOPO-AL** | 0.4624 | 0.4745 | 0.3194 | -0.00348 | 1.254E-36 |
| | **LOPO-all** | 0.4624 | 0.4746 | 0.3194 | - | - |
| **SIGMA** | **LOPO-EA** | 0.7286 | 0.6317 | 0.3336 | -0.02180 | < 1.34E-307 |
| | **LOPO-EL** | 0.7902 | 0.6855 | 0.3085 | 0.00580 | 3.94E-173 |
| | **LOPO-AL** | 0.7758 | 0.6727 | 0.3148 | 0.00014 | 0.9467682 |
| | **LOPO-all** | 0.7737 | 0.6689 | 0.3168 | - | - |
| **Bambui** | **LOPO-EA** | 0.7321 | 0.6527 | 0.3037 | -0.00460 | 5.31E-232 |
| | **LOPO-EL** | 0.7344 | 0.6581 | 0.2977 | -0.00285 | 1.621E-32 |
| | **LOPO-AL** | 0.7529 | 0.6666 | 0.3017 | 0.00002 | > 0.99 |
| | **LOPO-all** | 0.7459 | 0.6625 | 0.3009 | - | - |

| | | | | | | |
|---|---|---|---|---|---|---|
| **Pelotas** | **LOPO-EA** | 0.7118 | 0.6414 | 0.3012 | -0.00484 | < 1.34E-307 |
| | **LOPO-EL** | 0.7099 | 0.6439 | 0.2953 | -0.00520 | 1.27E-121 |
| | **LOPO-AL** | 0.7326 | 0.6543 | 0.2995 | -0.00034 | 0.0775884 |
| | **LOPO-all** | 0.7253 | 0.6511 | 0.2979 | - | - |
| **Salvador** | **LOPO-EA** | 0.6813 | 0.6192 | 0.3010 | -0.00342 | 4.5E-141 |
| | **LOPO-EL** | 0.6475 | 0.5987 | 0.2975 | -0.02268 | < 1.34E-307 |
| | **LOPO-AL** | 0.7072 | 0.6375 | 0.2982 | 0.00630 | < 1.34E-307 |
| | **LOPO-all** | 0.6912 | 0.6262 | 0.2991 | - | - |

ian of differences calculated using a paired, two-sided Wilcoxon signed rank test
ch the same direction of effect was seen. Complete pairwise comparison resutls

| # of replicates matching | Panel | Chromosome<br>Median | Mean | Std dev |
|---|---|---|---|---|
| 5/5 | **LOPO-EA** | 0.5264 | 0.5107 | 0.2973 |
| 5/5 | **LOPO-EL** | 0.5341 | 0.5188 | 0.2915 |
| 5/5 | **LOPO-AL** | 0.5453 | 0.5290 | 0.2893 |
| | **LOPO-all** | 0.5416 | 0.5252 | 0.2913 |
| 5/5 | **LOPO-EA** | 0.5105 | 0.5015 | 0.2974 |
| 3/5 | **LOPO-EL** | 0.5138 | 0.5067 | 0.2927 |
| 5/5 | **LOPO-AL** | 0.5283 | 0.5181 | 0.2906 |
| | **LOPO-all** | 0.5237 | 0.5140 | 0.2925 |
| 5/5 | **LOPO-EA** | 0.6330 | 0.5777 | 0.3097 |
| 5/5 | **LOPO-EL** | 0.6795 | 0.6178 | 0.2933 |
| 5/5 | **LOPO-AL** | 0.6724 | 0.6148 | 0.2923 |
| | **LOPO-all** | 0.6708 | 0.6125 | 0.2948 |
| 5/5 | **LOPO-EA** | 0.6276 | 0.5872 | 0.2943 |
| 5/5 | **LOPO-EL** | 0.6572 | 0.6136 | 0.2825 |
| 5/5 | **LOPO-AL** | 0.6580 | 0.6163 | 0.2796 |
| | **LOPO-all** | 0.6610 | 0.6168 | 0.2819 |
| 5/5 | **LOPO-EA** | 0.9012 | 0.7734 | 0.2749 |
| 5/5 | **LOPO-EL** | 0.9212 | 0.8003 | 0.2548 |
| 1/5 | **LOPO-AL** | 0.9177 | 0.7993 | 0.2534 |
| | **LOPO-all** | 0.9172 | 0.7972 | 0.2561 |
| 5/5 | **LOPO-EA** | 0.8488 | 0.7543 | 0.2545 |
| 5/5 | **LOPO-EL** | 0.8463 | 0.7539 | 0.2515 |
| 3/5 | **LOPO-AL** | 0.8542 | 0.7618 | 0.2485 |
| | **LOPO-all** | 0.8545 | 0.7611 | 0.2494 |

| | | | | |
|---|---|---|---|---|
| 5/5 | **LOPO-EA** | 0.8320 | 0.7518 | 0.2414 |
| 5/5 | **LOPO-EL** | 0.8258 | 0.7495 | 0.2382 |
| 5/5 | **LOPO-AL** | 0.8367 | 0.7591 | 0.2349 |
| | **LOPO-all** | 0.8375 | 0.7585 | 0.2362 |
| 5/5 | **LOPO-EA** | 0.7729 | 0.7179 | 0.2379 |
| 5/5 | **LOPO-EL** | 0.7523 | 0.7006 | 0.2408 |
| 5/5 | **LOPO-AL** | 0.7818 | 0.7281 | 0.2309 |
| | **LOPO-all** | 0.7757 | 0.7219 | 0.2344 |

t, using panel LOPO-all as a
can be found in Table S9.

**X**

| Median of differences | p-value | # of replicates matching |
|---|---|---|
| -0.01196 | < 1.34E-307 | 5/5 |
| -0.00596 | 7.65E-191 | 5/5 |
| 0.00353 | 2.35E-122 | 5/5 |
| - | - | |
| -0.01034 | < 1.34E-307 | 5/5 |
| -0.00676 | 1.83E-220 | 5/5 |
| 0.00383 | 4.81E-138 | 5/5 |
| - | - | |
| -0.02654 | < 1.34E-307 | 5/5 |
| 0.00287 | 2.247E-29 | 5/5 |
| 0.00129 | 4.455E-10 | 4/5 |
| - | - | |
| -0.0238003 | < 1.34E-307 | 5/5 |
| 0.0004182 | 0.5533102 | 2/5 |
| -0.0010898 | 5.205E-06 | 5/5 |
| - | - | |
| -0.0113516 | < 1.34E-307 | 5/5 |
| 0.0014122 | 3.61E-165 | 5/5 |
| 0.0003822 | 3.658E-22 | 5/5 |
| - | - | |
| -0.0036412 | < 1.34E-307 | 5/5 |
| -0.0032944 | 9.94E-242 | 5/5 |
| 8.591E-06 | > 0.99 | 5/5 |
| - | - | |

| | | |
|---|---|---|
| -0.0037427 | < 1.34E-307 | 5/5 |
| -0.0045718 | < 1.34E-307 | 5/5 |
| -2.647E-05 | > 0.99 | 3/5 |
| - | - | |
| -0.0019028 | 6.27E-144 | 5/5 |
| -0.016376 | < 1.34E-307 | 5/5 |
| 0.0042327 | < 1.34E-307 | 5/5 |
| - | - | |

**Table S9. Full pairwise comparison results for LOPO reference panels.** Medians,
paired, two-sided Wilcoxon signed rank test, replicated five times (A-E). P-values are
median of differences are in the upper triangle of the table (above the "-"). The media

| Population | | | | | | |
|---|---|---|---|---|---|---|
| | | | CUSCH-LOAD_chr7 | | | |
| | **Panel** | **Median** | **P-values** | LOPO-EA | LOPO-EL | LOPO-AL |
| | LOPO-EA | 0.4014 | LOPO-EA | - | 0.0060629 | 0.0048791 |
| | LOPO-EL | 0.43189 | LOPO-EL | 1.01E-22 | - | -0.0001292 |
| | LOPO-AL | 0.421165 | LOPO-AL | 5.45E-33 | > 0.99 | - |
| | LOPO-all | 0.416395 | LOPO-all | 1.39E-75 | 8.59E-01 | > 0.99 |
| | | | DR_chr7 | | | |
| | **Panel** | **Median** | **P-values** | LOPO-EA | LOPO-EL | LOPO-AL |
| | LOPO-EA | 0.394345 | LOPO-EA | - | 0.0038186 | 0.0044708 |
| | LOPO-EL | 0.421035 | LOPO-EL | 4.96E-10 | - | 0.0011393 |
| | LOPO-AL | 0.413235 | LOPO-AL | 1.17E-28 | 8.43E-02 | - |
| | LOPO-all | 0.40869 | LOPO-all | 8.85E-57 | > 0.99 | > 0.99 |
| | | | PR_chr7 | | | |
| | **Panel** | **Median** | **P-values** | LOPO-EA | LOPO-EL | LOPO-AL |
| | LOPO-EA | 0.46631 | LOPO-EA | - | 0.0191850 | 0.0065980 |
| | LOPO-EL | 0.52318 | LOPO-EL | 1.18E-128 | - | -0.0099737 |
| | LOPO-AL | 0.48929 | LOPO-AL | 2.41E-37 | 4.27E-55 | - |
| | LOPO-all | 0.49278 | LOPO-all | 9.55E-131 | 5.80E-38 | 4.20E-08 |
| | | | LARGE-PD_chr7 | | | |
| | **Panel** | **Median** | **P-values** | LOPO-EA | LOPO-EL | LOPO-AL |
| | LOPO-EA | 0.40744 | LOPO-EA | - | 0.0454559 | 0.0240921 |
| | LOPO-EL | 0.49904 | LOPO-EL | < 1.34E-307 | - | -0.0162744 |
| | LOPO-AL | 0.4624 | LOPO-AL | < 1.34E-307 | 1.84E-303 | - |
| | LOPO-all | 0.46243 | LOPO-all | < 1.34E-307 | 2.04E-195 | 1.25E-36 |
| | | | SIGMA_chr7 | | | |
| | **Panel** | **Median** | **P-values** | LOPO-EA | LOPO-EL | LOPO-AL |
| | LOPO-EA | 0.72864 | LOPO-EA | - | 0.0330466 | 0.0231190 |
| | LOPO-EL | 0.79016 | LOPO-EL | < 1.34E-307 | - | -0.0051150 |
| | LOPO-AL | 0.77582 | LOPO-AL | < 1.34E-307 | 4.03E-141 | - |
| | LOPO-all | 0.77371 | LOPO-all | < 1.34E-307 | 3.94E-173 | 9.47E-01 |
| | | | Bambui_chr7 | | | |
| | **Panel** | **Median** | **P-values** | LOPO-EA | LOPO-EL | LOPO-AL |
| | LOPO-EA | 0.73211 | LOPO-EA | - | 0.0022758 | 0.0057286 |

| | | | | | |
|---|---|---|---|---|---|
| LOPO-EL | 0.73439 | LOPO-EL | 1.40E-15 | - | 0.0035271 |
| LOPO-AL | 0.75293 | LOPO-AL | 8.98E-156 | 3.73E-51 | - |
| LOPO-all | 0.74593 | LOPO-all | 5.31E-232 | 1.62E-32 | > 0.99 |

| | | | Pelotas_chr7 | | |
|---|---|---|---|---|---|
| **Panel** | **Median** | **P-values** | LOPO-EA | LOPO-EL | LOPO-AL |
| LOPO-EA | 0.71182 | LOPO-EA | - | 0.0003302 | 0.0054297 |
| LOPO-EL | 0.7099 | LOPO-EL | 9.56E-01 | - | 0.0055652 |
| LOPO-AL | 0.73255 | LOPO-AL | 1.82E-175 | 5.75E-138 | - |
| LOPO-all | 0.72525 | LOPO-all | < 1.34E-307 | 1.27E-121 | 7.76E-02 |

| | | | Salvador_chr7 | | |
|---|---|---|---|---|---|
| **Panel** | **Median** | **P-values** | LOPO-EA | LOPO-EL | LOPO-AL |
| LOPO-EA | 0.68126 | LOPO-EA | - | -0.0175962 | 0.0112689 |
| LOPO-EL | 0.64745 | LOPO-EL | < 1.34E-307 | - | 0.0308890 |
| LOPO-AL | 0.70716 | LOPO-AL | < 1.34E-307 | < 1.34E-307 | - |
| LOPO-all | 0.69123 | LOPO-all | 4.50E-141 | < 1.34E-307 | < 1.34E-307 |

p-values and median of differences calculated using a
found in the lower triangle of the comparison, while the
n difference is column - row.

Comparison A

**CUSCH-LOAD_chrX**

| LOPO-all | Panel | Median | P-values | LOPO-EA | LOPO-EL |
|---|---|---|---|---|---|
| 0.0039277 | LOPO-EA | 0.526395 | LOPO-EA | - | 0.0066130 |
| -0.0006632 | LOPO-EL | 0.534105 | LOPO-EL | 6.23E-106 | - |
| -0.0000869 | LOPO-AL | 0.54527 | LOPO-AL | < 1.34E-307 | 1.39E-253 |
| - | LOPO-all | 0.541625 | LOPO-all | < 1.34E-307 | 7.65E-191 |

**DR_chrX**

| LOPO-all | Panel | Median | P-values | LOPO-EA | LOPO-EL |
|---|---|---|---|---|---|
| 0.0033161 | LOPO-EA | 0.51047 | LOPO-EA | - | 0.0043612 |
| 0.0005332 | LOPO-EL | 0.51381 | LOPO-EL | 1.01E-47 | - |
| -0.0002574 | LOPO-AL | 0.52826 | LOPO-AL | < 1.34E-307 | 3.54E-293 |
| - | LOPO-all | 0.52368 | LOPO-all | < 1.34E-307 | 1.83E-220 |

**PR_chrX**

| LOPO-all | Panel | Median | P-values | LOPO-EA | LOPO-EL |
|---|---|---|---|---|---|
| 0.0079269 | LOPO-EA | 0.63295 | LOPO-EA | - | 0.0314439 |
| -0.0080678 | LOPO-EL | 0.67952 | LOPO-EL | < 1.34E-307 | - |
| 0.0020566 | LOPO-AL | 0.67236 | LOPO-AL | < 1.34E-307 | 4.13E-02 |
| - | LOPO-all | 0.67075 | LOPO-all | < 1.34E-307 | 2.25E-29 |

**LARGE-PD_chrX**

| LOPO-all | Panel | Median | P-values | LOPO-EA | LOPO-EL |
|---|---|---|---|---|---|
| 0.0262579 | LOPO-EA | 0.627575 | LOPO-EA | - | 0.0244466 |
| -0.0129255 | LOPO-EL | 0.657245 | LOPO-EL | < 1.34E-307 | - |
| 0.0034814 | LOPO-AL | 0.657995 | LOPO-AL | < 1.34E-307 | 2.57E-01 |
| - | LOPO-all | 0.661025 | LOPO-all | < 1.34E-307 | 5.53E-01 |

**SIGMA_chrX**

| LOPO-all | Panel | Median | P-values | LOPO-EA | LOPO-EL |
|---|---|---|---|---|---|
| 0.0218017 | LOPO-EA | 0.90115 | LOPO-EA | - | 0.0140128 |
| -0.0058028 | LOPO-EL | 0.92118 | LOPO-EL | < 1.34E-307 | - |
| -0.0001377 | LOPO-AL | 0.917715 | LOPO-AL | < 1.34E-307 | 8.67E-18 |
| - | LOPO-all | 0.917155 | LOPO-all | < 1.34E-307 | 3.61E-165 |

**Bambui_chrX**

| LOPO-all | Panel | Median | P-values | LOPO-EA | LOPO-EL |
|---|---|---|---|---|---|
| 0.0046028 | LOPO-EA | 0.848805 | LOPO-EA | - | 0.0004611 |

| | | | | | |
|---|---|---|---|---|---|
| 0.0028516 | LOPO-EL | 0.84631 | LOPO-EL | 4.71E-04 | - |
| -0.0000240 | LOPO-AL | 0.854235 | LOPO-AL | 5.59E-296 | 5.65E-190 |
| - | LOPO-all | 0.854455 | LOPO-all | < 1.34E-307 | 9.94E-242 |

**Pelotas_chrX**

| LOPO-all | **Panel** | **Median** | **P-values** | LOPO-EA | LOPO-EL |
|---|---|---|---|---|---|
| 0.0048370 | LOPO-EA | 0.831965 | LOPO-EA | - | -0.0006792 |
| 0.0051991 | LOPO-EL | 0.825845 | LOPO-EL | 7.39E-09 | - |
| 0.0003391 | LOPO-AL | 0.836665 | LOPO-AL | < 1.34E-307 | < 1.34E-307 |
| - | LOPO-all | 0.837515 | LOPO-all | < 1.34E-307 | < 1.34E-307 |

**Salvador_chrX**

| LOPO-all | **Panel** | **Median** | **P-values** | LOPO-EA | LOPO-EL |
|---|---|---|---|---|---|
| 0.0034181 | LOPO-EA | 0.77285 | LOPO-EA | - | -0.0132777 |
| 0.0226768 | LOPO-EL | 0.75227 | LOPO-EL | < 1.34E-307 | - |
| -0.0062961 | LOPO-AL | 0.78182 | LOPO-AL | < 1.34E-307 | < 1.34E-307 |
| - | LOPO-all | 0.77565 | LOPO-all | 6.27E-144 | < 1.34E-307 |

-

C

| LOPO-AL | LOPO-all | **Panel** | **Median** | **P-values** |
|---|---|---|---|---|
| 0.0154978 | 0.0119638 | LOPO-EA | 0.400895 | LOPO-EA |
| 0.0098617 | 0.0059622 | LOPO-EL | 0.43263 | LOPO-EL |
| - | -0.0035259 | LOPO-AL | 0.420395 | LOPO-AL |
| 2.35E-122 | - | LOPO-all | 0.417755 | LOPO-all |

| LOPO-AL | LOPO-all | **Panel** | **Median** | **P-values** |
|---|---|---|---|---|
| 0.0142668 | 0.0103424 | LOPO-EA | 0.39572 | LOPO-EA |
| 0.0109904 | 0.0067581 | LOPO-EL | 0.42285 | LOPO-EL |
| - | -0.0038339 | LOPO-AL | 0.41403 | LOPO-AL |
| 4.81E-138 | - | LOPO-all | 0.40999 | LOPO-all |

| LOPO-AL | LOPO-all | **Panel** | **Median** | **P-values** |
|---|---|---|---|---|
| 0.0280671 | 0.0265398 | LOPO-EA | 0.46485 | LOPO-EA |
| -0.0008620 | -0.0028729 | LOPO-EL | 0.52557 | LOPO-EL |
| - | -0.0012948 | LOPO-AL | 0.4873 | LOPO-AL |
| 4.46E-10 | - | LOPO-all | 0.48988 | LOPO-all |

| LOPO-AL | LOPO-all | **Panel** | **Median** | **P-values** |
|---|---|---|---|---|
| 0.0224788 | 0.0238003 | LOPO-EA | 0.41037 | LOPO-EA |
| -0.0006516 | -0.0004182 | LOPO-EL | 0.498535 | LOPO-EL |
| - | 0.0010898 | LOPO-AL | 0.46764 | LOPO-AL |
| 5.20E-06 | - | LOPO-all | 0.46434 | LOPO-all |

| LOPO-AL | LOPO-all | **Panel** | **Median** | **P-values** |
|---|---|---|---|---|
| 0.0115939 | 0.0113516 | LOPO-EA | 0.729975 | LOPO-EA |
| -0.0004332 | -0.0014122 | LOPO-EL | 0.79083 | LOPO-EL |
| - | -0.0003822 | LOPO-AL | 0.778365 | LOPO-AL |
| 3.66E-22 | - | LOPO-all | 0.77306 | LOPO-all |

| LOPO-AL | LOPO-all | **Panel** | **Median** | **P-values** |
|---|---|---|---|---|
| 0.0034960 | 0.0036412 | LOPO-EA | 0.73259 | LOPO-EA |

| | |
|---|---|
| 0.0037220 | 0.0032944 |
| - | -0.0000072 |
| > 0.99 | - |

| | | |
|---|---|---|
| LOPO-EL | 0.73398 | LOPO-EL |
| LOPO-AL | 0.753935 | LOPO-AL |
| LOPO-all | 0.74455 | LOPO-all |

| LOPO-AL | LOPO-all |
|---|---|
| 0.0036789 | 0.0037427 |
| 0.0049808 | 0.0045718 |
| - | 0.0000265 |
| > 0.99 | - |

| **Panel** | **Median** | **P-values** |
|---|---|---|
| LOPO-EA | 0.71288 | LOPO-EA |
| LOPO-EL | 0.70981 | LOPO-EL |
| LOPO-AL | 0.73367 | LOPO-AL |
| LOPO-all | 0.72468 | LOPO-all |

| LOPO-AL | LOPO-all |
|---|---|
| 0.0062003 | 0.0019028 |
| 0.0218033 | 0.0163760 |
| - | -0.0042327 |
| < 1.34E-307 | - |

| **Panel** | **Median** | **P-values** |
|---|---|---|
| LOPO-EA | 0.681905 | LOPO-EA |
| LOPO-EL | 0.64693 | LOPO-EL |
| LOPO-AL | 0.70788 | LOPO-AL |
| LOPO-all | 0.6905 | LOPO-all |

Comparison B

**USCH-LOAD_chr7**

| LOPO-EA | LOPO-EL | LOPO-AL | LOPO-all | Panel | Median |
|---|---|---|---|---|---|
| - | 0.0065059 | 0.0046061 | 0.0044612 | LOPO-EA | 0.526185 |
| 5.62E-27 | - | -0.0006907 | -0.0001467 | LOPO-EL | 0.53156 |
| 2.52E-32 | 7.73E-01 | - | 0.0004328 | LOPO-AL | 0.545035 |
| 9.77E-105 | > 0.99 | 5.88E-01 | - | LOPO-all | 0.54135 |

**DR_chr7**

| LOPO-EA | LOPO-EL | LOPO-AL | LOPO-all | Panel | Median |
|---|---|---|---|---|---|
| - | 0.0043662 | 0.0042705 | 0.0036919 | LOPO-EA | 0.51112 |
| 4.85E-13 | - | 0.0007163 | 0.0009579 | LOPO-EL | 0.51301 |
| 1.43E-27 | 8.05E-01 | - | 0.0000516 | LOPO-AL | 0.52755 |
| 2.88E-74 | 2.39E-01 | > 0.99 | - | LOPO-all | 0.52354 |

**PR_chr7**

| LOPO-EA | LOPO-EL | LOPO-AL | LOPO-all | Panel | Median |
|---|---|---|---|---|---|
| - | 0.0201282 | 0.0060760 | 0.0082711 | LOPO-EA | 0.63378 |
| 1.25E-139 | - | -0.0112224 | -0.0084475 | LOPO-EL | 0.67835 |
| 3.65E-33 | 2.27E-67 | - | 0.0025417 | LOPO-AL | 0.67105 |
| 4.52E-154 | 1.11E-39 | 8.71E-13 | - | LOPO-all | 0.66933 |

**LARGE-PD_chr7**

| LOPO-EA | LOPO-EL | LOPO-AL | LOPO-all | Panel | Median |
|---|---|---|---|---|---|
| - | 0.0453600 | 0.0243243 | 0.0253863 | LOPO-EA | 0.62658 |
| < 1.34E-307 | - | -0.0155603 | -0.0134957 | LOPO-EL | 0.65667 |
| < 1.34E-307 | 1.90E-276 | - | 0.0027947 | LOPO-AL | 0.66012 |
| < 1.34E-307 | 3.24E-209 | 6.57E-25 | - | LOPO-all | 0.661465 |

**SIGMA_chr7**

| LOPO-EA | LOPO-EL | LOPO-AL | LOPO-all | Panel | Median |
|---|---|---|---|---|---|
| - | 0.0326441 | 0.0227249 | 0.0208959 | LOPO-EA | 0.9014 |
| < 1.34E-307 | - | -0.0050861 | -0.0061192 | LOPO-EL | 0.92108 |
| < 1.34E-307 | 3.77E-142 | - | -0.0003019 | LOPO-AL | 0.91824 |
| < 1.34E-307 | 1.94E-188 | 1.57E-02 | - | LOPO-all | 0.91769 |

**Bambui_chr7**

| LOPO-EA | LOPO-EL | LOPO-AL | LOPO-all | Panel | Median |
|---|---|---|---|---|---|
| - | 0.0018490 | 0.0062748 | 0.0046868 | LOPO-EA | 0.84894 |

| | | | |
|---|---|---|---|
| 1.08E-10 | - | 0.0040769 | 0.0031094 |
| 1.03E-179 | 1.17E-64 | - | -0.0003241 |
| 8.76E-252 | 9.87E-39 | 1.05E-01 | - |

| | |
|---|---|
| LOPO-EL | 0.84629 |
| LOPO-AL | 0.85438 |
| LOPO-all | 0.854815 |

**Pelotas_chr7**

| LOPO-EA | LOPO-EL | LOPO-AL | LOPO-all |
|---|---|---|---|
| - | 0.0000979 | 0.0054333 | 0.0043996 |
| > 0.99 | - | 0.0057773 | 0.0051463 |
| 3.51E-180 | 4.32E-146 | - | -0.0001288 |
| < 1.34E-307 | 4.87E-118 | > 0.99 | - |

| **Panel** | **Median** |
|---|---|
| LOPO-EA | 0.83226 |
| LOPO-EL | 0.82525 |
| LOPO-AL | 0.83755 |
| LOPO-all | 0.83767 |

**Salvador_chr7**

| LOPO-EA | LOPO-EL | LOPO-AL | LOPO-all |
|---|---|---|---|
| - | -0.0184320 | 0.0112598 | 0.0028631 |
| < 1.34E-307 | - | 0.0316622 | 0.0228343 |
| < 1.34E-307 | < 1.34E-307 | - | -0.0069269 |
| 1.05E-106 | < 1.34E-307 | < 1.34E-307 | - |

| **Panel** | **Median** |
|---|---|
| LOPO-EA | 0.77333 |
| LOPO-EL | 0.750945 |
| LOPO-AL | 0.78208 |
| LOPO-all | 0.77653 |

**CUSCH-LOAD_chrX**

| **P-values** | LOPO-EA | LOPO-EL | LOPO-AL | LOPO-all |
|---|---|---|---|---|
| LOPO-EA | - | 0.0054912 | 0.0149358 | 0.0109596 |
| LOPO-EL | 1.38E-73 | - | 0.0105449 | 0.0062770 |
| LOPO-AL | 0.00E+00 | 6.44E-276 | - | -0.0038089 |
| LOPO-all | 0.00E+00 | 2.06E-199 | 2.99E-144 | - |

**DR_chrX**

| **P-values** | LOPO-EA | LOPO-EL | LOPO-AL | LOPO-all |
|---|---|---|---|---|
| LOPO-EA | - | 0.0031079 | 0.0135645 | 0.0091110 |
| LOPO-EL | 1.60E-24 | - | 0.0116766 | 0.0069608 |
| LOPO-AL | 0.00E+00 | | - | -0.0042632 |
| LOPO-all | 0.00E+00 | 6.94E-220 | 7.60E-174 | - |

**PR_chrX**

| **P-values** | LOPO-EA | LOPO-EL | LOPO-AL | LOPO-all |
|---|---|---|---|---|
| LOPO-EA | - | 0.0305602 | 0.0269264 | 0.0262587 |
| LOPO-EL | < 1.34E-307 | - | -0.0005342 | -0.0021499 |
| LOPO-AL | < 1.34E-307 | 5.84E-01 | - | -0.0010874 |
| LOPO-all | < 1.34E-307 | 3.12E-17 | 2.34E-07 | - |

**LARGE-PD_chrX**

| **P-values** | LOPO-EA | LOPO-EL | LOPO-AL | LOPO-all |
|---|---|---|---|---|
| LOPO-EA | - | 0.0249569 | 0.0235567 | 0.0245880 |
| LOPO-EL | < 1.34E-307 | - | -0.0004494 | -0.0000427 |
| LOPO-AL | < 1.34E-307 | 9.28E-01 | - | 0.0011385 |
| LOPO-all | < 1.34E-307 | > 0.99 | 1.99E-06 | - |

**SIGMA_chrX**

| **P-values** | LOPO-EA | LOPO-EL | LOPO-AL | LOPO-all |
|---|---|---|---|---|
| LOPO-EA | - | 0.0143063 | 0.0116576 | 0.0117250 |
| LOPO-EL | < 1.34E-307 | - | -0.0004286 | -0.0012433 |
| LOPO-AL | < 1.34E-307 | 4.57E-16 | - | -0.0003214 |
| LOPO-all | < 1.34E-307 | 3.72E-132 | 1.26E-16 | - |

**Bambui_chrX**

| **P-values** | LOPO-EA | LOPO-EL | LOPO-AL | LOPO-all |
|---|---|---|---|---|
| LOPO-EA | - | 0.0003372 | 0.0035756 | 0.0036743 |

| | | | | |
|---|---|---|---|---|
| LOPO-EL | 3.59E-02 | - | 0.0039784 | 0.0035549 |
| LOPO-AL | < 1.34E-307 | 1.39E-213 | - | -0.0000525 |
| LOPO-all | < 1.34E-307 | 3.48E-273 | > 0.99 | - |

**Pelotas_chrX**

| **P-values** | LOPO-EA | LOPO-EL | LOPO-AL | LOPO-all |
|---|---|---|---|---|
| LOPO-EA | - | -0.0010830 | 0.0037972 | 0.0038919 |
| LOPO-EL | 1.17E-21 | - | 0.0056426 | 0.0052280 |
| LOPO-AL | < 1.34E-307 | < 1.34E-307 | - | 0.0000222 |
| LOPO-all | < 1.34E-307 | < 1.34E-307 | < 1.34E-307 | - |

**Salvador_chrX**

| **P-values** | LOPO-EA | LOPO-EL | LOPO-AL | LOPO-all |
|---|---|---|---|---|
| LOPO-EA | - | -0.0141862 | 0.0062594 | 0.0020924 |
| LOPO-EL | < 1.34E-307 | - | 0.0227872 | 0.0174908 |
| LOPO-AL | < 1.34E-307 | < 1.34E-307 | - | -0.0041015 |
| LOPO-all | 1.08E-170 | < 1.34E-307 | < 1.34E-307 | - |

**CUSCH-LOAD_chr7**

| **Panel** | **Median** | **P-values** | LOPO-EA | LOPO-EL | LOPO-AL | LOPO-all |
|---|---|---|---|---|---|---|
| LOPO-EA | 0.40123 | LOPO-EA | - | 0.0065196 | 0.0055748 | 0.0047443 |
| LOPO-EL | 0.43116 | LOPO-EL | 1.47E-26 | - | 0.0002046 | 0.0001792 |
| LOPO-AL | 0.41944 | LOPO-AL | 1.04E-45 | > 0.99 | - | 0.0002602 |
| LOPO-all | 0.41784 | LOPO-all | 2.67E-110 | > 0.99 | > 0.99 | - |

**DR_chr7**

| **Panel** | **Median** | **P-values** | LOPO-EA | LOPO-EL | LOPO-AL | LOPO-all |
|---|---|---|---|---|---|---|
| LOPO-EA | 0.397075 | LOPO-EA | - | 0.0039836 | 0.0049382 | 0.0040957 |
| LOPO-EL | 0.420295 | LOPO-EL | 6.26E-11 | - | 0.0018475 | 0.0016768 |
| LOPO-AL | 0.41258 | LOPO-AL | 2.76E-36 | 5.30E-04 | - | 0.0001441 |
| LOPO-all | 0.410545 | LOPO-all | 1.55E-84 | 1.74E-03 | > 0.99 | - |

**PR_chr7**

| **Panel** | **Median** | **P-values** | LOPO-EA | LOPO-EL | LOPO-AL | LOPO-all |
|---|---|---|---|---|---|---|
| LOPO-EA | 0.466255 | LOPO-EA | - | 0.0191959 | 0.0068327 | 0.0081866 |
| LOPO-EL | 0.520595 | LOPO-EL | 2.00E-133 | - | -0.0098347 | -0.0072630 |
| LOPO-AL | 0.488955 | LOPO-AL | 6.68E-41 | 2.57E-55 | - | 0.0022936 |
| LOPO-all | 0.489635 | LOPO-all | 3.68E-145 | 3.18E-33 | 1.61E-10 | - |

**LARGE-PD_chr7**

| **Panel** | **Median** | **P-values** | LOPO-EA | LOPO-EL | LOPO-AL | LOPO-all |
|---|---|---|---|---|---|---|
| LOPO-EA | 0.41078 | LOPO-EA | - | 0.0443201 | 0.0236764 | 0.0253583 |
| LOPO-EL | 0.499355 | LOPO-EL | < 1.34E-307 | - | -0.0154141 | -0.0123878 |
| LOPO-AL | 0.465225 | LOPO-AL | < 1.34E-307 | 6.84E-280 | - | 0.0034024 |
| LOPO-all | 0.46466 | LOPO-all | < 1.34E-307 | 8.54E-187 | 5.45E-35 | - |

**SIGMA_chr7**

| **Panel** | **Median** | **P-values** | LOPO-EA | LOPO-EL | LOPO-AL | LOPO-all |
|---|---|---|---|---|---|---|
| LOPO-EA | 0.73149 | LOPO-EA | - | 0.0327486 | 0.0232490 | 0.0213520 |
| LOPO-EL | 0.79169 | LOPO-EL | < 1.34E-307 | - | -0.0047356 | -0.0059934 |
| LOPO-AL | 0.77921 | LOPO-AL | < 1.34E-307 | 5.20E-128 | - | -0.0003151 |
| LOPO-all | 0.77356 | LOPO-all | < 1.34E-307 | 9.75E-192 | 1.17E-02 | - |

**Bambui_chr7**

| **Panel** | **Median** | **P-values** | LOPO-EA | LOPO-EL | LOPO-AL | LOPO-all |
|---|---|---|---|---|---|---|
| LOPO-EA | 0.73314 | LOPO-EA | - | 0.0019810 | 0.0054755 | 0.0043900 |

| | | | | | | |
|---|---|---|---|---|---|---|
| LOPO-EL | 0.73571 | LOPO-EL | 2.32E-12 | - | 0.0034648 | 0.0027687 |
| LOPO-AL | 0.75332 | LOPO-AL | 6.58E-144 | 7.07E-49 | - | -0.0001348 |
| LOPO-all | 0.74709 | LOPO-all | 1.54E-220 | 3.46E-32 | > 0.99 | - |

**Pelotas_chr7**

| **Panel** | **Median** | **P-values** | LOPO-EA | LOPO-EL | LOPO-AL | LOPO-all |
|---|---|---|---|---|---|---|
| LOPO-EA | 0.71309 | LOPO-EA | - | -0.0000490 | 0.0053226 | 0.0043686 |
| LOPO-EL | 0.71038 | LOPO-EL | > 0.99 | - | 0.0059426 | 0.0051575 |
| LOPO-AL | 0.73348 | LOPO-AL | 7.45E-173 | 4.20E-157 | - | 0.0000686 |
| LOPO-all | 0.72566 | LOPO-all | < 1.34E-307 | 3.32E-125 | > 0.99 | - |

**Salvador_chr7**

| **Panel** | **Median** | **P-values** | LOPO-EA | LOPO-EL | LOPO-AL | LOPO-all |
|---|---|---|---|---|---|---|
| LOPO-EA | 0.68261 | LOPO-EA | - | -0.0173408 | 0.0116098 | 0.0030953 |
| LOPO-EL | 0.64936 | LOPO-EL | < 1.34E-307 | - | 0.0313946 | 0.0221018 |
| LOPO-AL | 0.70888 | LOPO-AL | < 1.34E-307 | < 1.34E-307 | - | -0.0070782 |
| LOPO-all | 0.69141 | LOPO-all | 1.41E-122 | < 1.34E-307 | < 1.34E-307 | - |

Comparison C

**CUSCH-LOAD_chrX**

| **Panel** | **Median** | **P-values** | LOPO-EA | LOPO-EL | LOPO-AL |
|---|---|---|---|---|---|
| LOPO-EA | 0.52639 | LOPO-EA | - | 0.0059476 | 0.0141365 |
| LOPO-EL | 0.53198 | LOPO-EL | 6.00E-86 | - | 0.0093207 |
| LOPO-AL | 0.54084 | LOPO-AL | < 1.34E-307 | 3.81E-223 | - |
| LOPO-all | 0.54093 | LOPO-all | < 1.34E-307 | 2.33E-243 | 5.03E-42 |

**DR_chrX**

| **Panel** | **Median** | **P-values** | LOPO-EA | LOPO-EL | LOPO-AL |
|---|---|---|---|---|---|
| LOPO-EA | 0.509445 | LOPO-EA | - | 0.0034369 | 0.0129061 |
| LOPO-EL | 0.511255 | LOPO-EL | 3.17E-30 | - | 0.0106585 |
| LOPO-AL | 0.52446 | LOPO-AL | < 1.34E-307 | 2.70E-272 | - |
| LOPO-all | 0.524515 | LOPO-all | < 1.34E-307 | 1.36E-272 | 1.34E-58 |

**PR_chrX**

| **Panel** | **Median** | **P-values** | LOPO-EA | LOPO-EL | LOPO-AL |
|---|---|---|---|---|---|
| LOPO-EA | 0.631485 | LOPO-EA | - | 0.0308435 | 0.0259983 |
| LOPO-EL | 0.67768 | LOPO-EL | < 1.34E-307 | - | -0.0017458 |
| LOPO-AL | 0.66883 | LOPO-AL | < 1.34E-307 | 3.27E-07 | - |
| LOPO-all | 0.671535 | LOPO-all | < 1.34E-307 | 9.25E-07 | 1.48E-05 |

**LARGE-PD_chrX**

| **Panel** | **Median** | **P-values** | LOPO-EA | LOPO-EL | LOPO-AL |
|---|---|---|---|---|---|
| LOPO-EA | 0.62758 | LOPO-EA | - | 0.0242842 | 0.0201909 |
| LOPO-EL | 0.65427 | LOPO-EL | < 1.34E-307 | - | -0.0022415 |
| LOPO-AL | 0.65285 | LOPO-AL | < 1.34E-307 | 1.33E-10 | - |
| LOPO-all | 0.66342 | LOPO-all | < 1.34E-307 | 4.07E-16 | 1.38E-86 |

**SIGMA_chrX**

| **Panel** | **Median** | **P-values** | LOPO-EA | LOPO-EL | LOPO-AL |
|---|---|---|---|---|---|
| LOPO-EA | 0.901175 | LOPO-EA | - | 0.0138734 | 0.0109310 |
| LOPO-EL | 0.921 | LOPO-EL | < 1.34E-307 | - | -0.0005744 |
| LOPO-AL | 0.916565 | LOPO-AL | < 1.34E-307 | 1.29E-25 | - |
| LOPO-all | 0.918015 | LOPO-all | < 1.34E-307 | 1.06E-109 | 3.16E-02 |

**Bambui_chrX**

| **Panel** | **Median** | **P-values** | LOPO-EA | LOPO-EL | LOPO-AL |
|---|---|---|---|---|---|
| LOPO-EA | 0.84794 | LOPO-EA | - | 0.0003473 | 0.0035376 |

| | | | | | |
|---|---|---|---|---|---|
| LOPO-EL | 0.84543 | LOPO-EL | 2.80E-02 | - | 0.0037551 |
| LOPO-AL | 0.85325 | LOPO-AL | 2.51E-290 | 2.53E-190 | - |
| LOPO-all | 0.85432 | LOPO-all | < 1.34E-307 | 8.50E-275 | 8.44E-02 |

**Pelotas_chrX**

| **Panel** | **Median** | **P-values** | LOPO-EA | LOPO-EL | LOPO-AL |
|---|---|---|---|---|---|
| LOPO-EA | 0.831665 | LOPO-EA | - | -0.0010915 | 0.0031275 |
| LOPO-EL | 0.825365 | LOPO-EL | 1.09E-20 | - | 0.0048130 |
| LOPO-AL | 0.835875 | LOPO-AL | 3.66E-282 | < 1.34E-307 | - |
| LOPO-all | 0.837435 | LOPO-all | < 1.34E-307 | < 1.34E-307 | 1.55E-08 |

**Salvador_chrX**

| **Panel** | **Median** | **P-values** | LOPO-EA | LOPO-EL | LOPO-AL |
|---|---|---|---|---|---|
| LOPO-EA | 0.77283 | LOPO-EA | - | -0.0143850 | 0.0056612 |
| LOPO-EL | 0.75058 | LOPO-EL | < 1.34E-307 | - | 0.0222247 |
| LOPO-AL | 0.78024 | LOPO-AL | < 1.34E-307 | < 1.34E-307 | - |
| LOPO-all | 0.77617 | LOPO-all | 6.51E-158 | < 1.34E-307 | < 1.34E-307 |

| LOPO-all | | | | | |
|---|---|---|---|---|---|
| | | | | | CUSCH-LOAI |
| LOPO-all | **Panel** | **Median** | | LOPO-EA | |
| 0.0126264 | LOPO-EA | 0.40178 | LOPO-EA | - | |
| 0.0069971 | LOPO-EL | 0.43479 | LOPO-EL | 3.97466E-29 | |
| -0.0023201 | LOPO-AL | 0.42018 | LOPO-AL | 2.91386E-45 | |
| - | LOPO-all | 0.41494 | LOPO-all | 2.6095E-104 | |

| | | | | |
|---|---|---|---|---|
| | | | | **DR_chr7** |
| LOPO-all | **Panel** | **Median** | | LOPO-EA |
| 0.0108025 | LOPO-EA | 0.39696 | LOPO-EA | - |
| 0.0078254 | LOPO-EL | 0.42247 | LOPO-EL | 3.59E-13 |
| -0.0028122 | LOPO-AL | 0.41357 | LOPO-AL | 1.85E-38 |
| - | LOPO-all | 0.40635 | LOPO-all | 9.56E-78 |

| | | | | |
|---|---|---|---|---|
| | | | | **PR_chr7** |
| LOPO-all | **Panel** | **Median** | | LOPO-EA |
| 0.0280142 | LOPO-EA | 0.467125 | LOPO-EA | - |
| -0.0012751 | LOPO-EL | 0.521615 | LOPO-EL | 3.03E-131 |
| 0.0010049 | LOPO-AL | 0.490315 | LOPO-AL | 1.67E-45 |
| - | LOPO-all | 0.49094 | LOPO-all | 2.08E-166 |

| | | | | |
|---|---|---|---|---|
| | | | | **LARGE-PD_** |
| LOPO-all | **Panel** | **Median** | | LOPO-EA |
| 0.0270989 | LOPO-EA | 0.40899 | LOPO-EA | - |
| 0.0022863 | LOPO-EL | 0.49922 | LOPO-EL | < 1.34E-307 |
| 0.0052241 | LOPO-AL | 0.46597 | LOPO-AL | < 1.34E-307 |
| - | LOPO-all | 0.46656 | LOPO-all | < 1.34E-307 |

| | | | | |
|---|---|---|---|---|
| | | | | **SIGMA_ch** |
| LOPO-all | **Panel** | **Median** | | LOPO-EA |
| 0.0120114 | LOPO-EA | 0.72788 | LOPO-EA | - |
| -0.0011117 | LOPO-EL | 0.789845 | LOPO-EL | < 1.34E-307 |
| -0.0000944 | LOPO-AL | 0.7784 | LOPO-AL | < 1.34E-307 |
| - | LOPO-all | 0.772615 | LOPO-all | < 1.34E-307 |

| | | | | |
|---|---|---|---|---|
| | | | | **Bambui_ch** |
| LOPO-all | **Panel** | **Median** | | LOPO-EA |
| 0.0038266 | LOPO-EA | 0.73235 | LOPO-EA | - |

| | | | | |
|---|---|---|---|---|
| 0.0035748 | LOPO-EL | 0.73529 | LOPO-EL | 5.68E-19 |
| 0.0001631 | LOPO-AL | 0.75538 | LOPO-AL | 9.94E-182 |
| - | LOPO-all | 0.74554 | LOPO-all | 3.27E-234 |
| | | | | **Pelotas_ch** |
| LOPO-all | **Panel** | **Median** | | LOPO-EA |
| 0.0040090 | LOPO-EA | 0.713575 | LOPO-EA | - |
| 0.0050199 | LOPO-EL | 0.71126 | LOPO-EL | > 0.99 |
| 0.0004250 | LOPO-AL | 0.735095 | LOPO-AL | 2.37E-192 |
| - | LOPO-all | 0.72482 | LOPO-all | 2.59E-296 |
| | | | | **Salvador_c** |
| LOPO-all | **Panel** | **Median** | | LOPO-EA |
| 0.0020227 | LOPO-EA | 0.68159 | LOPO-EA | - |
| 0.0175845 | LOPO-EL | 0.64805 | LOPO-EL | < 1.34E-307 |
| -0.0037340 | LOPO-AL | 0.70861 | LOPO-AL | < 1.34E-307 |
| - | LOPO-all | 0.689095 | LOPO-all | 4.28E-106 |

Comparison D

**D_chr7** **C**

| LOPO-EL | LOPO-AL | LOPO-all | Panel | Median | P-values |
|---|---|---|---|---|---|
| 0.006973814 | 0.005720161 | 0.004668136 | LOPO-EA | 0.52493 | LOPO-EA |
| - | -8.02131E-05 | -3.37084E-05 | LOPO-EL | 0.53247 | LOPO-EL |
| > 0.99 | - | -0.000111909 | LOPO-AL | 0.54554 | LOPO-AL |
| > 0.99 | > 0.99 | - | LOPO-all | 0.54112 | LOPO-all |

**7**

| LOPO-EL | LOPO-AL | LOPO-all | Panel | Median | P-values |
|---|---|---|---|---|---|
| 0.0045417 | 0.0052886 | 0.0039953 | LOPO-EA | 0.509155 | LOPO-EA |
| - | 0.0012174 | 0.0011038 | LOPO-EL | 0.513155 | LOPO-EL |
| 7.01E-02 | - | -0.0002570 | LOPO-AL | 0.52968 | LOPO-AL |
| 1.06E-01 | > 0.99 | - | LOPO-all | 0.5245 | LOPO-all |

**7**

| LOPO-EL | LOPO-AL | LOPO-all | Panel | Median | P-values |
|---|---|---|---|---|---|
| 0.0196423 | 0.0072313 | 0.0089877 | LOPO-EA | 0.632815 | LOPO-EA |
| - | -0.0098976 | -0.0069904 | LOPO-EL | 0.677035 | LOPO-EL |
| 3.73E-54 | - | 0.0020777 | LOPO-AL | 0.670635 | LOPO-AL |
| 1.87E-30 | 9.30E-09 | - | LOPO-all | 0.669235 | LOPO-all |

**_chr7**

| LOPO-EL | LOPO-AL | LOPO-all | Panel | Median | P-values |
|---|---|---|---|---|---|
| 0.0449818 | 0.0251686 | 0.0266984 | LOPO-EA | 0.622875 | LOPO-EA |
| - | -0.0152320 | -0.0120285 | LOPO-EL | 0.654865 | LOPO-EL |
| 6.53E-269 | - | 0.0031790 | LOPO-AL | 0.662855 | LOPO-AL |
| 7.13E-175 | 5.93E-31 | - | LOPO-all | 0.662795 | LOPO-all |

**ır7**

| LOPO-EL | LOPO-AL | LOPO-all | Panel | Median | P-values |
|---|---|---|---|---|---|
| 0.0336315 | 0.0238030 | 0.0220950 | LOPO-EA | 0.90075 | LOPO-EA |
| - | -0.0048914 | -0.0059966 | LOPO-EL | 0.920845 | LOPO-EL |
| 2.95E-133 | - | -0.0003982 | LOPO-AL | 0.91813 | LOPO-AL |
| 1.70E-184 | 1.32E-03 | - | LOPO-all | 0.91831 | LOPO-all |

**ır7**

| LOPO-EL | LOPO-AL | LOPO-all | Panel | Median | P-values |
|---|---|---|---|---|---|
| 0.0025742 | 0.0062829 | 0.0046052 | LOPO-EA | 0.84743 | LOPO-EA |

| | | | | | |
|---|---|---|---|---|---|
| - | 0.0035028 | 0.0026848 | LOPO-EL | 0.84568 | LOPO-EL |
| 2.28E-50 | - | -0.0004555 | LOPO-AL | 0.85438 | LOPO-AL |
| 4.62E-30 | 6.36E-03 | - | LOPO-all | 0.85433 | LOPO-all |

**r7**

| LOPO-EL | LOPO-AL | LOPO-all | **Panel** | **Median** | **P-values** |
|---|---|---|---|---|---|
| 0.0002643 | 0.0056268 | 0.0042986 | LOPO-EA | 0.83151 | LOPO-EA |
| - | 0.0057131 | 0.0046871 | LOPO-EL | 0.82442 | LOPO-EL |
| 4.39E-145 | - | -0.0001784 | LOPO-AL | 0.83746 | LOPO-AL |
| 2.02E-104 | 9.30E-01 | - | LOPO-all | 0.8374 | LOPO-all |

**hr7**

| LOPO-EL | LOPO-AL | LOPO-all | **Panel** | **Median** | **P-values** |
|---|---|---|---|---|---|
| -0.0178392 | 0.0116549 | 0.0029570 | LOPO-EA | 0.77208 | LOPO-EA |
| - | 0.0313394 | 0.0218941 | LOPO-EL | 0.7508 | LOPO-EL |
| < 1.34E-307 | - | -0.0072440 | LOPO-AL | 0.78212 | LOPO-AL |
| < 1.34E-307 | < 1.34E-307 | - | LOPO-all | 0.77709 | LOPO-all |

**USCH-LOAD_chrX**

| LOPO-EA | LOPO-EL | LOPO-AL | LOPO-all | **Panel** |
|---|---|---|---|---|
| - | 0.0066640 | 0.0165115 | 0.0131599 | LOPO-EA |
| 1.31E-106 | - | 0.0107113 | 0.0067337 | LOPO-EL |
| < 1.34E-307 | 1.57E-299 | - | -0.0036339 | LOPO-AL |
| < 1.34E-307 | 8.62E-223 | 6.60E-129 | - | LOPO-all |

**DR_chrX**

| LOPO-EA | LOPO-EL | LOPO-AL | LOPO-all | **Panel** |
|---|---|---|---|---|
| - | 0.0045046 | 0.0154273 | 0.0116076 | LOPO-EA |
| 1.34E-49 | - | 0.0119366 | 0.0075268 | LOPO-EL |
| < 1.34E-307 | < 1.34E-307 | - | -0.0039114 | LOPO-AL |
| < 1.34E-307 | 1.60E-249 | 1.14E-147 | - | LOPO-all |

**PR_chrX**

| LOPO-EA | LOPO-EL | LOPO-AL | LOPO-all | **Panel** |
|---|---|---|---|---|
| - | 0.0301841 | 0.0276493 | 0.0270041 | LOPO-EA |
| < 1.34E-307 | - | -0.0002062 | -0.0015771 | LOPO-EL |
| < 1.34E-307 | > 0.99 | - | -0.0010340 | LOPO-AL |
| < 1.34E-307 | 6.97E-10 | 1.22E-06 | - | LOPO-all |

**LARGE-PD_chrX**

| LOPO-EA | LOPO-EL | LOPO-AL | LOPO-all | **Panel** |
|---|---|---|---|---|
| - | 0.0251162 | 0.0273328 | 0.0287754 | LOPO-EA |
| < 1.34E-307 | - | 0.0025624 | 0.0026012 | LOPO-EL |
| < 1.34E-307 | 3.23E-13 | - | 0.0007793 | LOPO-AL |
| < 1.34E-307 | 5.01E-20 | 1.90E-03 | - | LOPO-all |

**SIGMA_chrX**

| LOPO-EA | LOPO-EL | LOPO-AL | LOPO-all | **Panel** |
|---|---|---|---|---|
| - | 0.0145216 | 0.0124336 | 0.0124188 | LOPO-EA |
| < 1.34E-307 | - | -0.0003895 | -0.0011501 | LOPO-EL |
| < 1.34E-307 | 1.95E-13 | - | -0.0003184 | LOPO-AL |
| < 1.34E-307 | 5.73E-113 | 1.94E-15 | - | LOPO-all |

**Bambui_chrX**

| LOPO-EA | LOPO-EL | LOPO-AL | LOPO-all | **Panel** |
|---|---|---|---|---|
| - | 0.0003473 | 0.0038285 | 0.0039866 | LOPO-EA |

| | | | | |
|---|---|---|---|---|
| 3.42E-02 | - | 0.0042284 | 0.0038465 | LOPO-EL |
| < 1.34E-307 | 1.67E-243 | - | -0.0000185 | LOPO-AL |
| < 1.34E-307 | < 1.34E-307 | > 0.99 | - | LOPO-all |

**Pelotas_chrX**

| LOPO-EA | LOPO-EL | LOPO-AL | LOPO-all | **Panel** |
|---|---|---|---|---|
| - | -0.0010122 | 0.0041595 | 0.0043446 | LOPO-EA |
| 2.52E-18 | - | 0.0058041 | 0.0053851 | LOPO-EL |
| < 1.34E-307 | < 1.34E-307 | - | 0.0000336 | LOPO-AL |
| < 1.34E-307 | < 1.34E-307 | > 0.99 | - | LOPO-all |

**Salvador_chrX**

| LOPO-EA | LOPO-EL | LOPO-AL | LOPO-all | **Panel** |
|---|---|---|---|---|
| - | -0.0134855 | 0.0066715 | 0.0026340 | LOPO-EA |
| < 1.34E-307 | - | 0.0226283 | 0.0173608 | LOPO-EL |
| < 1.34E-307 | < 1.34E-307 | - | -0.0040931 | LOPO-AL |
| 2.72E-249 | < 1.34E-307 | < 1.34E-307 | - | LOPO-all |

Comparison E

**CUSCH-LOAD_chr7**

| **Median** | **P-values** | LOPO-EA | LOPO-EL | LOPO-AL | LOPO-all |
|---|---|---|---|---|---|
| 0.40098 | LOPO-EA | - | 0.0060250 | 0.0050265 | 0.0046847 |
| 0.43313 | LOPO-EL | 1.74E-22 | - | -0.0000631 | 0.0002509 |
| 0.41938 | LOPO-AL | 8.46E-38 | > 0.99 | - | 0.0001477 |
| 0.41613 | LOPO-all | 1.26E-107 | > 0.99 | > 0.99 | - |

**DR_chr7**

| **Median** | **P-values** | LOPO-EA | LOPO-EL | LOPO-AL | LOPO-all |
|---|---|---|---|---|---|
| 0.39635 | LOPO-EA | - | 0.0035165 | 0.0045525 | 0.0038886 |
| 0.420725 | LOPO-EL | 1.35E-08 | - | 0.0014563 | 0.0015113 |
| 0.414755 | LOPO-AL | 6.59E-31 | 2.05E-03 | - | -0.0000062 |
| 0.40867 | LOPO-all | 5.91E-76 | 7.09E-03 | > 0.99 | - |

**PR_chr7**

| **Median** | **P-values** | LOPO-EA | LOPO-EL | LOPO-AL | LOPO-all |
|---|---|---|---|---|---|
| 0.46885 | LOPO-EA | - | 0.0195316 | 0.0065137 | 0.0087225 |
| 0.52364 | LOPO-EL | 8.83E-132 | - | -0.0106084 | -0.0072642 |
| 0.4879 | LOPO-AL | 6.85E-39 | 3.26E-61 | - | 0.0027269 |
| 0.48925 | LOPO-all | 6.40E-158 | 7.80E-33 | 2.07E-14 | - |

**LARGE-PD_chr7**

| **Median** | **P-values** | LOPO-EA | LOPO-EL | LOPO-AL | LOPO-all |
|---|---|---|---|---|---|
| 0.41122 | LOPO-EA | - | 0.0451994 | 0.0245280 | 0.0264393 |
| 0.50094 | LOPO-EL | < 1.34E-307 | - | -0.0154234 | -0.0125705 |
| 0.47019 | LOPO-AL | < 1.34E-307 | 1.96E-277 | - | 0.0030314 |
| 0.46844 | LOPO-all | < 1.34E-307 | 1.01E-193 | 2.61E-29 | - |

**SIGMA_chr7**

| **Median** | **P-values** | LOPO-EA | LOPO-EL | LOPO-AL | LOPO-all |
|---|---|---|---|---|---|
| 0.72903 | LOPO-EA | - | 0.0331267 | 0.0236141 | 0.0214073 |
| 0.79008 | LOPO-EL | < 1.34E-307 | - | -0.0046256 | -0.0058070 |
| 0.77774 | LOPO-AL | < 1.34E-307 | 3.82E-122 | - | -0.0006106 |
| 0.77313 | LOPO-all | < 1.34E-307 | 1.02E-183 | 2.20E-07 | - |

**Bambui_chr7**

| **Median** | **P-values** | LOPO-EA | LOPO-EL | LOPO-AL | LOPO-all |
|---|---|---|---|---|---|
| 0.73153 | LOPO-EA | - | 0.0025615 | 0.0061061 | 0.0044403 |

| | | | | | |
|---|---|---|---|---|---|
| 0.734905 | LOPO-EL | 1.83E-19 | - | 0.0036631 | 0.0025566 |
| 0.754005 | LOPO-AL | 5.21E-176 | 1.83E-53 | - | -0.0004270 |
| 0.74548 | LOPO-all | 3.14E-224 | 2.78E-27 | 1.50E-02 | - |

**Pelotas_chr7**

| **Median** | **P-values** | LOPO-EA | LOPO-EL | LOPO-AL | LOPO-all |
|---|---|---|---|---|---|
| 0.7125 | LOPO-EA | - | -0.0000085 | 0.0054914 | 0.0042860 |
| 0.70977 | LOPO-EL | > 0.99 | - | 0.0058400 | 0.0049676 |
| 0.7328 | LOPO-AL | 1.73E-188 | 4.45E-149 | - | -0.0001299 |
| 0.72404 | LOPO-all | 5.84E-296 | 3.69E-116 | > 0.99 | - |

**Salvador_chr7**

| **Median** | **P-values** | LOPO-EA | LOPO-EL | LOPO-AL | LOPO-all |
|---|---|---|---|---|---|
| 0.68132 | LOPO-EA | - | -0.0179283 | 0.0113993 | 0.0031298 |
| 0.64749 | LOPO-EL | < 1.34E-307 | - | 0.0315407 | 0.0225283 |
| 0.70776 | LOPO-AL | < 1.34E-307 | < 1.34E-307 | - | -0.0069811 |
| 0.68995 | LOPO-all | 3.80E-123 | < 1.34E-307 | < 1.34E-307 | - |

**CUSCH-LOAD_chrX**

| **Panel** | **Median** | **P-values** | LOPO-EA | LOPO-EL | LOPO-AL | LOPO-all |
|---|---|---|---|---|---|---|
| LOPO-EA | 0.52611 | LOPO-EA | - | 0.0051604 | 0.0151414 | 0.0113635 |
| LOPO-EL | 0.530255 | LOPO-EL | 1.36E-67 | - | 0.0109467 | 0.0067716 |
| LOPO-AL | 0.54219 | LOPO-AL | < 1.34E-307 | 9.10E-305 | - | -0.0037960 |
| LOPO-all | 0.53964 | LOPO-all | < 1.34E-307 | 4.98E-233 | 1.27E-144 | - |

**DR_chrX**

| **Panel** | **Median** | **P-values** | LOPO-EA | LOPO-EL | LOPO-AL | LOPO-all |
|---|---|---|---|---|---|---|
| LOPO-EA | 0.5106 | LOPO-EA | - | 0.0028441 | 0.0138654 | 0.0097161 |
| LOPO-EL | 0.51044 | LOPO-EL | 1.04E-21 | - | 0.0121456 | 0.0075537 |
| LOPO-AL | 0.52712 | LOPO-AL | < 1.34E-307 | < 1.34E-307 | - | -0.0042423 |
| LOPO-all | 0.52257 | LOPO-all | < 1.34E-307 | 1.29E-259 | 1.69E-171 | - |

**PR_chrX**

| **Panel** | **Median** | **P-values** | LOPO-EA | LOPO-EL | LOPO-AL | LOPO-all |
|---|---|---|---|---|---|---|
| LOPO-EA | 0.630015 | LOPO-EA | - | 0.0302679 | 0.0280810 | 0.0272286 |
| LOPO-EL | 0.674545 | LOPO-EL | < 1.34E-307 | - | 0.0004991 | -0.0013677 |
| LOPO-AL | 0.67275 | LOPO-AL | < 1.34E-307 | 0.7087368 | - | -0.0012245 |
| LOPO-all | 0.670925 | LOPO-all | < 1.34E-307 | 1.94702E-07 | 6.82108E-09 | - |

**LARGE-PD_chrX**

| **Panel** | **Median** | **P-values** | LOPO-EA | LOPO-EL | LOPO-AL | LOPO-all |
|---|---|---|---|---|---|---|
| LOPO-EA | 0.62576 | LOPO-EA | - | 0.0238795 | 0.0241624 | 0.0255048 |
| LOPO-EL | 0.65399 | LOPO-EL | < 1.34E-307 | - | 0.0015179 | 0.0010429 |
| LOPO-AL | 0.659705 | LOPO-AL | < 1.34E-307 | 3.23E-05 | - | 0.0007826 |
| LOPO-all | 0.662755 | LOPO-all | < 1.34E-307 | 6.05E-04 | 2.17E-03 | - |

**SIGMA_chrX**

| **Panel** | **Median** | **P-values** | LOPO-EA | LOPO-EL | LOPO-AL | LOPO-all |
|---|---|---|---|---|---|---|
| LOPO-EA | 0.90117 | LOPO-EA | - | 0.0136976 | 0.0113673 | 0.0116276 |
| LOPO-EL | 0.92092 | LOPO-EL | < 1.34E-307 | - | -0.0003660 | -0.0011601 |
| LOPO-AL | 0.91764 | LOPO-AL | < 1.34E-307 | 2.59E-12 | - | -0.0002723 |
| LOPO-all | 0.91801 | LOPO-all | < 1.34E-307 | 1.95E-113 | 5.82E-12 | - |

**Bambui_chrX**

| **Panel** | **Median** | **P-values** | LOPO-EA | LOPO-EL | LOPO-AL | LOPO-all |
|---|---|---|---|---|---|---|
| LOPO-EA | 0.848415 | LOPO-EA | - | 0.0002878 | 0.0037679 | 0.0036919 |

| | | | | | | |
|---|---|---|---|---|---|---|
| LOPO-EL | 0.844815 | LOPO-EL | 9.32E-02 | - | 0.0040503 | 0.0034595 |
| LOPO-AL | 0.85416 | LOPO-AL | < 1.34E-307 | 3.34E-222 | - | -0.0001522 |
| LOPO-all | 0.854005 | LOPO-all | < 1.34E-307 | 1.46E-260 | 1.16E-01 | - |

**Pelotas_chrX**

| **Panel** | **Median** | **P-values** | LOPO-EA | LOPO-EL | LOPO-AL | LOPO-all |
|---|---|---|---|---|---|---|
| LOPO-EA | 0.831125 | LOPO-EA | - | -0.0011575 | 0.0037896 | 0.0037941 |
| LOPO-EL | 0.82503 | LOPO-EL | 6.29E-24 | - | 0.0055569 | 0.0051277 |
| LOPO-AL | 0.836335 | LOPO-AL | < 1.34E-307 | < 1.34E-307 | - | -0.0000523 |
| LOPO-all | 0.837275 | LOPO-all | < 1.34E-307 | < 1.34E-307 | > 0.99 | - |

**Salvador_chrX**

| **Panel** | **Median** | **P-values** | LOPO-EA | LOPO-EL | LOPO-AL | LOPO-all |
|---|---|---|---|---|---|---|
| LOPO-EA | 0.77255 | LOPO-EA | - | -0.0137305 | 0.0064195 | 0.0019892 |
| LOPO-EL | 0.75119 | LOPO-EL | < 1.34E-307 | - | 0.0225769 | 0.0167913 |
| LOPO-AL | 0.782025 | LOPO-AL | < 1.34E-307 | < 1.34E-307 | - | -0.0044760 |
| LOPO-all | 0.77646 | LOPO-all | 3.66E-155 | < 1.34E-307 | < 1.34E-307 | - |

**Table S10. Pairwise comparison results for PIAA reference panels.** Media
test, using PIAA-1 as a reference. Complete pairwise comparison results can l

| | | Chromosome 7 | | | |
|---|---|---|---|---|---|
| | **Panel** | **Median** | **Mean** | **Std dev** | **Median of differences** |
| **CUSCH-LOAD** | **PIAA-1** | 0.4231 | 0.4430 | 0.3264 | - |
| | **PIAA-2** | 0.4309 | 0.4468 | 0.3258 | 0.00126 |
| | **PIAA-3** | 0.4304 | 0.4465 | 0.3260 | 0.00112 |
| | **PIAA-4** | 0.4238 | 0.4439 | 0.3263 | 0.00009 |
| | **PIAA-5** | 0.4239 | 0.4429 | 0.3267 | -0.00035 |
| | **PIAA-6** | 0.4147 | 0.4385 | 0.3277 | -0.00193 |
| | **PIAA-7** | 0.4089 | 0.4347 | 0.3284 | -0.00353 |
| | **PIAA-8** | 0.3947 | 0.4272 | 0.3308 | -0.00633 |
| | **PIAA-9** | 0.4200 | 0.4404 | 0.3281 | -0.00050 |
| **DR** | **PIAA-1** | 0.4166 | 0.4398 | 0.3257 | - |
| | **PIAA-2** | 0.4243 | 0.4437 | 0.3250 | 0.00129 |
| | **PIAA-3** | 0.4234 | 0.4429 | 0.3252 | 0.00078 |
| | **PIAA-4** | 0.4158 | 0.4395 | 0.3257 | -0.00074 |
| | **PIAA-5** | 0.4152 | 0.4384 | 0.3259 | -0.00143 |
| | **PIAA-6** | 0.4093 | 0.4341 | 0.3269 | -0.00297 |
| | **PIAA-7** | 0.4019 | 0.4307 | 0.3276 | -0.00421 |
| | **PIAA-8** | 0.3888 | 0.4236 | 0.3298 | -0.00690 |
| | **PIAA-9** | 0.4128 | 0.4367 | 0.3273 | -0.00106 |
| **PR** | **PIAA-1** | 0.4935 | 0.4811 | 0.3366 | - |
| | **PIAA-2** | 0.4975 | 0.4859 | 0.3361 | 0.00141 |
| | **PIAA-3** | 0.5031 | 0.4893 | 0.3357 | 0.00386 |
| | **PIAA-4** | 0.5095 | 0.4920 | 0.3350 | 0.00661 |
| | **PIAA-5** | 0.5094 | 0.4929 | 0.3347 | 0.00796 |
| | **PIAA-6** | 0.4928 | 0.4843 | 0.3367 | 0.00378 |
| | **PIAA-7** | 0.4797 | 0.4760 | 0.3381 | -0.00024 |
| | **PIAA-8** | 0.4662 | 0.4670 | 0.3411 | -0.00339 |
| | **PIAA-9** | 0.4965 | 0.4836 | 0.3379 | 0.00294 |
| **LARGE** | **PIAA-1** | 0.4431 | 0.4603 | 0.3211 | - |
| | **PIAA-2** | 0.4586 | 0.4695 | 0.3200 | 0.00402 |
| | **PIAA-3** | 0.4595 | 0.4706 | 0.3201 | 0.00575 |
| | **PIAA-4** | 0.4576 | 0.4698 | 0.3198 | 0.00636 |

| | | | | | |
|---|---|---|---|---|---|
| **LARGE-PD** | **PIAA-5** | 0.4596 | 0.4707 | 0.3201 | 0.00832 |
| | **PIAA-6** | 0.4597 | 0.4714 | 0.3202 | 0.01073 |
| | **PIAA-7** | 0.4581 | 0.4707 | 0.3206 | 0.01087 |
| | **PIAA-8** | 0.4530 | 0.4678 | 0.3211 | 0.00974 |
| | **PIAA-9** | 0.4656 | 0.4743 | 0.3195 | 0.01190 |
| | **PIAA-1** | 0.7461 | 0.6456 | 0.3276 | - |
| | **PIAA-2** | 0.7540 | 0.6527 | 0.3243 | 0.00220 |
| | **PIAA-3** | 0.7566 | 0.6557 | 0.3227 | 0.00396 |
| | **PIAA-4** | 0.7560 | 0.6562 | 0.3222 | 0.00476 |
| **SIGMA** | **PIAA-5** | 0.7599 | 0.6584 | 0.3212 | 0.00684 |
| | **PIAA-6** | 0.7622 | 0.6616 | 0.3195 | 0.00999 |
| | **PIAA-7** | 0.7691 | 0.6670 | 0.3174 | 0.01369 |
| | **PIAA-8** | 0.7736 | 0.6695 | 0.3170 | 0.01650 |
| | **PIAA-9** | 0.7700 | 0.6666 | 0.3178 | 0.01214 |
| | **PIAA-1** | 0.7469 | 0.6645 | 0.2993 | - |
| | **PIAA-2** | 0.7506 | 0.6681 | 0.2979 | 0.00077 |
| | **PIAA-3** | 0.7497 | 0.6675 | 0.2979 | 0.00077 |
| | **PIAA-4** | 0.7453 | 0.6644 | 0.2986 | -0.00002 |
| **Bambui** | **PIAA-5** | 0.7430 | 0.6619 | 0.2999 | -0.00062 |
| | **PIAA-6** | 0.7370 | 0.6566 | 0.3020 | -0.00245 |
| | **PIAA-7** | 0.7326 | 0.6533 | 0.3039 | -0.00404 |
| | **PIAA-8** | 0.7269 | 0.6490 | 0.3058 | -0.00641 |
| | **PIAA-9** | 0.7442 | 0.6625 | 0.3004 | -0.00039 |
| | **PIAA-1** | 0.7239 | 0.6522 | 0.2966 | - |
| | **PIAA-2** | 0.7293 | 0.6555 | 0.2956 | 0.00080 |
| | **PIAA-3** | 0.7277 | 0.6547 | 0.2957 | 0.00070 |
| | **PIAA-4** | 0.7247 | 0.6524 | 0.2962 | -0.00001 |
| **Pelotas** | **PIAA-5** | 0.7204 | 0.6496 | 0.2972 | -0.00080 |
| | **PIAA-6** | 0.7140 | 0.6440 | 0.2997 | -0.00302 |
| | **PIAA-7** | 0.7115 | 0.6409 | 0.3012 | -0.00479 |
| | **PIAA-8** | 0.7075 | 0.6369 | 0.3033 | -0.00684 |
| | **PIAA-9** | 0.7229 | 0.6499 | 0.2981 | -0.00074 |
| | **PIAA-1** | 0.6956 | 0.6299 | 0.2973 | - |
| | **PIAA-2** | 0.7003 | 0.6337 | 0.2965 | 0.00136 |
| | **PIAA-3** | 0.6978 | 0.6318 | 0.2969 | 0.00056 |
| | **PIAA-4** | 0.6923 | 0.6281 | 0.2976 | -0.00133 |

| | | | | | |
|---|---|---|---|---|---|
| **Salvador** | **PIAA-5** | 0.6866 | 0.6241 | 0.2987 | -0.00356 |
| | **PIAA-6** | 0.6778 | 0.6174 | 0.3004 | -0.00724 |
| | **PIAA-7** | 0.6731 | 0.6139 | 0.3019 | -0.00956 |
| | **PIAA-8** | 0.6708 | 0.6107 | 0.3037 | -0.01146 |
| | **PIAA-9** | 0.6893 | 0.6249 | 0.2994 | -0.00280 |

an empR2, p-values and median of differences calculated using a paired, two
be found in Table S11.

| | | | | |
|---|---|---|---|---|
| | | **Chromosome X** | | |
| **p-value** | **Panel** | **Median** | **Mean** | **Std dev** |
| - | **PIAA-1** | 0.5449 | 0.5281 | 0.2903 |
| 5.01E-34 | **PIAA-2** | 0.5513 | 0.5329 | 0.2884 |
| 2.38E-16 | **PIAA-3** | 0.5480 | 0.5311 | 0.2891 |
| > 0.99 | **PIAA-4** | 0.5406 | 0.5246 | 0.2915 |
| 0.32683 | **PIAA-5** | 0.5353 | 0.5198 | 0.2935 |
| 1.35E-23 | **PIAA-6** | 0.5324 | 0.5172 | 0.2942 |
| 2.65E-45 | **PIAA-7** | 0.5295 | 0.5143 | 0.2955 |
| 1.83E-73 | **PIAA-8** | 0.5292 | 0.5140 | 0.2955 |
| 0.025011 | **PIAA-9** | 0.5416 | 0.5248 | 0.2916 |
| - | **PIAA-1** | 0.5303 | 0.5191 | 0.2905 |
| 8.30E-32 | **PIAA-2** | 0.5358 | 0.5236 | 0.2890 |
| 4.44E-08 | **PIAA-3** | 0.5306 | 0.5205 | 0.2901 |
| 3.09E-06 | **PIAA-4** | 0.5230 | 0.5126 | 0.2931 |
| 1.94E-18 | **PIAA-5** | 0.5167 | 0.5071 | 0.2953 |
| 4.52E-48 | **PIAA-6** | 0.5142 | 0.5051 | 0.2958 |
| 1.20E-60 | **PIAA-7** | 0.5131 | 0.5039 | 0.2961 |
| 2.50E-84 | **PIAA-8** | 0.5127 | 0.5036 | 0.2960 |
| 5.72E-10 | **PIAA-9** | 0.5234 | 0.5137 | 0.2928 |
| - | **PIAA-1** | 0.6487 | 0.5941 | 0.3020 |
| 2.60E-18 | **PIAA-2** | 0.6612 | 0.6041 | 0.2975 |
| 3.54E-65 | **PIAA-3** | 0.6741 | 0.6158 | 0.2936 |
| 6.72E-139 | **PIAA-4** | 0.6825 | 0.6229 | 0.2911 |
| 4.18E-166 | **PIAA-5** | 0.6808 | 0.6231 | 0.2910 |
| 2.34E-38 | **PIAA-6** | 0.6722 | 0.6134 | 0.2942 |
| > 0.99 | **PIAA-7** | 0.6471 | 0.5904 | 0.3049 |
| 1.34E-14 | **PIAA-8** | 0.6453 | 0.5893 | 0.3052 |
| 8.89E-31 | **PIAA-9** | 0.6697 | 0.6123 | 0.2944 |
| - | **PIAA-1** | 0.6573 | 0.6122 | 0.2836 |
| 1.12E-143 | **PIAA-2** | 0.6595 | 0.6139 | 0.2830 |
| 6.78E-153 | **PIAA-3** | 0.6600 | 0.6137 | 0.2834 |
| 1.57E-175 | **PIAA-4** | 0.6562 | 0.6112 | 0.2848 |

| | | | | |
|---|---|---|---|---|
| 1.21E-247 | **PIAA-5** | 0.6560 | 0.6131 | 0.2835 |
| 1.08E-298 | **PIAA-6** | 0.6569 | 0.6142 | 0.2828 |
| 2.58E-249 | **PIAA-7** | 0.6538 | 0.6120 | 0.2837 |
| 5.92E-148 | **PIAA-8** | 0.6519 | 0.6107 | 0.2846 |
| < 8.03E-307 | **PIAA-9** | 0.6615 | 0.6170 | 0.2817 |
| - | **PIAA-1** | 0.9096 | 0.7859 | 0.2652 |
| 8.40E-184 | **PIAA-2** | 0.9111 | 0.7882 | 0.2636 |
| 3.74E-276 | **PIAA-3** | 0.9129 | 0.7902 | 0.2621 |
| < 8.03E-307 | **PIAA-4** | 0.9131 | 0.7908 | 0.2616 |
| < 8.03E-307 | **PIAA-5** | 0.9145 | 0.7932 | 0.2597 |
| < 8.03E-307 | **PIAA-6** | 0.9169 | 0.7971 | 0.2562 |
| < 8.03E-307 | **PIAA-7** | 0.9197 | 0.8005 | 0.2535 |
| < 8.03E-307 | **PIAA-8** | 0.9214 | 0.8031 | 0.2513 |
| < 8.03E-307 | **PIAA-9** | 0.9179 | 0.7976 | 0.2558 |
| - | **PIAA-1** | 0.8560 | 0.7628 | 0.2485 |
| 7.19E-25 | **PIAA-2** | 0.8570 | 0.7638 | 0.2481 |
| 1.92E-15 | **PIAA-3** | 0.8566 | 0.7632 | 0.2488 |
| > 0.99 | **PIAA-4** | 0.8549 | 0.7614 | 0.2498 |
| 1.22E-08 | **PIAA-5** | 0.8533 | 0.7601 | 0.2502 |
| 1.84E-82 | **PIAA-6** | 0.8519 | 0.7586 | 0.2508 |
| 2.92E-156 | **PIAA-7** | 0.8509 | 0.7575 | 0.2515 |
| 7.24E-223 | **PIAA-8** | 0.8508 | 0.7571 | 0.2518 |
| 0.004096775 | **PIAA-9** | 0.8559 | 0.7617 | 0.2492 |
| - | **PIAA-1** | 0.8399 | 0.7611 | 0.2345 |
| 3.15E-42 | **PIAA-2** | 0.8394 | 0.7609 | 0.2348 |
| 6.40E-18 | **PIAA-3** | 0.8386 | 0.7604 | 0.2348 |
| > 0.99 | **PIAA-4** | 0.8373 | 0.7586 | 0.2364 |
| 4.66E-21 | **PIAA-5** | 0.8358 | 0.7568 | 0.2372 |
| 1.51E-170 | **PIAA-6** | 0.8343 | 0.7552 | 0.2379 |
| 1.71E-282 | **PIAA-7** | 0.8335 | 0.7538 | 0.2389 |
| < 8.03E-307 | **PIAA-8** | 0.8329 | 0.7534 | 0.2392 |
| 1.01E-16 | **PIAA-9** | 0.8375 | 0.7585 | 0.2360 |
| - | **PIAA-1** | 0.7793 | 0.7253 | 0.2329 |
| 6.39E-68 | **PIAA-2** | 0.7801 | 0.7258 | 0.2327 |
| 4.85E-07 | **PIAA-3** | 0.7793 | 0.7251 | 0.2327 |
| 6.82E-41 | **PIAA-4** | 0.7774 | 0.7230 | 0.2339 |

| | | | | |
|---|---|---|---|---|
| 2.03E-224 | **PIAA-5** | 0.7750 | 0.7206 | 0.2352 |
| < 8.03E-307 | **PIAA-6** | 0.7726 | 0.7183 | 0.2360 |
| < 8.03E-307 | **PIAA-7** | 0.7713 | 0.7165 | 0.2370 |
| < 8.03E-307 | **PIAA-8** | 0.7714 | 0.7165 | 0.2371 |
| 1.29E-133 | **PIAA-9** | 0.7769 | 0.7222 | 0.2343 |

-sided Wilcoxon signed rank

| Median of differences | p-value |
|---|---|
| - | - |
| 0.00387 | < 8.03E-307 |
| 0.00224 | 7.22E-99 |
| -0.00294 | 4.54E-142 |
| -0.00712 | < 8.03E-307 |
| -0.00932 | < 8.03E-307 |
| -0.01190 | < 8.03E-307 |
| -0.01231 | < 8.03E-307 |
| -0.00280 | 2.41E-150 |
| - | - |
| 0.00371 | < 8.03E-307 |
| 0.00109 | 7.00E-23 |
| -0.00530 | < 8.03E-307 |
| -0.01016 | < 8.03E-307 |
| -0.01201 | < 8.03E-307 |
| -0.01303 | < 8.03E-307 |
| -0.01343 | < 8.03E-307 |
| -0.00448 | < 8.03E-307 |
| - | - |
| 0.00652 | < 8.03E-307 |
| 0.01554 | < 8.03E-307 |
| 0.02095 | < 8.03E-307 |
| 0.02107 | < 8.03E-307 |
| 0.01375 | < 8.03E-307 |
| -0.00255 | 7.29E-31 |
| -0.00324 | 1.37E-45 |
| 0.01245 | < 8.03E-307 |
| - | - |
| 0.00119 | 2.58E-13 |
| 0.00201 | 1.05E-15 |
| 0.00059 | 0.4562 |

| | |
|---|---|
| 0.00211 | 2.45E-14 |
| 0.00333 | 3.90E-29 |
| 0.00203 | 1.85E-10 |
| 0.00104 | 0.010656 |
| 0.00504 | 3.17E-86 |
| - | - |
| 0.00071 | 1.43E-111 |
| 0.00149 | 9.77E-174 |
| 0.00174 | 8.18E-202 |
| 0.00285 | < 8.03E-307 |
| 0.00491 | < 8.03E-307 |
| 0.00679 | < 8.03E-307 |
| 0.00816 | < 8.03E-307 |
| 0.00517 | < 8.03E-307 |
| - | - |
| 0.00035 | 4.00E-18 |
| 0.00017 | 0.026421 |
| -0.00055 | 1.84E-18 |
| -0.00118 | 6.68E-75 |
| -0.00193 | 9.86E-168 |
| -0.00264 | 3.21E-274 |
| -0.00294 | < 8.03E-307 |
| -0.00043 | 3.43E-15 |
| - | - |
| -0.00004 | > 0.99 |
| -0.00034 | 1.49E-09 |
| -0.00114 | 6.20E-100 |
| -0.00200 | 4.64E-266 |
| -0.00305 | < 8.03E-307 |
| -0.00402 | < 8.03E-307 |
| -0.00433 | < 8.03E-307 |
| -0.00119 | 6.40E-134 |
| - | - |
| 0.00020 | 2.84E-04 |
| -0.00024 | 0.005194 |
| -0.00158 | 5.68E-128 |

| | |
|---|---|
| -0.00319 | < 8.03E-307 |
| -0.00482 | < 8.03E-307 |
| -0.00609 | < 8.03E-307 |
| -0.00614 | < 8.03E-307 |
| -0.00197 | 3.53E-243 |

**Table S11. Full pairwise comparison results for PIAA reference panels.** Medians, [illegible]
paired, two-sided Wilcoxon signed rank test. P-values are found in the lower triangle o[illegible]
the upper triangle of the table (above the "-"). The median difference is column - row.

**Population**

| | | | | | | CUSCH-LO[illegible] |
|---|---|---|---|---|---|---|
| | **Panel** | **Median** | **p-values** | PIAA-1 | PIAA-2 | PIAA-3 |
| | **PIAA-1** | 0.42306 | PIAA-1 | - | 0.00126319 | 0.00112068 |
| | **PIAA-2** | 0.43094 | PIAA-2 | 5.01E-34 | - | -1.74E-04 |
| | **PIAA-3** | 0.43036 | PIAA-3 | 2.38E-16 | > 0.99 | - |
| | **PIAA-4** | 0.4238 | PIAA-4 | > 0.99 | 2.09E-17 | 6.10E-26 |
| | **PIAA-5** | 0.42392 | PIAA-5 | 3.27E-01 | 4.38E-26 | 1.14E-22 |
| | **PIAA-6** | 0.41466 | PIAA-6 | 1.35E-23 | 8.15E-61 | 1.51E-56 |
| | **PIAA-7** | 0.40885 | PIAA-7 | 2.65E-45 | 1.83E-86 | 4.29E-81 |
| | **PIAA-8** | 0.39471 | PIAA-8 | 1.83E-73 | 1.91E-112 | 3.16E-104 |
| | **PIAA-9** | 0.41996 | PIAA-9 | 2.50E-02 | 3.78E-27 | 9.90E-23 |

| | | | | | | DR_c[illegible] |
|---|---|---|---|---|---|---|
| | **Panel** | **Median** | | PIAA-1 | PIAA-2 | PIAA-3 |
| | **PIAA-1** | 0.416595 | PIAA-1 | - | 0.00129002 | 0.00078142 |
| | **PIAA-2** | 0.4243 | PIAA-2 | 8.30E-32 | - | -0.0005138 |
| | **PIAA-3** | 0.423415 | PIAA-3 | 4.44E-08 | 2.57E-08 | - |
| | **PIAA-4** | 0.415765 | PIAA-4 | 3.09E-06 | 1.35E-50 | 7.20E-60 |
| | **PIAA-5** | 0.415185 | PIAA-5 | 1.94E-18 | 5.19E-64 | 1.66E-46 |
| | **PIAA-6** | 0.40926 | PIAA-6 | 4.52E-48 | 7.16E-96 | 1.90E-77 |
| | **PIAA-7** | 0.401875 | PIAA-7 | 1.20E-60 | 1.76E-105 | 9.56E-89 |
| | **PIAA-8** | 0.38876 | PIAA-8 | 2.50E-84 | 5.66E-126 | 7.03E-108 |
| | **PIAA-9** | 0.412775 | PIAA-9 | 5.72E-10 | 3.93E-44 | 1.65E-25 |

| | | | | | | PR_c[illegible] |
|---|---|---|---|---|---|---|
| | **Panel** | **Median** | | PIAA-1 | PIAA-2 | PIAA-3 |
| | **PIAA-1** | 0.49347 | PIAA-1 | - | 0.00140902 | 0.00386061 |
| | **PIAA-2** | 0.49747 | PIAA-2 | 2.60E-18 | - | 0.002274 |
| | **PIAA-3** | 0.503125 | PIAA-3 | 3.54E-65 | 3.78E-48 | - |
| | **PIAA-4** | 0.50951 | PIAA-4 | 6.72E-139 | 3.00E-93 | 4.75E-52 |
| | **PIAA-5** | 0.50937 | PIAA-5 | 4.18E-166 | 4.04E-112 | 4.08E-50 |
| | **PIAA-6** | 0.49278 | PIAA-6 | 2.34E-38 | 2.78E-11 | > 0.99 |
| | **PIAA-7** | 0.479695 | PIAA-7 | > 0.99 | 1.94E-08 | 1.03E-41 |
| | **PIAA-8** | 0.46616 | PIAA-8 | 1.34E-14 | 5.86E-34 | 2.45E-68 |
| | **PIAA-9** | 0.496515 | PIAA-9 | 8.89E-31 | 1.52E-06 | 1.81E-03 |

| | | | | | | LARGE-P[illegible] |
|---|---|---|---|---|---|---|
| | **Panel** | **Median** | | PIAA-1 | PIAA-2 | PIAA-3 |
| | **PIAA-1** | 0.44309 | PIAA-1 | - | 0.00402062 | 0.00575044 |

| | | | | | |
|---|---|---|---|---|---|
| **PIAA-2** | 0.45861 | PIAA-2 | 1.12E-143 | - | 0.0014328 |
| **PIAA-3** | 0.45945 | PIAA-3 | 6.78E-153 | 4.89E-27 | - |
| **PIAA-4** | 0.45762 | PIAA-4 | 1.57E-175 | 1.27E-20 | 8.28E-07 |
| **PIAA-5** | 0.45955 | PIAA-5 | 1.21E-247 | 9.66E-63 | 4.55E-34 |
| **PIAA-6** | 0.45973 | PIAA-6 | 1.08E-298 | 4.36E-106 | 1.87E-79 |
| **PIAA-7** | 0.45812 | PIAA-7 | 2.58E-249 | 1.36E-86 | 2.63E-69 |
| **PIAA-8** | 0.45298 | PIAA-8 | 5.92E-148 | 5.44E-36 | 9.24E-27 |
| **PIAA-9** | 0.46557 | PIAA-9 | < 8.03E-307 | 1.65E-164 | 2.50E-124 |

**SIGMA**

| **Panel** | **Median** | | PIAA-1 | PIAA-2 | PIAA-3 |
|---|---|---|---|---|---|
| **PIAA-1** | 0.7461 | PIAA-1 | - | 0.00219754 | 0.00395724 |
| **PIAA-2** | 0.75403 | PIAA-2 | 8.40E-184 | - | 0.00126375 |
| **PIAA-3** | 0.75658 | PIAA-3 | 3.74E-276 | 6.77E-89 | - |
| **PIAA-4** | 0.756 | PIAA-4 | < 8.03E-307 | 1.46E-90 | 1.51E-22 |
| **PIAA-5** | 0.75986 | PIAA-5 | < 8.03E-307 | 2.74E-222 | 8.48E-100 |
| **PIAA-6** | 0.76223 | PIAA-6 | < 8.03E-307 | < 8.03E-307 | 1.54E-260 |
| **PIAA-7** | 0.7691 | PIAA-7 | < 8.03E-307 | < 8.03E-307 | < 8.03E-307 |
| **PIAA-8** | 0.77362 | PIAA-8 | < 8.03E-307 | < 8.03E-307 | < 8.03E-307 |
| **PIAA-9** | 0.77004 | PIAA-9 | < 8.03E-307 | < 8.03E-307 | < 8.03E-307 |

**Bambui**

| **Panel** | **Median** | | PIAA-1 | PIAA-2 | PIAA-3 |
|---|---|---|---|---|---|
| **PIAA-1** | 0.74688 | PIAA-1 | - | 0.00077473 | 0.00077119 |
| **PIAA-2** | 0.75059 | PIAA-2 | 7.19E-25 | - | 0.00010721 |
| **PIAA-3** | 0.74971 | PIAA-3 | 1.92E-15 | > 0.99 | - |
| **PIAA-4** | 0.74532 | PIAA-4 | > 0.99 | 4.33E-17 | 1.48E-24 |
| **PIAA-5** | 0.74297 | PIAA-5 | 1.22E-08 | 2.22E-48 | 6.62E-50 |
| **PIAA-6** | 0.73704 | PIAA-6 | 1.84E-82 | 1.40E-162 | 8.25E-172 |
| **PIAA-7** | 0.73262 | PIAA-7 | 2.92E-156 | 8.04E-244 | 3.37E-240 |
| **PIAA-8** | 0.72686 | PIAA-8 | 7.24E-223 | < 8.03E-307 | < 8.03E-307 |
| **PIAA-9** | 0.74418 | PIAA-9 | 4.10E-03 | 4.88E-33 | 6.14E-34 |

**Pelotas**

| **Panel** | **Median** | | PIAA-1 | PIAA-2 | PIAA-3 |
|---|---|---|---|---|---|
| **PIAA-1** | 0.72386 | PIAA-1 | - | 0.00079697 | 0.0006989 |
| **PIAA-2** | 0.72933 | PIAA-2 | 3.15E-42 | - | -7.94E-05 |
| **PIAA-3** | 0.72766 | PIAA-3 | 6.40E-18 | > 0.99 | - |
| **PIAA-4** | 0.72474 | PIAA-4 | > 0.99 | 4.66E-30 | 9.10E-34 |
| **PIAA-5** | 0.72035 | PIAA-5 | 4.66E-21 | 1.16E-96 | 3.53E-84 |
| **PIAA-6** | 0.71399 | PIAA-6 | 1.51E-170 | 1.73E-296 | 7.48E-281 |
| **PIAA-7** | 0.71145 | PIAA-7 | 1.71E-282 | < 8.03E-307 | < 8.03E-307 |
| **PIAA-8** | 0.70746 | PIAA-8 | < 8.03E-307 | < 8.03E-307 | < 8.03E-307 |

| | | | | | |
|---|---|---|---|---|---|
| **PIAA-9** | 0.72293 | PIAA-9 | 1.01E-16 | 2.72E-75 | 6.28E-55 |

| | | | | | **Salvador** |
|---|---|---|---|---|---|
| **Panel** | **Median** | | PIAA-1 | PIAA-2 | PIAA-3 |
| **PIAA-1** | 0.69559 | PIAA-1 | - | 0.00136498 | 0.00055727 |
| **PIAA-2** | 0.70025 | PIAA-2 | 6.39E-68 | - | -0.0008435 |
| **PIAA-3** | 0.69784 | PIAA-3 | 4.85E-07 | 1.29E-36 | - |
| **PIAA-4** | 0.69226 | PIAA-4 | 6.82E-41 | 3.26E-182 | 1.60E-130 |
| **PIAA-5** | 0.68657 | PIAA-5 | 2.03E-224 | < 8.03E-307 | 5.63E-289 |
| **PIAA-6** | 0.67779 | PIAA-6 | < 8.03E-307 | < 8.03E-307 | < 8.03E-307 |
| **PIAA-7** | 0.67313 | PIAA-7 | < 8.03E-307 | < 8.03E-307 | < 8.03E-307 |
| **PIAA-8** | 0.6708 | PIAA-8 | < 8.03E-307 | < 8.03E-307 | < 8.03E-307 |
| **PIAA-9** | 0.68928 | PIAA-9 | 1.29E-133 | 4.31E-277 | 4.17E-163 |

-values and median of differences calculated using a
f the comparison, while the median of differences are in

**AD_chr7**

| PIAA-4 | PIAA-5 | PIAA-6 | PIAA-7 | PIAA-8 | PIAA-9 |
|---|---|---|---|---|---|
| 8.79E-05 | -0.000352 | -0.0019279 | -0.0035321 | -0.0063273 | -0.0005047 |
| -0.0011548 | -0.0016411 | -0.0034762 | -0.0052726 | -0.0085338 | -0.0019398 |
| -0.0009447 | -0.0014295 | -0.0031594 | -0.0049366 | -0.008029 | -0.0016876 |
| - | -0.0003383 | -0.0017536 | -0.0034512 | -0.0062028 | -0.0004327 |
| 6.59E-03 | - | -0.0011414 | -0.002734 | -0.0055332 | -7.16E-06 |
| 1.14E-23 | 1.53E-24 | - | -0.0009717 | -0.0029653 | 0.00117178 |
| 4.48E-50 | 1.95E-44 | 1.16E-19 | - | -0.0012502 | 0.00242335 |
| 5.36E-73 | 1.98E-74 | 3.87E-39 | 9.30E-18 | - | 0.00456459 |
| 1.17E-01 | > 0.99 | 1.86E-20 | 9.63E-60 | 6.28E-79 | - |

**hr7**

| PIAA-4 | PIAA-5 | PIAA-6 | PIAA-7 | PIAA-8 | PIAA-9 |
|---|---|---|---|---|---|
| -0.0007375 | -0.0014288 | -0.0029718 | -0.0042148 | -0.0069026 | -0.0010648 |
| -0.0021202 | -0.0028131 | -0.0046118 | -0.0060291 | -0.0092657 | -0.0026129 |
| -0.0015674 | -0.0021359 | -0.0038123 | -0.0052833 | -0.0082455 | -0.0018192 |
| - | -0.0004677 | -0.0018215 | -0.0030912 | -0.0056668 | -4.96E-05 |
| 2.01E-05 | - | -0.0010128 | -0.0022578 | -0.0049539 | 0.00041521 |
| 7.85E-24 | 5.67E-19 | - | -0.0007211 | -0.0026276 | 0.00151088 |
| 2.49E-40 | 7.81E-31 | 1.21E-10 | - | -0.0011546 | 0.0025057 |
| 1.35E-62 | 9.70E-61 | 3.17E-30 | 2.52E-15 | - | 0.00455453 |
| > 0.99 | 4.40E-02 | 2.02E-32 | 1.77E-60 | 8.69E-77 | - |

**hr7**

| PIAA-4 | PIAA-5 | PIAA-6 | PIAA-7 | PIAA-8 | PIAA-9 |
|---|---|---|---|---|---|
| 0.00661079 | 0.00795988 | 0.00378091 | -0.0002446 | -0.0033883 | 0.00293756 |
| 0.00488322 | 0.00609839 | 0.00204253 | -0.001955 | -0.0055401 | 0.00132471 |
| 0.00240153 | 0.00359135 | -0.0003435 | -0.0046543 | -0.0083552 | -0.0009343 |
| - | 0.00099465 | -0.0025799 | -0.0074027 | -0.0110249 | -0.0034358 |
| 3.13E-09 | - | -0.0036364 | -0.0085324 | -0.0126372 | -0.0047151 |
| 2.14E-20 | 1.80E-71 | - | -0.0037777 | -0.00666 | -0.0006467 |
| 5.11E-92 | 5.55E-136 | 3.66E-84 | - | -0.0016945 | 0.00287702 |
| 8.10E-110 | 6.30E-153 | 1.19E-79 | 1.34E-14 | - | 0.00552219 |
| 8.53E-41 | 8.03E-82 | 8.07E-03 | 1.08E-36 | 1.53E-61 | - |

**D_chr7**

| PIAA-4 | PIAA-5 | PIAA-6 | PIAA-7 | PIAA-8 | PIAA-9 |
|---|---|---|---|---|---|
| 0.00635887 | 0.00831623 | 0.01073289 | 0.01087133 | 0.00974126 | 0.01189818 |

| | | | | | |
|---|---|---|---|---|---|
| 0.00184847 | 0.00371383 | 0.00599698 | 0.00610758 | 0.00452625 | 0.0070146 |
| 0.00069748 | 0.00242072 | 0.00471874 | 0.00507073 | 0.00369518 | 0.00571445 |
| - | 0.00201848 | 0.00411307 | 0.00439887 | 0.00317797 | 0.0050297 |
| 8.72E-46 | - | 0.00242679 | 0.00295211 | 0.00191818 | 0.00321539 |
| 2.67E-68 | 5.60E-53 | - | 0.0008286 | 0.00039965 | 0.00074479 |
| 5.47E-58 | 2.05E-36 | 5.56E-08 | - | 0.00012135 | -3.96E-05 |
| 7.97E-22 | 4.35E-10 | > 0.99 | > 0.99 | - | 0.00041312 |
| 7.46E-109 | 2.43E-59 | 8.13E-05 | > 0.99 | > 0.99 | - |

**_chr7**

| PIAA-4 | PIAA-5 | PIAA-6 | PIAA-7 | PIAA-8 | PIAA-9 |
|---|---|---|---|---|---|
| 0.00476301 | 0.00683858 | 0.00999294 | 0.0136924 | 0.01650416 | 0.01214456 |
| 0.00192303 | 0.00366746 | 0.00647069 | 0.00980602 | 0.01212816 | 0.00834915 |
| 0.00057907 | 0.00216366 | 0.0048638 | 0.00801577 | 0.01013848 | 0.00649939 |
| - | 0.00142473 | 0.00409703 | 0.00728645 | 0.00959289 | 0.00583167 |
| 1.17E-94 | - | 0.00244229 | 0.00531903 | 0.00761312 | 0.00387346 |
| 2.18E-224 | 2.88E-166 | - | 0.00272985 | 0.0047275 | 0.00110074 |
| < 8.03E-307 | 7.45E-300 | 7.67E-219 | - | 0.00190921 | -0.0013397 |
| < 8.03E-307 | < 8.03E-307 | 2.05E-200 | 2.74E-85 | - | -0.0032557 |
| < 8.03E-307 | 4.56E-256 | 5.70E-40 | 8.08E-51 | 7.29E-125 | - |

**_chr7**

| PIAA-4 | PIAA-5 | PIAA-6 | PIAA-7 | PIAA-8 | PIAA-9 |
|---|---|---|---|---|---|
| -1.63E-05 | -0.0006223 | -0.0024513 | -0.004042 | -0.0064122 | -0.0003865 |
| -0.0008134 | -0.0015598 | -0.0036954 | -0.0054111 | -0.0082912 | -0.0013608 |
| -0.0006508 | -0.0015158 | -0.0037251 | -0.0052129 | -0.0080982 | -0.0013245 |
| - | -0.0005259 | -0.0024096 | -0.0039601 | -0.0064031 | -0.0003091 |
| 1.13E-14 | - | -0.0014108 | -0.0029032 | -0.0051355 | 0.00022354 |
| 1.93E-91 | 6.39E-64 | - | -0.0008802 | -0.0025516 | 0.0017115 |
| 3.66E-160 | 3.44E-110 | 1.11E-28 | - | -0.0010567 | 0.00294168 |
| 1.78E-228 | 9.26E-176 | 4.45E-70 | 9.28E-29 | - | 0.00501516 |
| 1.97E-02 | 2.42E-01 | 2.86E-76 | 2.42E-169 | 1.63E-232 | - |

**_chr7**

| PIAA-4 | PIAA-5 | PIAA-6 | PIAA-7 | PIAA-8 | PIAA-9 |
|---|---|---|---|---|---|
| -6.24E-06 | -0.000803 | -0.0030233 | -0.0047889 | -0.0068406 | -0.0007369 |
| -0.0008697 | -0.0018193 | -0.0043743 | -0.0062595 | -0.0087481 | -0.001779 |
| -0.000652 | -0.0016247 | -0.0040657 | -0.0059464 | -0.0083628 | -0.0014107 |
| - | -0.0007721 | -0.0029608 | -0.004848 | -0.0069763 | -0.0006269 |
| 2.80E-41 | - | -0.0017093 | -0.003457 | -0.0053571 | 0.00028756 |
| 5.46E-186 | 4.15E-129 | - | -0.0011042 | -0.0025637 | 0.00204588 |
| < 8.03E-307 | 3.29E-205 | 4.07E-65 | - | -0.000988 | 0.00349283 |
| < 8.03E-307 | 4.69E-247 | 2.29E-95 | 1.64E-34 | - | 0.00539002 |

| | | | | | |
|---|---|---|---|---|---|
| 3.69E-12 | 7.10E-03 | 3.93E-159 | < 8.03E-307 | < 8.03E-307 | - |

**_chr7**

| PIAA-4 | PIAA-5 | PIAA-6 | PIAA-7 | PIAA-8 | PIAA-9 |
|---|---|---|---|---|---|
| -0.0013324 | -0.003563 | -0.0072384 | -0.0095624 | -0.0114606 | -0.0028045 |
| -0.0029023 | -0.0051059 | -0.0093842 | -0.0118548 | -0.0139487 | -0.004481 |
| -0.0017421 | -0.0039676 | -0.0079157 | -0.010319 | -0.0123297 | -0.0032031 |
| - | -0.0017575 | -0.0053786 | -0.0078194 | -0.0095866 | -0.0011972 |
| 5.59E-128 | - | -0.0029047 | -0.0051405 | -0.0066954 | 0.00073796 |
| < 8.03E-307 | 2.58E-227 | - | -0.0014678 | -0.0027496 | 0.00389305 |
| < 8.03E-307 | < 8.03E-307 | 1.80E-75 | - | -0.0006902 | 0.00581638 |
| < 8.03E-307 | 1.02E-294 | 6.68E-83 | 5.65E-12 | - | 0.00720716 |
| 2.44E-28 | 1.35E-13 | < 8.03E-307 | < 8.03E-307 | < 8.03E-307 | - |

| | | | | | CUSCH-LOAD_chrX | |
|---|---|---|---|---|---|---|
| **Panel** | **Median** | | PIAA-1 | PIAA-2 | PIAA-3 | PIAA-4 |
| **PIAA-1** | 0.544895 | PIAA-1 | - | 0.00386706 | 0.00223694 | -0.0029385 |
| **PIAA-2** | 0.551285 | PIAA-2 | < 8.03E-307 | - | -0.0014676 | -0.0069536 |
| **PIAA-3** | 0.54796 | PIAA-3 | 7.2179E-99 | 3.3077E-96 | - | -0.0053166 |
| **PIAA-4** | 0.54064 | PIAA-4 | 4.5371E-142 | < 8.03E-307 | < 8.03E-307 | - |
| **PIAA-5** | 0.53529 | PIAA-5 | < 8.03E-307 | < 8.03E-307 | < 8.03E-307 | < 8.03E-307 |
| **PIAA-6** | 0.532415 | PIAA-6 | < 8.03E-307 | < 8.03E-307 | < 8.03E-307 | < 8.03E-307 |
| **PIAA-7** | 0.529455 | PIAA-7 | < 8.03E-307 | < 8.03E-307 | < 8.03E-307 | < 8.03E-307 |
| **PIAA-8** | 0.52924 | PIAA-8 | < 8.03E-307 | < 8.03E-307 | < 8.03E-307 | < 8.03E-307 |
| **PIAA-9** | 0.541575 | PIAA-9 | 2.4077E-150 | < 8.03E-307 | < 8.03E-307 | 0.82980504 |

| | | | | | DR_chrX | |
|---|---|---|---|---|---|---|
| **Panel** | **Median** | | PIAA-1 | PIAA-2 | PIAA-3 | PIAA-4 |
| **PIAA-1** | 0.53029 | PIAA-1 | - | 0.00370965 | 0.00108964 | -0.0052986 |
| **PIAA-2** | 0.53581 | PIAA-2 | < 8.03E-307 | - | -0.0025013 | -0.0092323 |
| **PIAA-3** | 0.53062 | PIAA-3 | 7.0001E-23 | 8.882E-232 | - | -0.0064722 |
| **PIAA-4** | 0.523 | PIAA-4 | < 8.03E-307 | < 8.03E-307 | < 8.03E-307 | - |
| **PIAA-5** | 0.5167 | PIAA-5 | < 8.03E-307 | < 8.03E-307 | < 8.03E-307 | < 8.03E-307 |
| **PIAA-6** | 0.51423 | PIAA-6 | < 8.03E-307 | < 8.03E-307 | < 8.03E-307 | < 8.03E-307 |
| **PIAA-7** | 0.51309 | PIAA-7 | < 8.03E-307 | < 8.03E-307 | < 8.03E-307 | < 8.03E-307 |
| **PIAA-8** | 0.51268 | PIAA-8 | < 8.03E-307 | < 8.03E-307 | < 8.03E-307 | < 8.03E-307 |
| **PIAA-9** | 0.52341 | PIAA-9 | < 8.03E-307 | < 8.03E-307 | < 8.03E-307 | 3.6458E-19 |

| | | | | | PR_chrX | |
|---|---|---|---|---|---|---|
| **Panel** | **Median** | | PIAA-1 | PIAA-2 | PIAA-3 | PIAA-4 |
| **PIAA-1** | 0.64868 | PIAA-1 | - | 0.00652139 | 0.0155384 | 0.02094654 |
| **PIAA-2** | 0.66117 | PIAA-2 | < 8.03E-307 | - | 0.00781516 | 0.01322645 |
| **PIAA-3** | 0.67409 | PIAA-3 | < 8.03E-307 | < 8.03E-307 | - | 0.00456093 |
| **PIAA-4** | 0.68252 | PIAA-4 | < 8.03E-307 | < 8.03E-307 | 6.108E-243 | - |
| **PIAA-5** | 0.68076 | PIAA-5 | < 8.03E-307 | < 8.03E-307 | 1.214E-132 | > 0.99 |
| **PIAA-6** | 0.67221 | PIAA-6 | < 8.03E-307 | 2.177E-160 | 3.1878E-16 | 6.718E-222 |
| **PIAA-7** | 0.64712 | PIAA-7 | 7.29159E-31 | 8.949E-285 | < 8.03E-307 | < 8.03E-307 |
| **PIAA-8** | 0.64534 | PIAA-8 | 1.3666E-45 | < 8.03E-307 | < 8.03E-307 | < 8.03E-307 |
| **PIAA-9** | 0.66965 | PIAA-9 | < 8.03E-307 | 3.55E-166 | 1.4144E-45 | < 8.03E-307 |

| | | | | | LARGE-PD_chrX | |
|---|---|---|---|---|---|---|
| **Panel** | **Median** | | PIAA-1 | PIAA-2 | PIAA-3 | PIAA-4 |
| **PIAA-1** | 0.6573 | PIAA-1 | - | 0.00118904 | 0.00201368 | 0.00059265 |

| | | | | | | |
|---|---|---|---|---|---|---|
| **PIAA-2** | 0.659545 | PIAA-2 | 2.58E-13 | - | 0.00058962 | -0.000561 |
| **PIAA-3** | 0.659985 | PIAA-3 | 1.05E-15 | 8.57E-04 | - | -0.000908 |
| **PIAA-4** | 0.656235 | PIAA-4 | 4.56E-01 | 5.36E-01 | 8.23E-09 | - |
| **PIAA-5** | 0.656 | PIAA-5 | 2.45E-14 | 3.05E-04 | 2.20E-03 | 7.51E-28 |
| **PIAA-6** | 0.656945 | PIAA-6 | 3.90E-29 | 1.32E-15 | 9.47E-13 | 6.66E-39 |
| **PIAA-7** | 0.65376 | PIAA-7 | 1.85E-10 | 4.44E-03 | 2.41E-01 | 1.98E-13 |
| **PIAA-8** | 0.65188 | PIAA-8 | 1.07E-02 | > 0.99 | > 0.99 | 1.08E-04 |
| **PIAA-9** | 0.661495 | PIAA-9 | 3.17E-86 | 2.03E-63 | 1.16E-64 | 6.75E-116 |

| | | | | | **SIGMA_chrX** | |
|---|---|---|---|---|---|---|
| **Panel** | **Median** | | PIAA-1 | PIAA-2 | PIAA-3 | PIAA-4 |
| **PIAA-1** | 0.909615 | PIAA-1 | - | 0.00070523 | 0.00148907 | 0.00174427 |
| **PIAA-2** | 0.9111 | PIAA-2 | 1.43E-111 | - | 0.00067495 | 0.00087365 |
| **PIAA-3** | 0.912855 | PIAA-3 | 9.77E-174 | 1.88E-100 | - | 0.00022413 |
| **PIAA-4** | 0.9131 | PIAA-4 | 8.18E-202 | 1.53E-79 | 3.72E-21 | - |
| **PIAA-5** | 0.91449 | PIAA-5 | < 8.03E-307 | 1.34E-221 | 1.78E-118 | 7.09E-144 |
| **PIAA-6** | 0.916855 | PIAA-6 | < 8.03E-307 | < 8.03E-307 | < 8.03E-307 | < 8.03E-307 |
| **PIAA-7** | 0.919655 | PIAA-7 | < 8.03E-307 | < 8.03E-307 | < 8.03E-307 | < 8.03E-307 |
| **PIAA-8** | 0.92139 | PIAA-8 | < 8.03E-307 | < 8.03E-307 | < 8.03E-307 | < 8.03E-307 |
| **PIAA-9** | 0.917945 | PIAA-9 | < 8.03E-307 | < 8.03E-307 | < 8.03E-307 | < 8.03E-307 |

| | | | | | **Bambui_chrX** | |
|---|---|---|---|---|---|---|
| **Panel** | **Median** | | PIAA-1 | PIAA-2 | PIAA-3 | PIAA-4 |
| **PIAA-1** | 0.85601 | PIAA-1 | - | 0.00035399 | 0.00016951 | -0.0005523 |
| **PIAA-2** | 0.857005 | PIAA-2 | 4.00E-18 | - | -0.0001527 | -0.0009628 |
| **PIAA-3** | 0.85658 | PIAA-3 | 2.64E-02 | 4.99E-03 | - | -0.000744 |
| **PIAA-4** | 0.85494 | PIAA-4 | 1.84E-18 | 1.83E-56 | 1.07E-80 | - |
| **PIAA-5** | 0.85334 | PIAA-5 | 6.68E-75 | 7.47E-149 | 8.45E-127 | 9.34E-55 |
| **PIAA-6** | 0.85185 | PIAA-6 | 9.86E-168 | 1.03E-257 | 2.46E-237 | 7.59E-110 |
| **PIAA-7** | 0.850945 | PIAA-7 | 3.21E-274 | < 8.03E-307 | < 8.03E-307 | 1.25E-213 |
| **PIAA-8** | 0.850805 | PIAA-8 | < 8.03E-307 | < 8.03E-307 | < 8.03E-307 | 2.68E-254 |
| **PIAA-9** | 0.85585 | PIAA-9 | 3.43E-15 | 8.46E-55 | 5.09E-34 | > 0.99 |

| | | | | | **Pelotas_chrX** | |
|---|---|---|---|---|---|---|
| **Panel** | **Median** | | PIAA-1 | PIAA-2 | PIAA-3 | PIAA-4 |
| **PIAA-1** | 0.839885 | PIAA-1 | - | -4.31E-05 | -0.0003354 | -0.0011393 |
| **PIAA-2** | 0.83943 | PIAA-2 | > 0.99 | - | -0.0002565 | -0.0010681 |
| **PIAA-3** | 0.838565 | PIAA-3 | 1.49164E-09 | 5.1872E-12 | - | -0.0007397 |
| **PIAA-4** | 0.83726 | PIAA-4 | 6.1953E-100 | 2.0152E-94 | 1.915E-105 | - |
| **PIAA-5** | 0.83577 | PIAA-5 | 4.6395E-266 | 1.506E-265 | 1.388E-236 | 6.034E-130 |
| **PIAA-6** | 0.83432 | PIAA-6 | < 8.03E-307 | < 8.03E-307 | < 8.03E-307 | 1.058E-272 |
| **PIAA-7** | 0.833465 | PIAA-7 | < 8.03E-307 | < 8.03E-307 | < 8.03E-307 | < 8.03E-307 |
| **PIAA-8** | 0.832885 | PIAA-8 | < 8.03E-307 | < 8.03E-307 | < 8.03E-307 | < 8.03E-307 |

| | | | | | | |
|---|---|---|---|---|---|---|
| **PIAA-9** | 0.837465 | PIAA-9 | 6.3957E-134 | 4.858E-116 | 2.7469E-73 | > 0.99 |

| | | | | | **Salvador_chrX** | |
|---|---|---|---|---|---|---|
| **Panel** | **Median** | | PIAA-1 | PIAA-2 | PIAA-3 | PIAA-4 |
| **PIAA-1** | 0.77928 | PIAA-1 | - | 0.00020084 | -0.0002374 | -0.0015773 |
| **PIAA-2** | 0.78006 | PIAA-2 | 2.84E-04 | - | -0.0004217 | -0.0018264 |
| **PIAA-3** | 0.7793 | PIAA-3 | 5.19E-03 | 6.79E-24 | - | -0.0012553 |
| **PIAA-4** | 0.77739 | PIAA-4 | 5.68E-128 | 7.23E-178 | 1.25E-206 | - |
| **PIAA-5** | 0.77499 | PIAA-5 | < 8.03E-307 | < 8.03E-307 | < 8.03E-307 | 1.10E-288 |
| **PIAA-6** | 0.77264 | PIAA-6 | < 8.03E-307 | < 8.03E-307 | < 8.03E-307 | < 8.03E-307 |
| **PIAA-7** | 0.77132 | PIAA-7 | < 8.03E-307 | < 8.03E-307 | < 8.03E-307 | < 8.03E-307 |
| **PIAA-8** | 0.77144 | PIAA-8 | < 8.03E-307 | < 8.03E-307 | < 8.03E-307 | < 8.03E-307 |
| **PIAA-9** | 0.77686 | PIAA-9 | 3.53E-243 | 1.17E-304 | 5.83E-211 | 1.78E-11 |

| PIAA-5 | PIAA-6 | PIAA-7 | PIAA-8 | PIAA-9 |
|---|---|---|---|---|
| -0.00712545 | -0.0093295 | -0.01189721 | -0.0123139 | -0.0028023 |
| -0.01138159 | -0.0135816 | -0.01615449 | -0.0165567 | -0.0068953 |
| -0.00965357 | -0.0119142 | -0.01454382 | -0.0149496 | -0.0051826 |
| -0.00382993 | -0.0061011 | -0.008649504 | -0.0090637 | 0.00016488 |
| - | -0.0019429 | -0.004331668 | -0.0047785 | 0.00404376 |
| 2.8936E-194 | - | -0.002226205 | -0.0025844 | 0.0062456 |
| < 8.03E-307 | 8.116E-274 | - | -0.0003242 | 0.00883062 |
| < 8.03E-307 | 3.943E-208 | 8.05237E-11 | - | 0.00925342 |
| < 8.03E-307 | < 8.03E-307 | < 8.03E-307 | < 8.03E-307 | - |

| PIAA-5 | PIAA-6 | PIAA-7 | PIAA-8 | PIAA-9 |
|---|---|---|---|---|
| -0.0101626 | -0.0120095 | -0.01303479 | -0.0134327 | -0.0044838 |
| -0.01420233 | -0.0160788 | -0.01710819 | -0.0175074 | -0.0084143 |
| -0.01142473 | -0.0132387 | -0.01434049 | -0.014708 | -0.0056132 |
| -0.00435842 | -0.0061933 | -0.007251028 | -0.0076506 | 0.00079745 |
| - | -0.0016298 | -0.002566727 | -0.0029144 | 0.00524816 |
| 2.305E-135 | - | -0.000917721 | -0.0011812 | 0.0070503 |
| 3.4815E-148 | 3.0715E-57 | - | -0.0002762 | 0.00812448 |
| 7.8316E-169 | 2.2719E-50 | 6.64479E-09 | - | 0.0084629 |
| < 8.03E-307 | < 8.03E-307 | < 8.03E-307 | < 8.03E-307 | - |

| PIAA-5 | PIAA-6 | PIAA-7 | PIAA-8 | PIAA-9 |
|---|---|---|---|---|
| 0.02107319 | 0.01374582 | -0.002549577 | -0.0032368 | 0.01245265 |
| 0.01330562 | 0.00626729 | -0.009681498 | -0.0104442 | 0.00543882 |
| 0.004629608 | -0.0015843 | -0.01855329 | -0.0195793 | -0.0022795 |
| 3.01E-05 | -0.0063739 | -0.02425223 | -0.0253289 | -0.0071505 |
| - | -0.0062546 | -0.02407363 | -0.0250273 | -0.0070614 |
| < 8.03E-307 | - | -0.01595331 | -0.0170693 | -0.0005181 |
| < 8.03E-307 | < 8.03E-307 | - | -0.000539 | 0.01554451 |
| < 8.03E-307 | < 8.03E-307 | 6.17749E-12 | - | 0.01659173 |
| < 8.03E-307 | 0.019322 | < 8.03E-307 | < 8.03E-307 | - |

| PIAA-5 | PIAA-6 | PIAA-7 | PIAA-8 | PIAA-9 |
|---|---|---|---|---|
| 0.002114146 | 0.00333127 | 0.002028581 | 0.00104029 | 0.00503944 |

| | | | | |
|---|---|---|---|---|
| 0.0010989 | 0.00227251 | 0.001037547 | 0.00029321 | 0.00397143 |
| 0.000850976 | 0.00186623 | 0.00069153 | 6.81E-05 | 0.00366553 |
| 0.001582302 | 0.00307695 | 0.001880645 | 0.00112217 | 0.00478197 |
| - | 0.00131877 | 0.000516377 | -0.0001819 | 0.00296664 |
| 7.34E-20 | - | -0.000469357 | -0.0012859 | 0.001345 |
| 3.99E-01 | 2.95E-03 | - | -0.0004832 | 0.00226404 |
| > 0.99 | 1.94E-12 | 2.89E-07 | - | 0.00287525 |
| 2.04E-53 | 5.23E-13 | 1.38E-31 | 7.39E-47 | - |

| PIAA-5 | PIAA-6 | PIAA-7 | PIAA-8 | PIAA-9 |
|---|---|---|---|---|
| 0.002848731 | 0.00490821 | 0.006793534 | 0.00816108 | 0.00516957 |
| 0.001861319 | 0.00380236 | 0.005666254 | 0.0070982 | 0.00413291 |
| 0.0010869 | 0.00290153 | 0.004735531 | 0.00610227 | 0.00309826 |
| 0.000785149 | 0.00258836 | 0.004289182 | 0.00571376 | 0.00271051 |
| - | 0.00150854 | 0.003064035 | 0.0043992 | 0.00158771 |
| < 8.03E-307 | - | 0.001357955 | 0.00245332 | 3.88E-05 |
| < 8.03E-307 | < 8.03E-307 | - | 0.00095518 | -0.0012408 |
| < 8.03E-307 | < 8.03E-307 | 1.25E-264 | - | -0.0025356 |
| 7.18E-250 | > 0.99 | 9.82E-184 | < 8.03E-307 | - |

| PIAA-5 | PIAA-6 | PIAA-7 | PIAA-8 | PIAA-9 |
|---|---|---|---|---|
| -0.00117695 | -0.0019321 | -0.002642609 | -0.0029406 | -0.0004322 |
| -0.00163254 | -0.0023935 | -0.003090764 | -0.0034036 | -0.0008127 |
| -0.00139525 | -0.0021495 | -0.002830523 | -0.0031375 | -0.0006228 |
| -0.00051626 | -0.0012631 | -0.00197881 | -0.0022141 | 8.46E-05 |
| - | -0.0005566 | -0.001158505 | -0.0014242 | 0.00064429 |
| 3.79E-61 | - | -0.000505475 | -0.0006915 | 0.00133747 |
| 1.18E-110 | 3.68E-55 | - | -0.0001571 | 0.00197062 |
| 4.30E-143 | 1.33E-55 | 2.59E-11 | - | 0.00224684 |
| 8.09E-45 | 5.57E-157 | 1.68E-292 | < 8.03E-307 | - |

| PIAA-5 | PIAA-6 | PIAA-7 | PIAA-8 | PIAA-9 |
|---|---|---|---|---|
| -0.0020007 | -0.003051 | -0.004017816 | -0.004328 | -0.0011875 |
| -0.00193195 | -0.0029345 | -0.00389818 | -0.0042346 | -0.0010552 |
| -0.00164757 | -0.0025835 | -0.003580907 | -0.0039136 | -0.0007494 |
| -0.00074538 | -0.0017276 | -0.002619396 | -0.0029425 | -9.49E-06 |
| - | -0.0008214 | -0.001675462 | -0.0020051 | 0.00075119 |
| 1.0765E-152 | - | -0.000686706 | -0.0009737 | 0.00168271 |
| 2.0674E-274 | 7.668E-138 | - | -0.000234 | 0.00256738 |
| < 8.03E-307 | 3.55E-138 | 4.22316E-26 | - | 0.0029213 |

| | | | | |
|---|---|---|---|---|
| 5.91284E-76 | < 8.03E-307 | < 8.03E-307 | < 8.03E-307 | - |

| PIAA-5 | PIAA-6 | PIAA-7 | PIAA-8 | PIAA-9 |
|---|---|---|---|---|
| -0.00319025 | -0.0048224 | -0.00609345 | -0.0061365 | -0.0019726 |
| -0.00343996 | -0.0050692 | -0.006321593 | -0.0063742 | -0.0021828 |
| -0.00285832 | -0.0045087 | -0.005752056 | -0.0057813 | -0.0016487 |
| -0.0014187 | -0.0029451 | -0.004100207 | -0.004119 | -0.0003246 |
| - | -0.0012998 | -0.002412113 | -0.0024886 | 0.00106166 |
| 1.15E-288 | - | -0.000957178 | -0.000981 | 0.00257798 |
| < 8.03E-307 | 1.01E-179 | - | -1.94E-05 | 0.00371363 |
| < 8.03E-307 | 5.57E-117 | > 0.99 | - | 0.00376898 |
| 3.76E-102 | < 8.03E-307 | < 8.03E-307 | < 8.03E-307 | - |
